# Modeling AP2M1 Developmental and Epileptic Encephalopathy in Drosophila

**DOI:** 10.1101/2025.04.11.648441

**Authors:** Robin A. Karge, Florian P. Fischer, Hannah Schüth, Aileen Wechner, Sabrina Peter, Lukas Kilo, Mato Dichter, Aaron Voigt, Gaia Tavosanis, Karen M. J. van Loo, Henner Koch, Yvonne G. Weber, Stefan Wolking

## Abstract

Genetic defects in *AP2M1*, which encodes the μ-subunit of the adaptor protein complex 2 (AP-2) essential for clathrin-mediated endocytosis (CME), cause a rare form of developmental and epileptic encephalopathy (DEE). In this study, we modeled *AP2M1*-DEE in *Drosophila melanogaster* to gain deeper insights into the underlying disease mechanisms.

Pan-neuronal knock-down of the *Drosophila AP2M1* ortholog, *AP-2µ*, resulted in a consistent heat-sensitive paralysis phenotype and altered morphology in class IV dendritic arborization (c4da) neurons. Unexpectedly, affected flies were resistant to antiseizure medications and exhibited increased resistance to electrically induced seizures. A CRISPR-engineered fly line carrying the recurrent human disease variant p.Arg170Trp displayed a milder seizure resistance phenotype. While these findings contrast with the human phenotype, they align with previous studies on other CME-related genes in *Drosophila*. Our results suggest that hyperexcitability and seizures in *AP2M1*-DEE may stem from broader defects in neuronal development rather than direct synaptic dysfunction.

## Introduction

Epilepsies are defined by recurring epileptic seizures and rank among the most prevalent neurological disorders (Ngugi et al., 2010). Genetic causes account for at least 20% of all people with epilepsy, encompassing polygenic syndromes such as idiopathic generalized epilepsies and focal epilepsies (Collaborative, 2024; International League Against Epilepsy Consortium on Complex, 2023; May et al., 2018). Although rare, monogenic epilepsies, particularly in the form of developmental and epileptic encephalopathies (DEE), have a significant clinical and socioeconomic impact (Salcedo-Perez-Juana et al., 2023). DEEs manifest in newborns or early childhood, characterized by commonly intractable seizures, intellectual impairment, and lifelong disability (Wolking & Weber, 2015).

The µ-subunit of the adaptor protein complex 2 (AP-2), encoded by the gene *AP2M1*, is involved in clathrin-mediated endocytosis (CME) and vesicle recycling and has been identified in association with DEE (Helbig et al., 2019). It is encoded by the gene *AP2M1* and a recurrent pathogenic *de novo* variant c.508C>T (p.Arg170Trp) (GenBank: NM_004068.3) has been detected in four patients with epilepsy with myoclonic-atonic seizures, a subform of DEE. *AP2M1* is highly intolerant to genetic variation. Gene constraint analyses (Karczewski et al., 2020; Lek et al., 2016) shows a probability of loss-of-function intolerance (pLI) score of 1, a low observed/expected ratio for loss-of-function variants (o/e = 0.09), and a high missense Z-score (Z = 6.37). These findings indicate that loss-of-function variants in AP2M1 are exceedingly rare (Helbig et al., 2019). Knockout studies in mice underscored the essential role of *Ap2m1* in embryonic development. Complete gene inactivation results in embryonic lethality as early as embryonic day 3.5 (E3.5) (Mitsunari et al., 2005). However, heterozygous knockout mice display no overt phenotype, suggesting that a single functional copy of *Ap2m1* is sufficient for normal development. Conditional knockout of *Ap2m1* in neurons resulted in reduced neuronal complexity and impaired neuronal viability, emphasizing its crucial role in neuronal development and survival (Kononenko et al., 2017).

At the molecular level, *AP2M1* is a key component of clathrin-mediated endocytosis of membrane proteins (Azarnia Tehran et al., 2019). CME is a fundamental cellular process for the internalization of essential molecules, including nutrients, vesicle proteins, hormones, and receptors, through vesicle-formation at the plasma membrane. Central to this process is the Adaptor Protein Complex 2, a heterotetramer composed of the α, β2, σ2 and µ2 subunits (Mitsunari et al., 2005). AP-2 acts as a central hub, coordinating interactions between clathrin, accessory proteins, phosphoinositides, and cargo proteins to ensure precise vesicle formation and internalization (Azarnia Tehran et al., 2019; Traub, 2009; Traub & Bonifacino, 2013). The µ2 subunit plays a crucial role in transmembrane cargo recognition through the YxxΦ motif ( a tyrosine-based sorting signal) and in binding to the plasma membrane via two recognition sites for phosphatidylinositol-4,5-bisphosphate (PIP2) (Gaidarov & Keen, 1999; Jackson et al., 2010; Kovtun et al., 2020). This interaction drives a structural rearrangement of AP-2 from a ‘closed’ cytosolic state to an ‘open’ membrane-bound state, exposing the cargo-binding sites of both µ2 and σ2 (Jackson et al., 2010; Kovtun et al., 2020). Additionally, phosphorylation of Thr156 in the µ2 subunit further stabilizes the active conformation of AP-2 (Jackson et al., 2010; Ricotta et al., 2002).

The Arg170 residue is located within a basic phospholipid binding patch and is thought to play a crucial role in stabilizing membrane interactions and maintaining the protein’s open conformation (Helbig et al., 2019; Jackson et al., 2010). Structural analyses including the R170W variant result in thermodynamic instability of the AP-2 complex, particularly in its open conformation, potentially reducing cargo-binding efficiency. This was later confirmed through a transferrin uptake assay (Helbig et al., 2019), demonstrating defects in endocytosis. While the R170W variant in *AP2M1* appears to impair CME, the precise molecular mechanism linking this defect to the DEE phenotype remains unclear.

To investigate the consequences of *AP2M1* dysfunction, we used *Drosophila melanogaster* as a model organism. Notably, 81% of human epilepsy genes have orthologous genes in *Drosophila*, making it a suitable model for studying epileptic disorders (Fischer et al., 2023). Previous studies have identified CME dysfunction as a common phenotype when the functionality of individual AP-2 subunits is diminished in *Drosophila* (Choudhury et al., 2016). Flies with downregulated expression of individual AP-2 subunits, or mutations in σ2, exhibited alterations in neuromuscular-junction (NMJ) morphology - a phenotype frequently associated with CME defects (Coyle et al., 2004; Koh et al., 2004; Stevens et al., 2012; Verstreken et al., 2002). Furthermore, pan-neuronal knockdown of the σ2 subunit resulted in heat-sensitive paralysis, a hallmark of CME dysfunction (Coyle et al., 2004; Koh et al., 2004; Kosaka & Ikeda, 1983; Zinsmaier et al., 1994). Interestingly, the temperature-sensitive dynamin mutant *shibire^ts^*can suppress seizures in flies with bang-sensitive mutations (Kroll et al., 2015). Similar seizure suppression was also observed with Rab GPTase mutations, altering endocytosis and suggesting a broader mechanism by which reduced CME activity may mitigate seizure susceptibility.

Previous studies have also investigated morphological defects resulting from impaired CME. Pan-neuronal knockdown of any AP-2 subunit in *Drosophila* L3 larvae caused altered NMJ bouton morphology (Choudhury et al., 2016). Additionally, RNA interference (RNAi) mediated knockdown of the α-subunit of AP2 negatively impacted the development of class 4 dendritic arborization (c4da) neurons, likely due to CME defects (Yang et al., 2011). A similar phenotype was observed for RNAi-mediated knockdown of *nak*, as well as mutations of that gene. The *nak* gene encodes numb-associated kinase, the *Drosophila* ortholog of human *AAK1,* and interacts with *AP-2µ* (Yang et al., 2011). In humans, *AAK1* phosphorylates *AP2M1* at Thr156, significantly enhancing the cargo binding affinity of the µ2 subunit (Conner & Schmid, 2002; Ricotta et al., 2002). Reduced neuronal complexity was also seen in *shibire^ts^* larvae, when reared at restrictive temperatures, reinforcing the link between CME dysfunction and impaired neuronal development (Yang et al., 2011).

To investigate the effects of *AP2M1* dysfunction, we employed RNAi knockdown of *AP-2µ* and generated a *Drosophila* model carrying the human pathogenic p.Arg170Trp variant. We assessed the functional consequences of these manipulations through behavioral assays, imaging, and electrophysiological seizure induction to better understand how *AP2M1* dysfunction affects neuronal function in flies.

## Materials & Methods

### Fly husbandry and utilized stocks

All *Drosophila melanogaster* flies were maintained on standard cornmeal food at 25°C with a 12-hour light/dark cycle. Fly lines were obtained from the Bloomington Drosophila stock center (BDSC), from other laboratories as indicated, or were specifically created. The following fly lines were used in this study: Canton-S (BDSC: 64349), *P{w[+mW.hs]=GawB}elav[C155]* (*elav-*Gal4) (BDSC: 458), *y[1] v[1]; P{y[+t7.7] v[+t1.8]=TRiP.JF02875}attP2/ TM3, Sb[1]* (*AP-2µ-*RNAi) (BDSC: 28040), *P{y[+t7.7] v[+t1.8]=TRiP.JF02875}attP2/ TM3, P{w[+mC]=ActGFP}JMR2, Ser[1]* (*AP-2µ-*RNAi) (BDSC: 28040 with balancer from 4534), *y[1] v[1]; P{y[+t7.7] v[+t1.8]=UAS-GFP.VALIUM10}attP2* (*UAS-*GFP) (BDSC: 35786), *[LWG228] w[1118]* (WellGenetics Inc., Taiwan), *w[*];; AP-2µ[R168W] / (TM3, Sb[1])* (*AP-2µ^R168W^*) (WellGenetics Inc., with *TM6B, Tb[1]* as original balancer), *w[1118]; Df(3R)BSC685/TM6C, Sb[1] cu[1]* (*AP-2µ* Df) (BDSC: 26537), *w, eas^2f^* (*eas*) (Richard Baines, University of Manchester, UK), *UAS-dcr2/ CKG; UAS-mcD8-GFP, ppk-Gal4* (*ppk*-Gal4) (Gaia Tavosanis, RWTH Aachen University, Germany)

### Introduction of the human p.Arg170Trp variant into the fly genome via CRISPR/ Cas9

The amino acid position of the human p.Arg170Trp (R170W) variant is analogous to the position 168 in the *Drosophila AP-2μ* protein based on protein sequence alignment via Clustal Omega Multiple Sequence Alignment (EMBL-EBI). CRISPR-mediated mutagenesis was carried out by WellGenetics Inc. (Taiwan) using a modified homology-dependent repair (HDR) strategy (Kondo & Ueda, 2013). Guide RNA (gRNA) sequences targeting *AP-2µ* TCACGTACTCCAATACGTCC[AGG] and GACCCTGCGGGCTCATCAGC[AGG] were cloned into U6-promoter plasmids. A donor plasmid was constructed in the pUC57-Kan backbone and included two homology arms, the R168W codon change (CGC → TGG), and a 3xP3-DsRed marker cassette flanked by PiggyBac terminal repeats. Together with hs-Cas9 and the gRNA plasmids, the donor was injected into w[1118] embryos. HDR resulted in insertion of the DsRed marker into exon 2 of *AP-2µ*, and positive transformants were selected by eye-specific fluorescence. To remove the selection marker, flies were crossed to a source of PiggyBac transposase. Excision left behind a single TTAA motif embedded in the coding exon along with the R168W point mutation and a silent codon change at position I164 (I164I). Sequencing confirmed the incorporation of the R168W variant and successful excision of the *PBacDsRed* marker in line 220862ex2. The resulting line termed *AP-2m^R168W^* was balanced over TM6B, Tb[1] and later TM3 and Sb[1]. While the DsRed-marked allele was only viable in heterozygous flies, homozygous viability was restored after marker removal. The unaltered w[1118] strain acted as control.

### AP-2µ knockdown and quantification of knockdown efficiency via RT-qPCR

To model loss-of-function effects of *AP-2µ*, we first induced a pan-neuronal knockdown by expressing a double stranded RNA against *AP-2µ* mRNA (Transgenic RNAi Project (TRiP) line: BDSC 28040) under the control of the pan-neuronal driver *elav^C155^*. Control groups included Canton-S and flies expressing *P{y[+t7.7] v[+t1.8]=UAS-GFP.VALIUM10}attP2* with the *elav*-Gal4 driver, which is also used for the RNAi, following BDSC TRiP recommendations for RNAi experiments. Adult fly heads (3-5 days post eclosion) were collected, and total RNA was extracted using a TRIzol-based protocol. Briefly, 20 frozen fly heads per sample were homogenized in TRIzol (Invitrogen, Karlsruhe, Germany) with 1.2 – 1.4 mm ceramic beads (Mühlmeier, Bärnau, Germany) using a SpeedMill (Analytik Jena, Jena, Germany). After chloroform addition and centrifugation (12,000 g, 15 min, 4 °C), the aqueous phase was collected, RNA was precipitated with isopropanol, washed with 75% ethanol, air-dried, and resuspended in nuclease-free water (Qiagen, Hilden, Germany). RNA concentrations were measured using a NanoDrop ND-100 spectrophotometer (VWR International, Darmstadt, Germany) and sample concentrations were normalized to the lowest concentration among the samples. cDNA synthesis was performed via reverse transcription, followed by qPCR using gene-specific primers for *AP-2µ* and normalized against the housekeeping gene *RpL32*. For this, 15 µl RNA per sample were collected and an iScript cDNA Synthesis Kit (Bio-Rad Laboratories, Feldkirchen, Germany) was used to generate cDNA. To each RNA sample, 4 µl of reaction mix and 1 µl of reverse transcriptase enzyme were added, followed by the addition of nuclease-free water to adjust the total reaction volume to 20 µl. The cDNA synthesis reaction was performed using a T-Professional Basic thermocycler (Biometra, Göttingen, Germany) with the following conditions: priming at 25°C for 5 minutes, reverse transcription at 46°C for 20 minutes, and a final enzyme inactivation step at 95°C for 1 minute. For qPCR, the generated cDNA was quantified using a QuantStudio I apparatus (Applied Biosystems, Foster City, CA, USA). Samples were prepared with 1x iQ SYBR Green supermix (Bio-Rad Laboratories, Feldkirchen, Germany), 5 pmol of each oligonucleotide primer and 2.5-fold diluted synthesized cDNA for a total volume of 20 µl and pipetted as triplicates on a 384-well plate (Bio-Rad Laboratories, Feldkirchen, Germany). The samples were then run for 3 min at 95°C, 40 cycles of 15s at 95°C and 1 min at 60°C. For *AP-2µ*, the forward primer 5ʹ-TCT TCC ACA TCA AGA GAG CAA A-3’ and reverse primer 5’-GCC GAA GTA GGA TTG CAT CAC-3’ were used, while for *RpL32*, the forward primer 5’-TGC TAA GCT GTC GCA CAA ATG-3’ and reverse primer 5’-ATC CGT AAC CGA TGT TGG GC-3’ were used. Gene expression levels were quantified using the ΔΔCt method (Livak & Schmittgen, 2001), comparing *AP-2µ* transcript levels in *elav-AP-2µ*-RNAi flies to the controls. Off-target effects are not reported for the utilized RNAi line based on UP-TORR (Hu et al., 2013), but remain a possibility. Statistical significance was assessed using an unpaired t-test.

### Drug feeding and experimental preparation

For behavioral testing, adult flies aged 1-3 days post eclosion were collected into food vials using CO₂ anesthesia (max. 15 animals per vial). The vials were prepared with 100 µl of dissolved anti-seizure medications (ASMs), which were pipetted onto the food and left to completely dry. The following concentrations and compounds were used: 0.3 mM valproate (VPA) (Sigma-Aldrich, Steinheim, Germany) in water, 3 mM levetiracetam (LEV) (Sigma-Aldrich, Steinheim, Germany) in water, 3 mM phenytoin (PHT) (Sigma-Aldrich, Steinheim, Germany) in ethanol, based on previous publications (Fischer et al., 2024; Pandey & Nichols, 2011). Solvents were used as controls. Flies were left on the prepared food for 2 days and used for experiments 3-5 days post eclosion. To ensure comparability, flies in all experiments received solvent (water) treated food.

### Behavioral assays

For behavioral testing, both vortex assay and heat assays were employed. Flies were collected into empty vials using CO₂ anesthesia with 5 animals per vial and left to recover for a minimum of 1 hour. For the vortex assay, flies were subjected to mechanical stress using a vortex mixer (Vortex Genie 2m, Scientific Industries) at maximum speed for 10 seconds (Fischer et al., 2024; Kuebler & Tanouye, 2000; Mituzaite et al., 2021). Seizure-like behavior was recorded via a video camera, and seizure probability (ratio of seizing to non-seizing flies) was determined. For the heat assay, vials containing flies were placed in a water bath at 40-41°C for 120 seconds. Fly behavior was recorded with a video camera and later analyzed in 5 second intervals. To assess paralysis, all flies were counted as paralyzed which lost posture or showed wing buzzing. The air temperature in the vials reached 35°C after approximately 45 seconds and 39°C after approximately 90 seconds.

### Electrophysiology of the Drosophila nervous system

To determine seizure susceptibility of the flies and functionality of the giant fiber system (GFS), *in vivo* electrophysiological recordings at the dorsal longitudinal flight muscle (DLM) were performed (Allen & Godenschwege, 2010; Kuebler & Tanouye, 2000; Lee & Wu, 2002). Flies were mounted on non-poisonous glue traps (Gelbtafeln) (Inseko, Hohenwarsleben, Germany) for recordings. A tungsten electrode serving as ground was inserted in the abdomen of the fly and a glass electrode of ∼10 MΩ resistance filled with saline (Allen & Godenschwege, 2010) was inserted in the DLM 45a (Allen & Godenschwege, 2010; Gu & O’Dowd, 2006). Two tungsten electrodes were inserted in the brain for stimulation. Signals from the DLM were recorded using an EXT-10-2F extracellular amplifier, housed in an EPMS-07 (npi Electronic GmbH, Tamm, Germany). Signals were digitalized via a CW-INT-20USB interface (npi Electronic GmbH, Tamm, Germany) and recorded in WinEDR software (University of Strathclyde, Glasgow, UK). Analysis was performed in WinEDR or MatLab using a custom script (The Mathworks Inc., Natick, Massachusetts). An ISO-01M-100 stimulus isolator (npi Electronic GmbH, Tamm, Germany) was used to deliver electric stimuli to the brain. Activation of the GFS was achieved via monophasic 10 V stimuli with 0.1 ms duration. Short single pulses with 0.1 ms duration and 10 V amplitude were used to activate the GFS, leading to a voltage response at the DLM after approximately 1.4 ms. Seizures were induced using a high-frequency stimulus (HFS) of varying voltage with 0.1 ms duration, 200 Hz frequency and 2000 ms total stimulus length. Two types of seizure induction protocols were used. The first protocol aimed to determine the seizure threshold of the tested flies starting at 5 V HFS with increments of 5 V every 5 minutes until 30 V. The second protocol started at 20 V with 5 V increments every 5 minutes until 30 V. During both protocols, giant fiber functionality was monitored via a 0.1 ms pulse with 10 V amplitude every 2 seconds. Successful seizure induction manifested in the DLM as characteristic seizure-like activity (Lee & Wu, 2002).

Metal electrodes were fabricated from 0.1 mm tungsten wire (Bowen, 2010; Hurkey et al., 2023), which was electrochemically sharpened in a 1 M KOH solution. The tungsten wire was attached to the anode of the stimulus isolator, while a platinum wire was attached to the cathode. The platinum wire was placed inside the KOH solution and a monophasic current with 100 Hz, 40 V and 1 ms duration was applied. The tungsten wire was repeatedly lowered into the solution until a conic shaped tip formed. Afterwards it was rinsed with deionized water.

### Imaging of c4da-neurons in L3 larvae

To evaluate the morphology of c4da-neurons in flies with *AP-2µ* dysfunction, the gene was knocked-down via RNAi specifically in c4da-neurons using a *ppk*-Gal4 driver line, with knockdown strength enhanced by co-expression of *dicer2* (Kim et al., 2006). Images were acquired at a Zeiss LSM900 confocal microscope (Carl Zeiss AG, Oberkochen, Germany). Larvae in the L3 stadium were collected and washed. Only larvae showing GFP expression exclusively in c4da-neurons were selected for imaging, as GFP in other tissues indicated the presence of a balancer chromosome. A single larva was then placed onto a microscope slide with a drop of halocarbon oil 27. A glass coverslip was used to flatten out the larva and fixed with double-sided tape. Neurons of the abdominal hemi-segment 4 were imaged and reconstructed using arivis Vision4D software (Carl Zeiss AG, Oberkochen, Germany). In the reconstructed neurons, the number of branch points, terminals and total dendritic length were analyzed and compared. Branch points were defined as sections in the dendrite where a single process bifurcates into two or more branches, while terminals were identified as the endpoints of dendritic branches, representing the final points of neuronal outgrowth. The spatial extent of the neurons was quantified using a convex hull analysis, which assesses the outer boundary enclosing all dendritic branches. Following 3D reconstruction, a pipeline-generated object of the dendritic arbor was used to compute the minimal convex volume enclosing all branches, using a custom script provided by arivis.

### Imaging of neuromuscular junction in L3 larvae

The neuromuscular junction of L3 larvae was imaged using an Echo Revolution automated hybrid widefield microscope (Echo Inc., San Diego, CA, USA). The larvae were dissected in cold phosphate-buffered saline (PBS) and the fillet stretched out using pin needles (Brent et al., 2009). Larvae were then fixated for 20 minutes in 4% paraformaldehyde and washed with PBS containing 0.3% Triton X-100 (PBS-T). Then, the samples were blocked with 10% goat serum in PBS blocking solution. Primary antibody staining was performed with mouse anti-CSP (1:50) (DCSP-3 (1G12), Developmental Studies Hybridoma Bank, IA, USA) and rabbit anti-HRP (1:100) (500-8634-HRP, AbboMax, CA, USA) antibodies in blocking solution over night at 4°C. Samples were washed again with PBS-T and secondary antibody incubation was conducted with goat anti-mouse AF555 and goat anti-rabbit AF488 (1:750) (Invitrogen, Thermo Fisher Scientific, Waltham, MA, USA) antibodies for 3 – 5 hours at room temperature (RT). Samples were then mounted in Vectashield antifade mounting medium without DAPI (H-1000-10, Vector Laboratories, NY, USA) and sealed with clear nail polish. Imaging was accomplished with a 20x and 40x air objective and Echo’s integrated software.

## Results

### AP2M1 is highly conserved and the human pathogenic p.Arg170Trp variant disturbs a conserved phospholipid-binding patch

*AP2M1* is evolutionary highly conserved with a DIOPT score of 14/15 (DRSC (Drosophila RNAi Screening Center) Integrative Ortholog Prediction Tool) (Hu et al., 2011). Alignment of the *Drosophila AP2-2µ* sequence with *C. elegans*, zebrafish, mouse and human showed a high degree of overlap (Fig. 1 A, B). 85.23% of the protein sequence of *AP-2µ* in *Drosophila* aligns with the human AP2M1 protein sequence. The pathogenic variant R170W is located in the N-terminal region of the C-terminal mu homology domain (PF00928) (Fig. 1 C, D), which harbors the cargo-binding YxxΦ motif. The region neighboring R170 is largely identical across species and enriched with basic, positively charged amino acids (Fig. 1 A). Notably, R170 is part of a conserved basic patch (residues Lys167, Arg169, Arg170, Lys421) that contributes to electrostatic interactions with negatively charged membrane components, such as PIP₂, and is implicated in AP2 membrane recruitment and cargo recognition (Helbig et al., 2019; Jackson et al., 2010). Replacement of arginine at the R170 residue with a large, hydrophobic amino acid such as tryptophan (Fig. 1 E, F) might therefore negatively influence cargo binding capabilities of the AP2 complex (Helbig et al., 2019; Kovtun et al., 2020).

**Figure 1:**
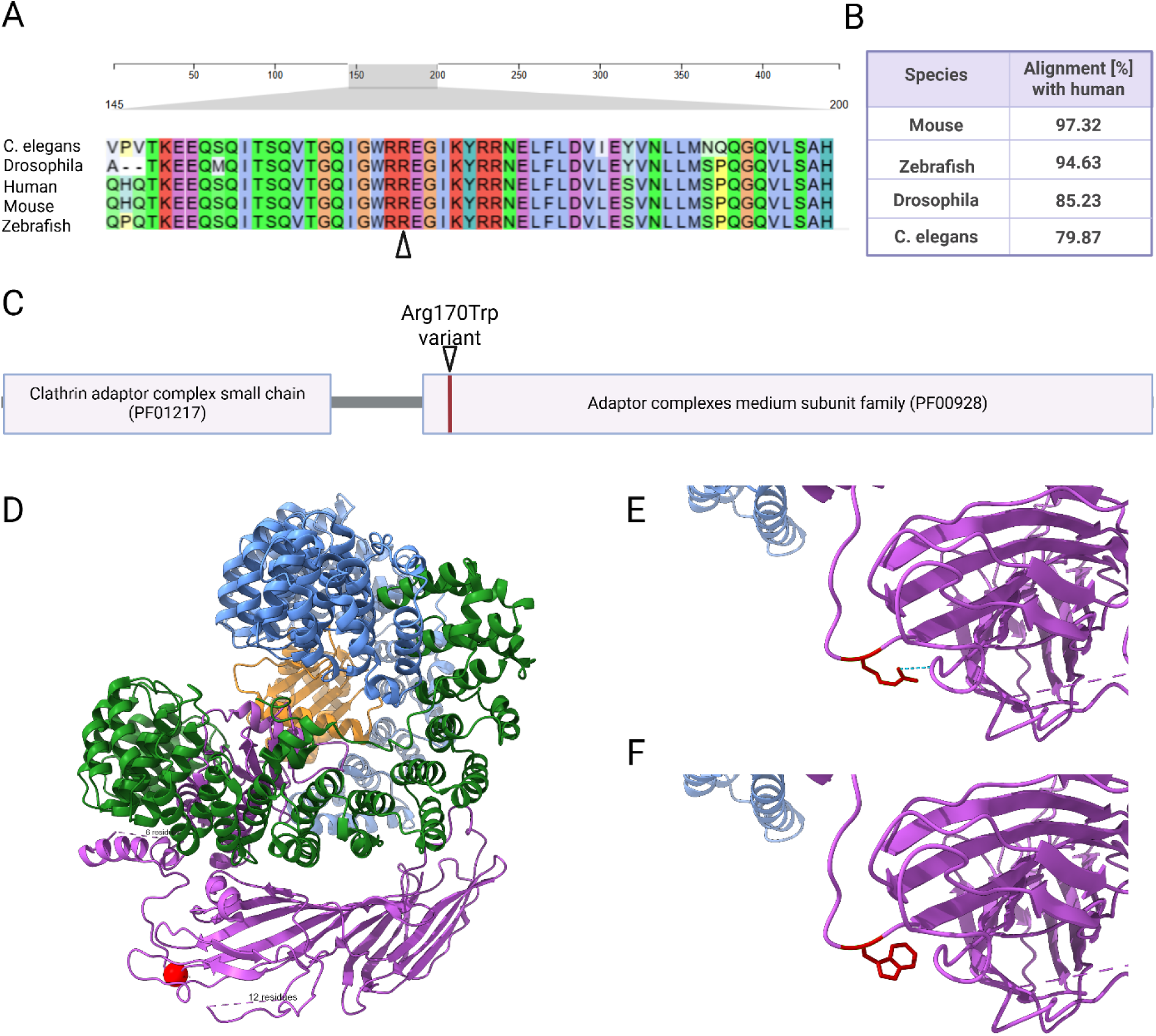
Evolutionary conservation and protein domain structure of AP2M1. **(A)** Clustal Omega Alignment (Madeira et al., 2024) of *AP2M1* segment bordering the R170 residue with orthologues genes from different species with Clustal2 color scheme. Red marks positively charged amino acids. The arrowhead marks the R170 position. **(B)** Pairwise alignment percentages of the AP2M1 protein between humans and other species. **(C)** Protein domains of AP2M1. The two major functional domains, the Clathrin adaptor complex small chain (PF01217) and Adaptor complex medium subunit family (PF00928), are highlighted. The R170W variant is located in the N-terminal region of the mu homology domain, where it is part of a basic phospholipid binding patch. **(D**) Modeling of AP-2 in the membrane-bound form with ChimeraX 1.9 (Meng et al., 2023) based on Protein Data Bank (PDB): 6YAH (Kovtun et al., 2020). The R170W variant is located near the linker region of the µ2-subunit and depicted as a red sphere. AP-2 subunits: AP-2α (blue), AP-2β (green), AP-2µ (magenta), AP-2σ (orange). **(E)** In the wildtype structure, the polar arginine residue at position 170 forms a hydrogen bond with the side chain of aspartate at position 428. **(F)** In the R170W variant, the polar arginine is replaced by a nonpolar tryptophan.

### Modeling of AP2M1 dysfunction in Drosophila melanogaster

To model *AP2M1* dysfunction in *Drosophila*, we used two different approaches. First, we induced a pan-neuronal knock-down of *AP-2µ,* the *Drosophila* homologue of *AP2M1* (Fig. 2 A). Using the pan-neuronal *elav^C155^*-Gal4 driver line, we overexpressed a construct that produces double-stranded RNA targeting *AP-2µ* (Perkins et al., 2015) and inducing its RNAi-mediated knockdown. Expression of GFP via the *elav^C155^*-Gal4 driver line was utilized as control (Fig. 2 B). The knockdown efficiency was confirmed by RT-qPCR of *AP-2µ* in adult fly heads (Fig. 2 C). Expression of *AP-2µ* in the knockdown was reduced by approximately 74% in male flies and 93% in female flies. As a second strategy to model AP2M1 dysfunction in *Drosophila*, we introduced the R168W mutation (homologous to human R170W) into *AP-2µ* (Fig. 2 D). Sanger sequencing confirmed successful genome editing at the target site, yielding homozygous viable mutant flies (Fig. 2 E). Both the knockdown and humanized lines lacked spontaneous behavioral or morphological abnormalities and displayed a normal development.

**Figure 2:**
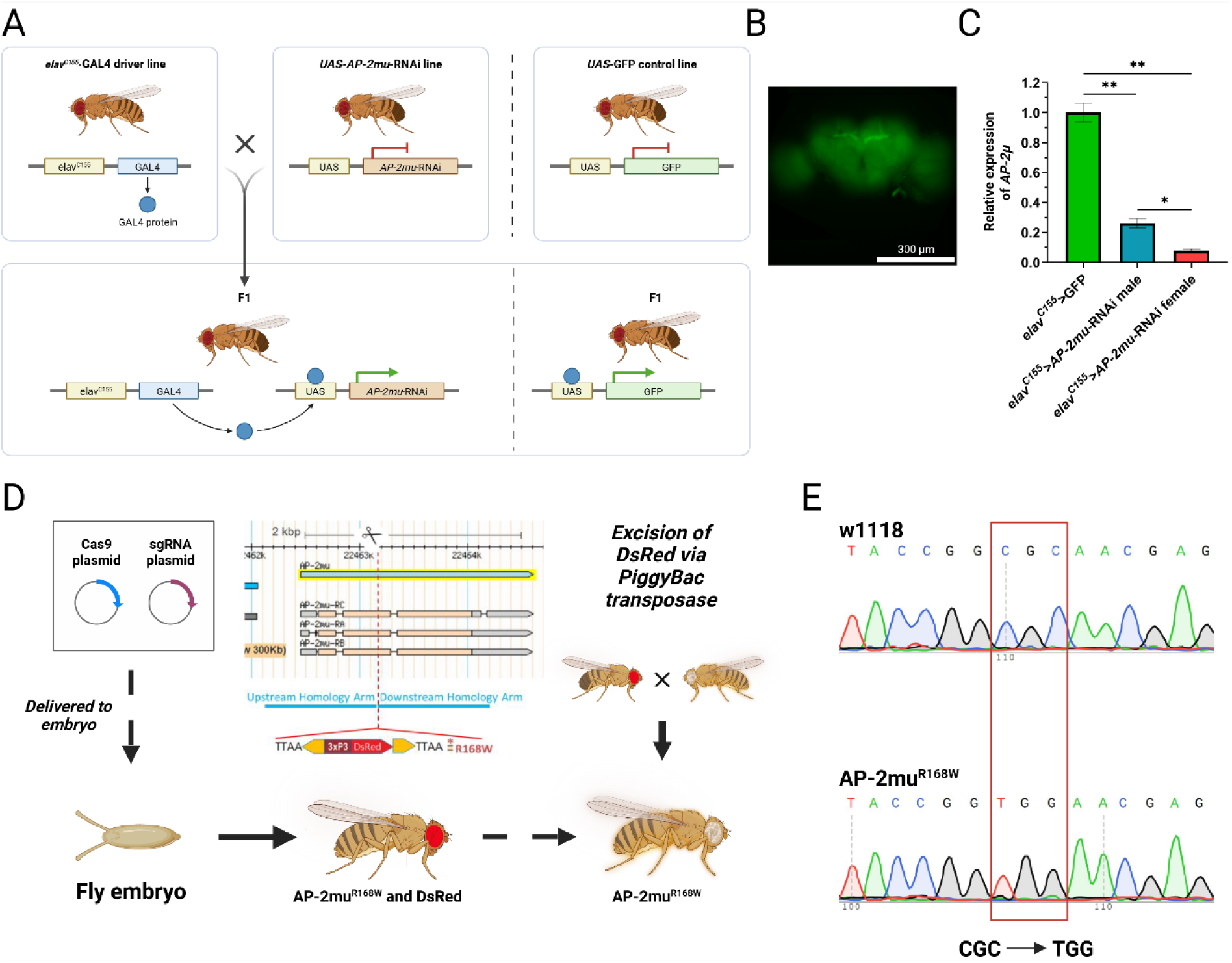
Genetic approaches to model *AP-2µ* dysfunction. **(A)** Crossing scheme to induce RNAi knockdown of *AP-2µ* in the first generation of offspring with GFP expression as control. **(B)** Expression of UAS-GFP in a male fly brain under control of *elav^C155^*-Gal4 (Scalebar = 300 µm) **(C)** Validation of *AP-2µ* knockdown via RT-qPCR. Expression levels of *AP-2µ* relative to *RpL32* shows a significant reduction in RNAi-targeted flies compared to controls. Nested t-test of 6 samples from 2 biological replicates of 20 fly heads each (*p < 0.05, **p < 0.01). **(D)** Schematic of the CRISPR-mediated mutagenesis workflow used to generate the *AP-2µ^R168W^* knock-in allele. The amino acid substitution R168W (homologous to human R170W) was introduced into the *Drosophila AP-2µ* locus using homology-directed repair (HDR). **(E)** Sanger sequencing chromatograms of the *AP-2µ* locus in background (w1118) and AP-2µ^R168W^ flies. The red box highlights the nucleotide triplet coding for the amino acid at position 168, corresponding to a change from arginine to tryptophane in the protein.

### Pan-neuronal knockdown of AP-2µ results in heat-sensitive paralysis

To start testing whether impairment of *AP-2µ* function caused seizure-like behavior in flies, we used two established assays (Mituzaite et al., 2021). The vortex assay is used to test bang-sensitivity (Fischer et al., 2024; Mituzaite et al., 2021; Parker et al., 2011), while the heat assay exploits the fact that elevated temperatures can trigger seizures in certain epilepsy models (Burg & Wu, 2012; Mituzaite et al., 2021). *AP-2µ* knockdown and *AP-2µ^R168W^*flies were tested, as well as the respective controls and bang-sensitive *eas^2f^* flies. While *eas^2f^* mutants that served as positive controls (Kroll & Tanouye, 2013) displayed clear bang sensitivity in the vortex assay, neither pan-neuronal knock-down of *AP-2µ*, nor the *AP-2µ^R168W^* mutation yielded a bang-sensitive phenotype (Fig. 3 A). In the heat assay, *AP-2µ* knockdown flies exhibited a pronounced paralysis phenotype (Fig. 3 B). After two minutes of heat exposure, 48.62% ± 5.51% of the male and 24.56% ± 3.96% of the female flies displayed paralysis, defined as a loss of posture with or without wing-buzzing. During the assay, the flies exhibited fluctuating behavioral states, shifting between paralysis and an upright position, with some flies remaining unaffected. Based on the observed behavior, we hypothesized that the phenotype might reflect seizure-like activity, prompting us to test whether ASMs could alleviate it (Fischer et al., 2024). We applied 100 µl of dissolved ASMs onto the fly food (Fischer et al., 2024) and exposed flies to the food for 2 days. We used the common ASMs valproate (VPA), levetiracetam (LEV) and phenytoin (PHT) to treat male knockdown flies, with the solvents itself as well as DMSO as control. Drug consumption was initially verified via food coloring in previous experiments (Fischer et al., 2024). However, no significant change in the heat-sensitive phenotype was observed, with 50.33% to 62.66% of the flies remaining paralyzed by the end of the assay (Fig. 3 D). *AP-2µ^R168W^* flies did not exhibit either bang sensitivity or heat sensitivity. To evaluate a potential effect of the rearing temperature on the phenotype as seen in other mutants (Garber et al., 2012), *AP-2µ^R168W^* flies were furthermore raised at 18°C, which did not lead to a heat-sensitive phenotype either. To assess whether the variant caused haploinsufficiency, we crossed the allele over a deficiency for *AP-2µ* (BDSC: 26537) (Cook et al., 2012), resulting in a fly line with only one functional copy of *AP-2µ* carrying the R168W variant. This cross did not lead to a heat sensitive phenotype either (data not shown).

**Figure 3:**
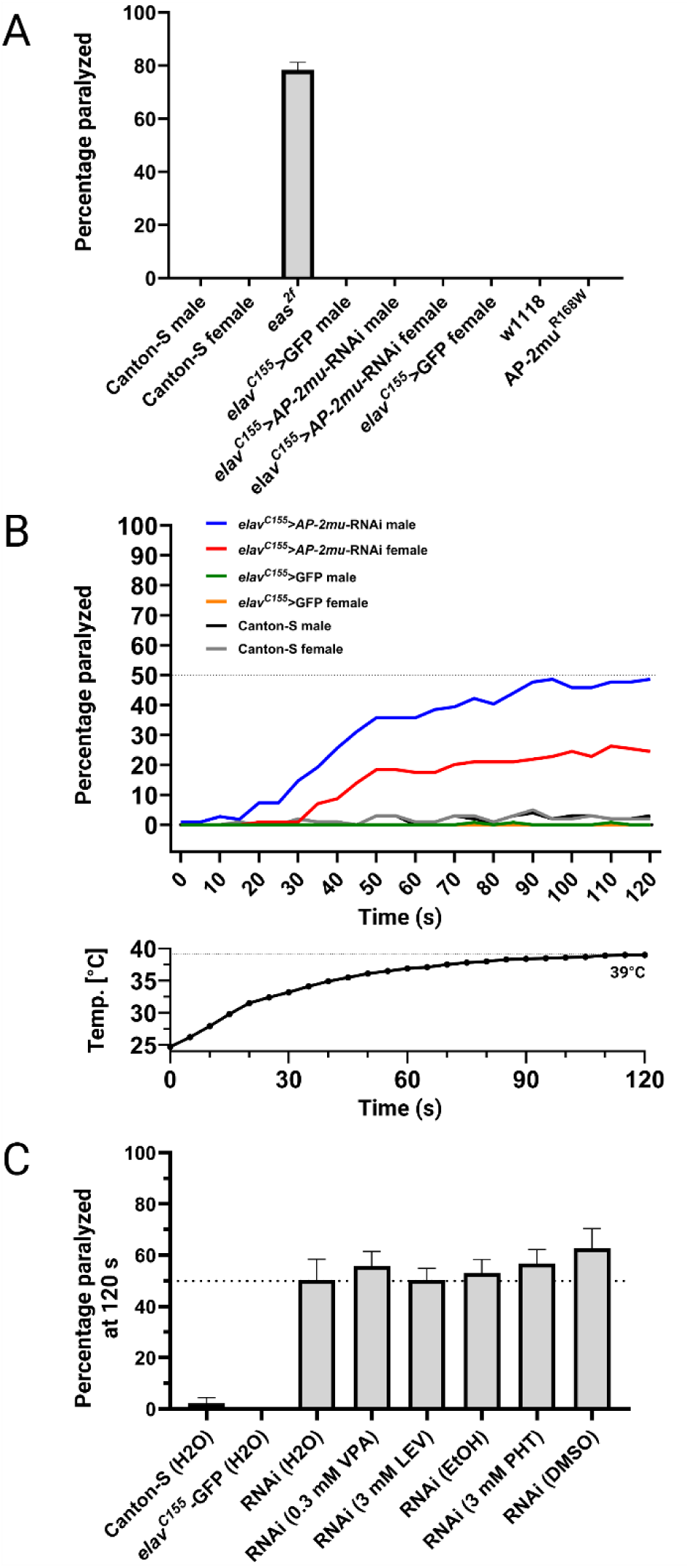
Assessment of behavioral phenotypes using vortex and heat assay. **(A)** Behavioral response to mechanical stimuli in the vortex assay, in which only bang-sensitive *eas^2f^* mutant flies showed paralysis. (n>25 flies per condition) **(B)** Heat induced paralysis of *AP-2µ* knockdown flies in the water bath assay. After 2 minutes in 40°C water, *elav^C155^-AP-2µ*-RNAi flies exhibit significantly more paralysis than *elav^C155^-*GFP controls. Male flies were affected more strongly and nearly 50% paralyzed at the end of the trial, while females exhibited a lower degree of paralysis. (n>100 flies per condition, n=36 for *elav^C155^>GFP* female). *eas^2f^*, w1118 and *AP-2µ^R168W^* male flies are not shown as they exhibited no heat sensitivity. Air temperature in vials during the heat assay is shown below. The temperature rises steadily from ∼24 °C in the beginning to 39 °C in the end. **(C)** Percentage of flies paralyzed at the end of the heat assay after drug-feeding for 2 days. Treatment of male knockdown flies with anti-seizure medication (ASM) did not alleviate the paralysis (n > 75 flies per treatment condition) (VPA = valproate; LEV = levetiracetam; PHT = phenytoin).

### Knock-down and AP-2µ^R168W^ flies exhibit increased resistance to electrically induced seizures

To assess the impact of *AP-2µ* knock-down or the R168W variant on neuronal conductivity, we performed electrophysiological recordings of the GFS at the DLM (Allen & Godenschwege, 2010) (Fig. 4 A). When providing short, single pulses to the brain, no differences were observed in the GFS response to individual stimuli between the tested genotypes (Fig. 4 B). To induce seizures, HFS were delivered to the brain at different voltage levels (Kuebler & Tanouye, 2000; Lee & Wu, 2002; Saras et al., 2017). Successful seizure induction was characterized by aberrant high-frequency firing at the DLM, with distinct phases, including an initial seizure, synaptic failure, and a recovery seizure (Fig. 4 C). Canton-S flies were used to establish the seizure induction protocol, starting with a 5 V stimulus and increasing in 5 V increments every 5 minutes up to 30 V. In Canton-S flies, seizure induction succeeded consistently with an average seizure threshold of 11.54 V ± 1.04 V. For *eas^2f^* flies, the protocol was adapted to start at 1 V with incremental increases of 1 V to address their reduced seizure threshold. Seizures could be induced in all *eas^2f^* flies, with an average seizure threshold of 3.11 V ± 0.20 V (Table 1). When applying the 5 V step protocol in *AP-2µ* knockdown and *AP-2µ*^R168W^ flies, we found that only a fraction of these flies responded to seizure induction (Fig. 4 D). *AP-2µ* knockdown flies exhibited a significantly lower percentage of successful seizure inductions compared to the GFP expressing controls (Table 1). Also, *AP-2µ*^R168W^ flies displayed significantly less seizure-like activity compared to controls (Fig. 4 D) (Table 1). In cases where a seizure was induced in knockdown or variant flies, the seizure thresholds were similar to those of their respective controls (Table 1), suggesting that other factors may contribute to the reduced likelihood of seizure occurrence. We next tested an alternative seizure induction protocol, starting at 20 V, to determine whether the incremental voltage increases in the primary protocol might be hampering seizure susceptibility. In the alternative protocol, we found no significant differences in the number of seizing flies compared to the controls (Fig. 4 E). This observation suggests a seizure-protecting effect of sub-threshold stimulation in flies with *AP-2µ* dysfunction.

**Figure 4:**
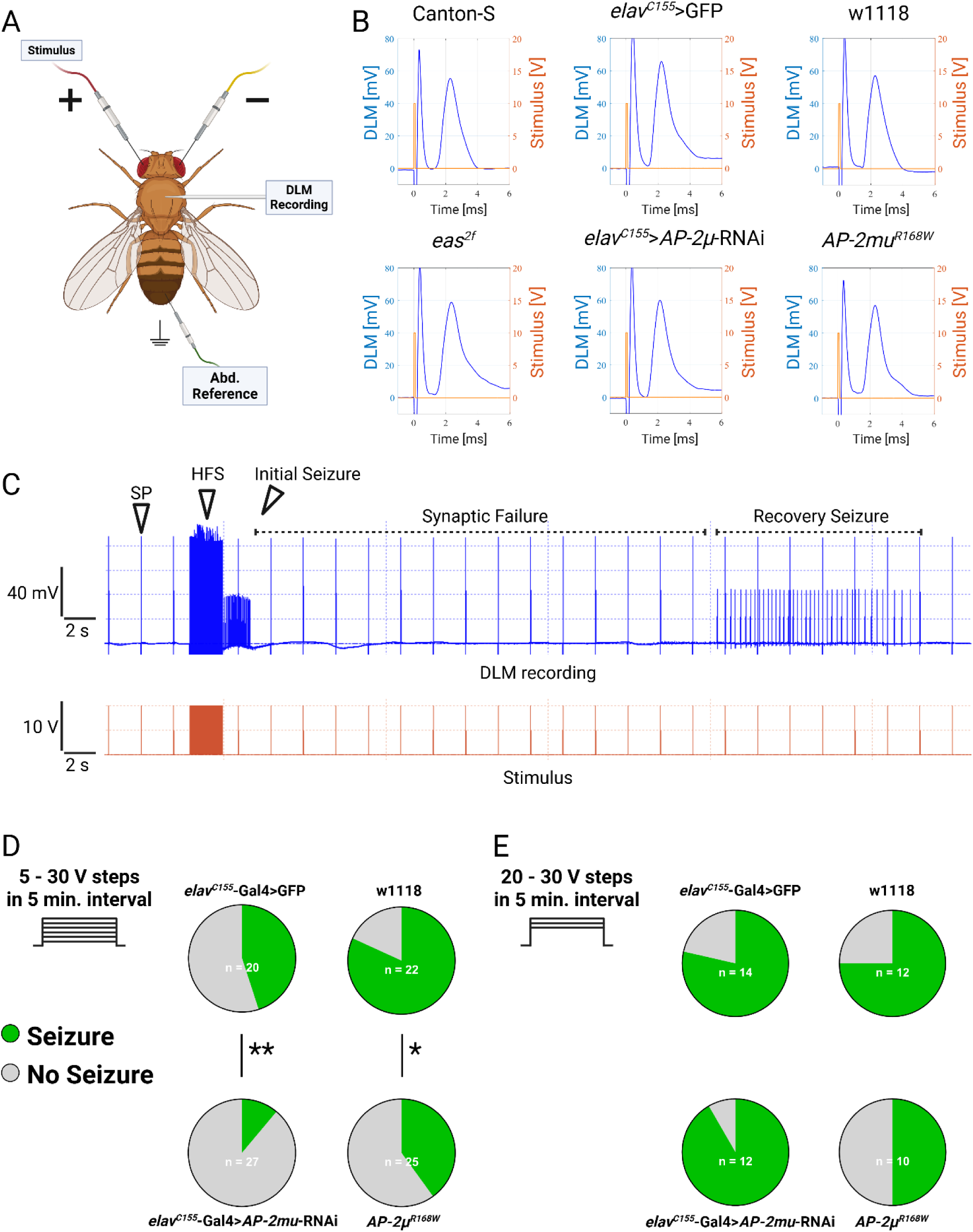
Electrophysiological characterization of neuronal conductivity and seizure susceptibility of knockdown and AP-2µ^R168W^ flies. **(A)** Electrophysiological setup including two tungsten stimulation electrodes in the brain, a saline filled glass recording electrode in the DLM, and an abdominal tungsten electrode as reference. **(B)** Representative giant fiber response measurements at the DLM in response to a single pulse (SP) with 0.1 ms duration and 10 V amplitude at the brain with a latency of ∼1.4 ms. No differences were observed between *AP-2µ* deficiency, *eas^2f^*or control flies. **(C)** Representative electrophysiological recording from the DLM (blue top trace) during stimulus application (orange bottom trace) in a male Canton-S fly. A 2 s long HFS with 0.1 ms stimulus duration at 200 Hz induces seizure-like activity at the DLM. This is characterized by an initial seizure, a period of unresponsiveness of the GFS termed synaptic failure, and a recovery seizure, after which the GFS is responsive to 0.1 ms SPs again. **(D)** Incremental seizure induction protocol starting at 5 V HFS and increasing in 5 V steps every 5 minutes up to 30 V. Pie charts show the proportion of seizing flies at any time during the stimulation protocol. *AP-2µ* knockdown and the *AP-2µ^R168W^* flies exhibited significantly less seizures. Fisher’s exact test (*p < 0.05, **p < 0.01). The calculated voltage threshold for flies at which seizures could be induced is displayed in table 1. **(E)** During the protocol starting at 20 V, no significant differences in seizure occurrence were observed.

**Table 1:**
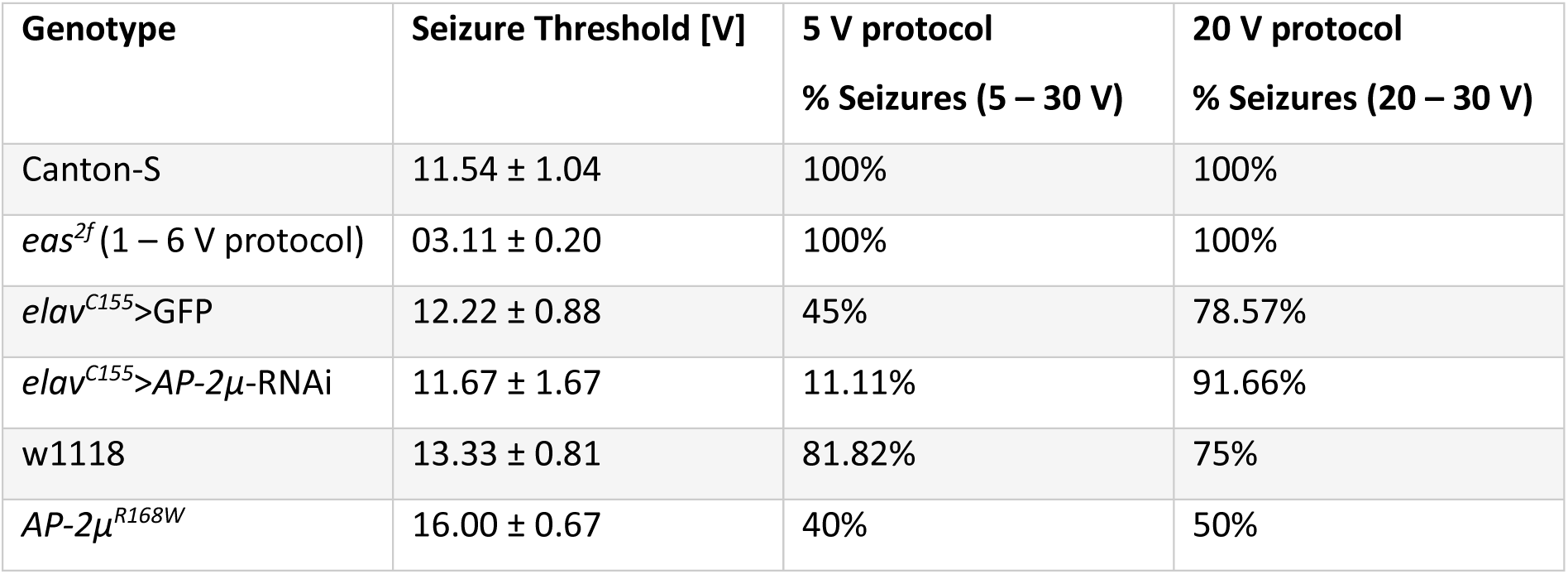
Genotype and seizure threshold based on successful seizure inductions in the 5 – 30 V. Voltages are displayed as mean plus standard error of the mean. Percentage of flies showing seizure-like activity at the DLM at any point during the protocol for 5 – 30 V and 20 – 30 V. (Canton-S: 5-30V n =13; 20-30V n=13, *eas^2f^*: 1-6V n=9; 20-30V n=8)

### AP-2µ knockdown flies exhibit altered morphology of c4da-neurons

Dysfunction of endo- and exocytosis has been linked to abnormal neuron morphology in *Drosophila* (Choudhury et al., 2016; Peng et al., 2015; Yang et al., 2011; Zong et al., 2018). To investigate the role of *AP-2µ* in controlling neuron morphology, we knocked-down *AP-2µ* in c4da-neurons.

We found that *AP-2µ* knockdown led to a reduction in branching pattern complexity (Fig. 5 A). Specifically, the number of branch points and terminals were reduced (branch points: control 519.8 ± 38.36, *AP-*2µ knock-down 371.8 ±24.23; terminals: control 549.3 ± 42.11, *AP-*2µ knock-down 403.8 ± 26.08) (Fig. 5 B, C) while the total length of processes showed no difference (control 16,174 µm ± 1264; *AP-*2µ knock-down 13,772 µm ± 794.5) (Fig. 5 D).

**Figure 5:**
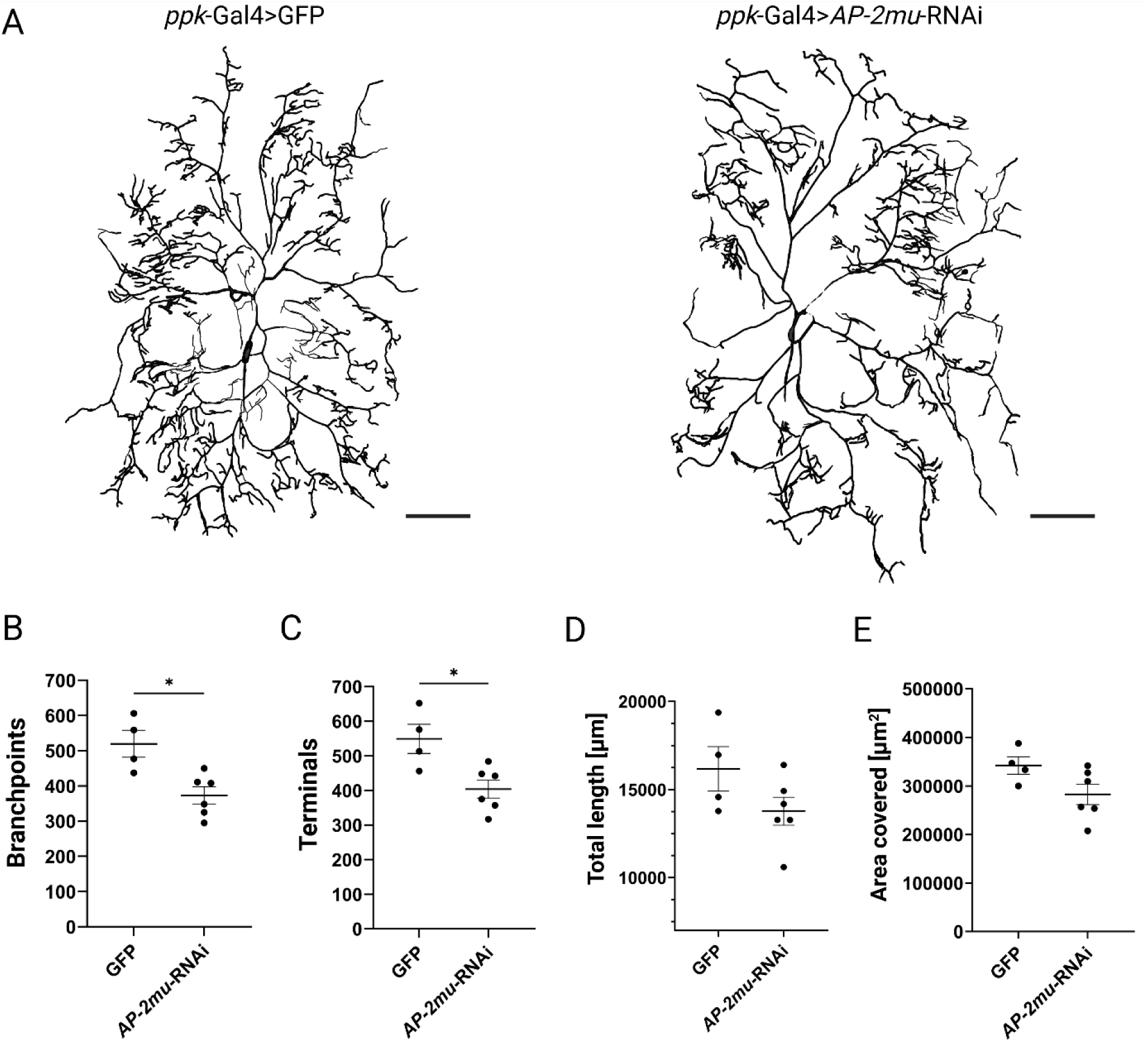
Morphological and synaptic defects in *AP-2µ* knockdown. **(A)** Representative reconstructions of c4da-neurons in L3 larvae expressing either *ppk*-Gal4>GFP as control (left) or *ppk*-Gal4>*AP-2µ*-RNAi (right). Neurons are oriented with anterior to the left and lateral to the top of the image. The *AP-2µ* RNAi-expressing neurons show reduced branching complexity compared to controls. Scalebar equals 100 µm. **(B)**, **(C)** Knockdown of *AP-2µ* led to a reduction in number of branch points and end terminals. **(D)**, **(E)** The total length of processes and the area covered by the dendritic arbor did not change significantly. Values displayed with SEM; Mann-Whitney-U test (*p < 0.05, **p < 0.01).

Convex hull analysis (Fig. S1) further revealed that the maximum spatial extent of the neurons remained unchanged upon *AP-2µ* knock-down (control 341,955 µm^2^ ± 18,200, *AP-2µ* knock-down 282,582 µm^2^ ± 21,074) (Fig. 5 E). Taken together, the knock-down of *AP-2µ* in c4da-neurons reduced the complexity of dendritic arborization.

## Discussion

In this study, we used *Drosophila melanogaster* to model *AP2M1-*associated DEE by generating a *Drosophila* line with pan-neuronal knockdown of *AP-2µ,* the *Drosophila* ortholog of the human *AP2M1* gene, as well as a CRISPR/Cas9-engineered humanized fly carrying the established recurrent R168W variant (analogous to the human R170W variant (Helbig et al., 2019)). In summary, *AP-2µ* knockdown flies exhibited a stable phenotype, characterized by thermosensitivity and alterations of neuronal morphology. Specifically, the heat-induced paralysis observed in these flies was reminiscent of heat-induced seizures seen in flies carrying pathogenic variants in the *SCN1A*-analogous fly gene *para (para^GEFS+^* or *para^DS^*), which mimic *SCN1A*-associated epilepsy syndromes (Roemmich et al., 2021; Schutte et al., 2014; Sun et al., 2012). In contrast to these models, which remain paralyzed throughout the trial after a certain time point, the *AP-2µ* knock-down flies shifted between seizure-like behavior and upright positions, which suggested a non-seizure origin of the paralysis. Furthermore, the paralysis phenotype after *AP-2µ* knock-down was not responsive to antiseizure treatment (Fischer et al., 2024) and flies exhibited increased resistance to electrical seizure induction. *AP-2µ^R168W^* flies showed a similar, albeit less pronounced, resistance to seizures. These findings suggest that *AP-2µ* deficiency in *Drosophila* is associated with a reproducible phenotype, but this phenotype does not entirely align with the epilepsy phenotype observed in humans. This discrepancy is not uncommon and has been reported in other genetic disorders (Bellen et al., 2019; Yamamoto et al., 2014). For instance, the *para^ts1^*allele also shows heat sensitivity along with increased resistance to seizure induction, a similar phenomenon to what we observed in the *AP-2µ* knockdown flies in the present study (Grigliatti et al., 1973; Pavlidis & Tanouye, 1995).

Previous clinical and functional data from our group suggest that the R170W variant causes impairment of CME by destabilizing the AP-2 complex, leading to defective synaptic vesicle recycling and a reduction in transferrin uptake in cell culture assays (Helbig et al., 2019). Additionally, a heat-sensitive paralysis phenotype has been described for the knockdown of *AP-2σ* in *Drosophila*, which encodes another AP-2 subunit (Choudhury et al., 2016). Mutations such as *shibire^ts^*also show thermosensitive paralysis (Kosaka & Ikeda, 1983). Interestingly, while knockdown of the α- and β2-subunit of AP2 was found to be lethal, knockdown of the µ2-subunit did not result in any observable phenotype (Choudhury et al., 2016), which contrasts with our findings. This discrepancy might be attributed to accumulated genetic differences between strains maintained in different laboratories or differences in the methodologies used for heat assays, such as using a sushi cooker for heating (Choudhury et al., 2016).

Given the resistance to anti-seizure treatment and to electrical seizure induction, we hypothesize that the heat-sensitivity in *AP-2µ* deficiency flies is similar to defects observed in other CME genes and likely associated with a failure in synaptic transmission. This could result from insufficient replenishment of synaptic vesicles (Choudhury et al., 2016; Jung et al., 2015; Rohrbough & Broadie, 2002). Of note, the s*hibire^ts^* allele - *shibire* being orthologous to human *DNM1* – also causes heat-related paralysis (Kosaka & Ikeda, 1983). Dynamin, the gene product of *shibire*, is a membrane-remodelling GTPase crucial for membrane fission, the final step in CME (Ferguson & De Camilli, 2012). Dynamin deficiency severely impairs synaptic vesicle recycling and leads to altered membrane structures with an accumulation of collared pits (Kosaka & Ikeda, 1983). Interestingly, *shibire^ts^*also reduces seizure occurrence in established *Drosophila* seizure models, including those involving *para*, *eas* or *sda* (Kroll et al., 2015). In these flies, heat treatment significantly reduced seizure susceptibility in behavioral and seizure induction assays. This effect was not limited to *shibire^ts^* but was also observed in mutants of Rab GTPases, which regulate vesicle trafficking (Kroll et al., 2015). Consequently, it was proposed that altering or reducing vesicle recycling might serve as a seizure suppressing mechanism and could be a potential therapeutic target for epilepsies. It is challenging to pinpoint such paralysis as either a result of seizure-like activity or disturbed neuronal-signaling, as spontaneous discharges of the DLM have been observed in *shibire^ts^* and might reflect impaired inhibitory control due to disrupted vesicle recycling, which might play a role in seizure-generation as well (Kroll et al., 2015).

In line with our findings, the human dynamin gene *DNM1* is associated with another form of DEE (von Spiczak et al., 2017), supporting the idea of differing phenotypical outcomes of CME mutations across species. One possible explanation for seizure resistance in CME deficient lines could be the reduced supply of recycled vesicles at the presynaptic membrane. Our observation that electrical seizure induction starting at 5 V resulted in a lower overall likelihood of seizure occurrence, compared to the protocol starting at 20 V, suggests that repetitive sub-seizure threshold stimulation could lead to cumulative vesicle depletion. This is corroborated by studies in *AP-2σ* mutants, where impaired vesicle regeneration during high-frequency stimulation at the NMJ leads to a progressive decline in synaptic transmission and neurotransmitter release (Choudhury et al., 2016).

This raises the question of how seizures can be explained in humans carrying variants in *AP2M1*, *DNM1,* and other genes involved in CME, such as *CDKL5* (Kontaxi et al., 2023), *CLTC* (Sveistrup & Myers, 2024) or Endophilin (Milosevic et al., 2011). A plausible explanation could be that CME defects do not directly cause neuronal hyperexcitability through alterations in synaptic transmission, but rather more generally by affecting neuronal development. Indeed, CME defects have been found to alter neuronal morphology in developing flies. In particular, morphological and functional alterations of NMJ boutons have been observed in Drosophila lines with knockdowns of all AP-2 subunits (Choudhury et al., 2016). Similar effects have also been observed in defects of other CME genes, such as *shibire* (Dickman et al., 2006), *endoA* (Dickman et al., 2006; Guichet et al., 2002; Rikhy et al., 2002), *Dap160* (Koh et al., 2004), *Eps15* (Koh et al., 2007), *stnA* (Petrovich et al., 1993; Stimson et al., 1998), *stnB* (Mohrmann et al., 2008), *Nwk* (Coyle et al., 2004), *Rab11* (Khodosh et al., 2006; Liu et al., 2014), and several others. Although we expected altered NMJ morphology, we could not reliably observe this phenotype (Fig. S2) and focused on a different class of neurons.

When monitoring neuronal morphology in c4da-neurons, we found that *AP-2µ* knockdown larvae exhibited less complex dendrite arborization patterns. This is consistent with previous studies in *shibire^ts^* and *AP-2α*-knockdown flies, which also exhibited reduced dendritic complexity in c4da-neurons (Peng et al., 2015; Yang et al., 2011). Similarly, flies carrying the *AP-2µ^NN20^* loss-off-function allele show defects in dendrite pruning (Zong et al., 2018). Furthermore, c4da-neuron morphology is also disrupted in other CME deficiency lines, such as *Rab5* (Copf, 2014; Satoh et al., 2008; Takayama et al., 2014), *Nak* (Yang et al., 2011) and *Dab* (Hattori et al., 2013).

Further investigation into the phenotypic differences between flies and humans could provide valuable insights into *AP2M1*-DEE. Over the past decade, our understanding of DEE has evolved. Traditionally, epileptic activity itself was considered the primary cause of cognitive and behavioral impairment (Berg et al., 2010), leading to the now less frequently used term ‘epileptic encephalopathy’. While this concept may apply to a subset of disorders, it has become increasingly clear that many DEE-associated conditions exhibit developmental deficits that occur independently of seizure activity, remain unaffected by seizure treatments, and, in some cases, even precede the onset of seizures (Scheffer et al., 2017). For instance, in Dravet syndrome, developmental regression and cognitive decline often arise prior to significant EEG abnormalities (Connolly, 2016). To better reflect these findings, the International League against Epilepsy coined the term ‘developmental and epileptic encephalopathy’ to acknowledge both developmental deficits and seizure activity as separate but interconnected aspects of these disorders.

In the case of *AP2M1*-DEE, our findings suggest that the developmental consequences of *AP2M1* loss-of-function may underlie the more substantial burden of the human disease phenotype, with epilepsy possibly being a secondary phenomenon resulting from defects of neuronal development. This aligns with clinical observations where some of the reported patients, despite achieving seizure freedom, did not experience improvements in cognitive and behavioral outcomes (Helbig et al., 2019). Furthermore, the co-occurrence of epilepsy with ataxia and muscular hypotonia points to a broader impact on the nervous system beyond cortical neurons.

Previous studies in *Drosophila* have demonstrated that disrupting neuronal activity during critical development windows can lead to persistent seizure-like behavior in adult flies (Giachello & Baines, 2015; Hunter et al., 2024), without the presence of a seizure mutation. Similarly, in mice, interference with neuronal activity during critical periods of neurogenesis exacerbates seizure phenotypes (Lybrand et al., 2021). Future studies should aim to identify critical time windows for development *AP2M1* deficiency in *Drosophila* and mammalian models and explore whether targeted interventions during these periods can mitigate disease severity.

## Conclusion

In this study, we investigated *AP2M1* dysfunction in *Drosophila* through *AP-2µ* knockdown and a CRISPR/Cas9-engineered *AP-2µ^R168W^*variant. While the knockdown led to heat-induced paralysis and neurodevelopmental defects, both models showed resistance against electrically induced seizures. This aligns with findings that CME dysfunction can suppress seizures in flies, suggesting that disruptions in vesicle recycling affect neuronal excitability differently across species. Our results underscore the role of CME in neurodevelopment and synaptic function, highlighting the need to investigate how endocytic defects contribute to epilepsy in humans and whether modulating vesicle trafficking could offer therapeutic potential.

## Supplementary Material

**Figure S1:**
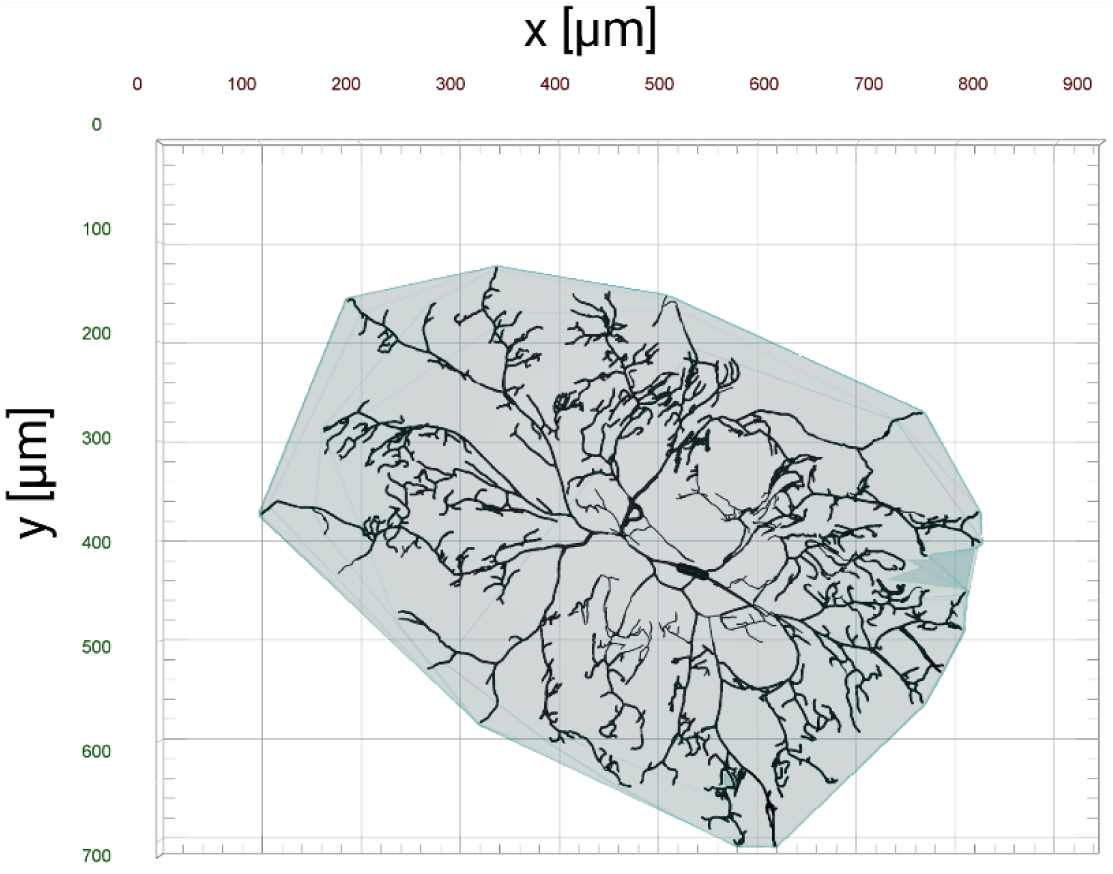
Example of convex hull analysis of a c4da-neuron. A custom pipeline and Python script were used in arivis vision 4D to calculate the smallest convex polygon enclosing all dendritic branches, providing a quantitative measure of the neuron’s spatial extent.

**Figure S2:**
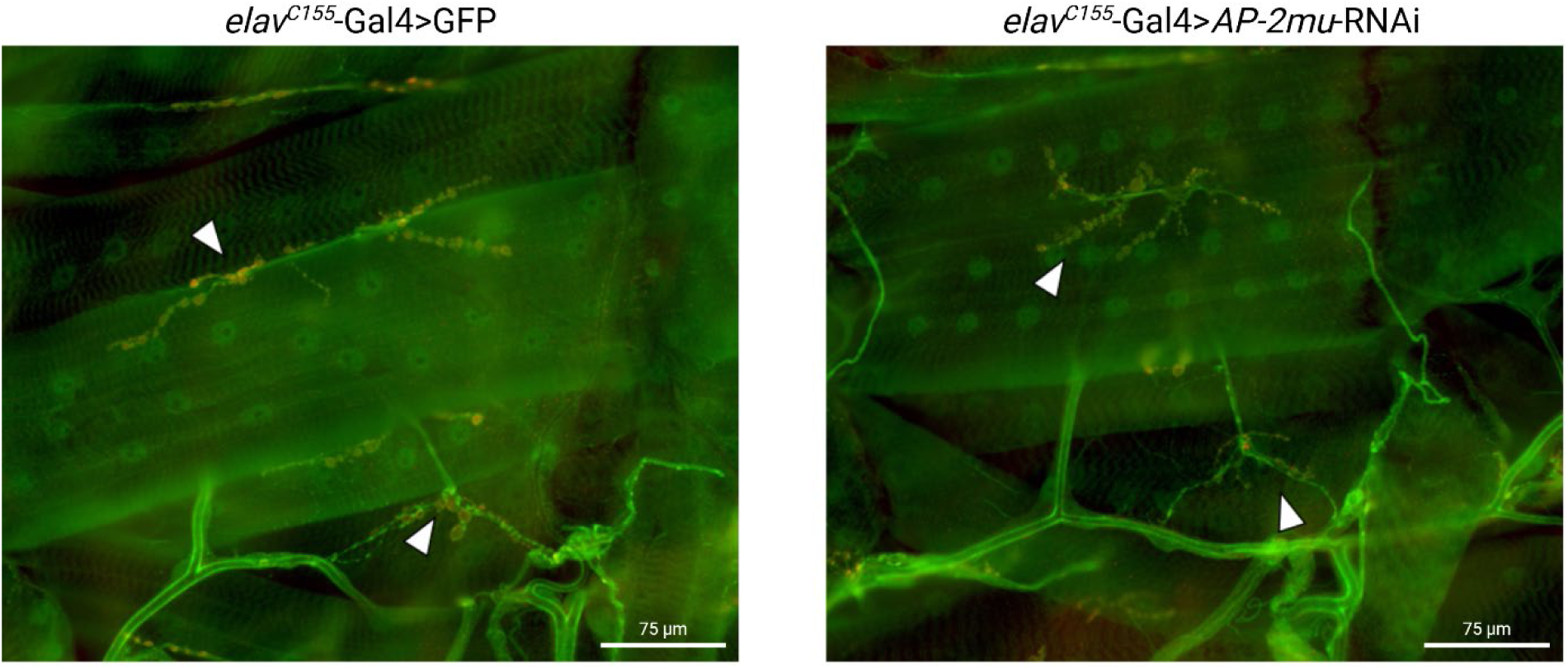
NMJ morphology in *AP-2µ* knockdown larvae. Representative images of NMJs in third instar larvae expressing *elav^C155^*-Gal4 driving either GFP (control, left) or *AP-2µ*-RNAi (knockdown, right). NMJs were labeled with anti-CSP (red) and anti-HRP (green) to visualize synaptic boutons and neuronal membranes. White arrowheads indicate NMJs. The scale bar represents 75 µm. No directly visible morphological differences were observed in *AP-2µ* knockdown larvae.

