## Supplemental Data 1 for "Modeling AP2M1 Developmental and Epileptic Encephalopathy in Drosophila"

### CRISPR Gene Editing Final Report

| | |
|---|---|
| **Report Number** | RWGa4453 |
| **Case Number** | 220862 |
| **Customer ID** | DE031 |
| **Gene** | *Ap-2mu /CG7057* |
| **Service** | CRISPR Service Package |
| **Project** | introducing a point mutation R168W of AP-2mu using PBac system to facilitate genetic screening |
| **Method** | CRISPR/Cas9-mediated genome editing by homology-dependent repair (HDR) using 1guide RNA(s) and a dsDNA plasmid donor |
| **Strain** | *[LWG228] w[1118]* |

****************************************************************************

WELLGENETICS

### Progress Summary

| Landmark | Complete Date | Remark |
|---|---|---|
| Order Received | 2023/02/01 | |
| Sample Received | NA | NA |
| Design Report | 2023/02/02 | RWGa3350 |
| Design Confirmation | 2023/02/08 | |
| Genomic Sequencing | 2023/02/14 | RWGa3459 |
| gRNA Confirmation | 2023/02/21 | |
| Guide RNA Cloning | 2023/03/10 | |
| Donor Plasmid Cloning | 2023/03/14 | |
| Cloning Report | 2023/03/15 | RWGa3634 |
| Plasmids Purification | 2023/03/17 | |
| Microinjection | 2023/03/20 | 256 Embryos |
| G0 Cross | 2023/04/06 | 216 Adult(s) |
| F1 Screen by Visible Marker | 2023/04/21 | 167 Fertile Crosses<br>1 Positive Line(s) |
| PCR Validation | 2023/05/10 | 0 Validated Line(s) |
| **The 2nd round** | | |
| Guide RNA Cloning | 2023/05/25 | |
| Plasmids Purification | 2023/06/02 | |
| Microinjection | 2023/06/08 | 205 Embryos |
| G0 Cross | 2023/06/21 | 162 Adult(s) |
| F1 Screen by Visible Marker | 2023/07/13 | 112 Fertile Crosses<br>3 Positive Line(s) |
| PCR Validation | 2023/07/12 | 3 Validated Line(s) |
| | *220862A SWGa7526* | *w[*];; AP-2mu R168W CRISPR{PBacDsRed} / TM6B, Tb[1]* |
| | *220862B SWGa7527* | *w[*];; AP-2mu R168W CRISPR{PBacDsRed} / TM6B, Tb[1]* |
| | *220862C* | *w[*];; AP-2mu R168W CRISPR{PBacDsRed} /* |

+8862-2651-1809
13F.-12, No. 93, Sec. 1, Xintai 5th Rd., Xizhi Dist., New Taipei City 221416, Taiwan (R.O.C)
wellgenetics.com

WELLGENETICS

| | | |
|---|---|---|
| | *SWGa7528* | *TM6B, Tb[1]* |
| Sequencing Validation | 2023/07/13 | 1 Validated Line(s) |
| | *220862A SWGa7526* | *w[*];; AP-2mu R168W CRISPR{PBacDsRed} / TM6B, Tb[1]* |
| Validation Report | 2023/07/21 | RWGa4451 |
| Integration Report | 2023/07/21 | RWGa4452 |
| Stock Shipment | 2023/08/14 | 3 Isogenized& Balanced Stock(s) |
| | *220862A SWGa7526* | *w[*];; AP-2mu R168W CRISPR{PBacDsRed} / TM6B, Tb[1]* |
| | *220862B SWGa7527* | *w[*];; AP-2mu R168W CRISPR{PBacDsRed} / TM6B, Tb[1]* |
| | *220862C SWGa7528* | *w[*];; AP-2mu R168W CRISPR{PBacDsRed} / TM6B, Tb[1]* |
| | 2023/08/14 | 1 Control Stock(s) |
| | *LWG228* | *w[1118]* |
| Final Report | 2023/07/21 | RWGa4453 |
| Stock Keeping Deadline | 2023/11/14 | Please make sure to characterize these lines and let us know any problems before the keeping deadline. |

+8862-2651-1809
13F.-12, No. 93, Sec. 1, Xintai 5th Rd., Xizhi Dist., New Taipei City 221416, Taiwan (R.O.C)
wellgenetics.com

### Method Summary

CRISPR-mediated mutagenesis was performed by WellGenetics Inc. using modified methods of Kondo and Ueda (2013). In brief, gRNAs sequence(s) TCACGTACTCCAATACGTCC[AGG] and GACCCTGCGGGCTCATCAGC[AGG] were cloned into U6 promoter plasmid(s). Cassette PBacDsRed, which contains 3xP3-DsRed flanked by PiggyBac terminal repeats, and two homology arms with point mutation(s) were cloned into pUC57-Kan as donor template for repair. *Ap-2mu/CG7057*-targeting gRNAs and hs-Cas9 were supplied in DNA plasmids, together with donor plasmid for microinjection into embryos of control strain *w[1118]*. F1 flies carrying selection marker of 3xP3-DsRed were further validated by genomic PCR and sequencing. CRISPR generates a break in *Ap-2mu/CG7057,* and is replaced by cassette PBacDsRed.

Please cite WellGenetics Inc. when using this CRISPR line in a publication.

+8862-2651-1809
13F.-12, No. 93, Sec. 1, Xintai 5th Rd., Xizhi Dist., New Taipei City 221416, Taiwan (R.O.C)
wellgenetics.com

WELLGENETICS

### CRISPR Design Report

**Report No.:** RWGa3350 **Date:** 2023.02.02

**************************************************************************

**Gene:** *Ap-2mu/CG7057* **Case No.:** 220862

**Project:** introducing a point mutation R168W of AP-2mu using PBac system to facilitate genetic screening

**Method:** CRISPR/Cas9-mediated genome editing by homology-dependent repair (HDR) using 1 guide RNA(s) and a dsDNA plasmid donor

+8862-2651-1809

13F.-12, No. 93, Sec. 1, Xintai 5th Rd., Xizhi Dist., New Taipei City 221416, Taiwan (R.O.C)

wellgenetics.com

#### Gene and Method

**Gene**: *Ap-2mu/CG7057*

**Location:** 3R (94A15-94A16)

**Method:** CRISPR/Cas9-mediated genome editing by homology-dependent repair (HDR) using a guide RNA and a dsDNA plasmid donor

**Knock-in cassette:** PBacDsRed

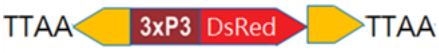

**Injection strain:** *w[1118]*

**Genome Editing:**

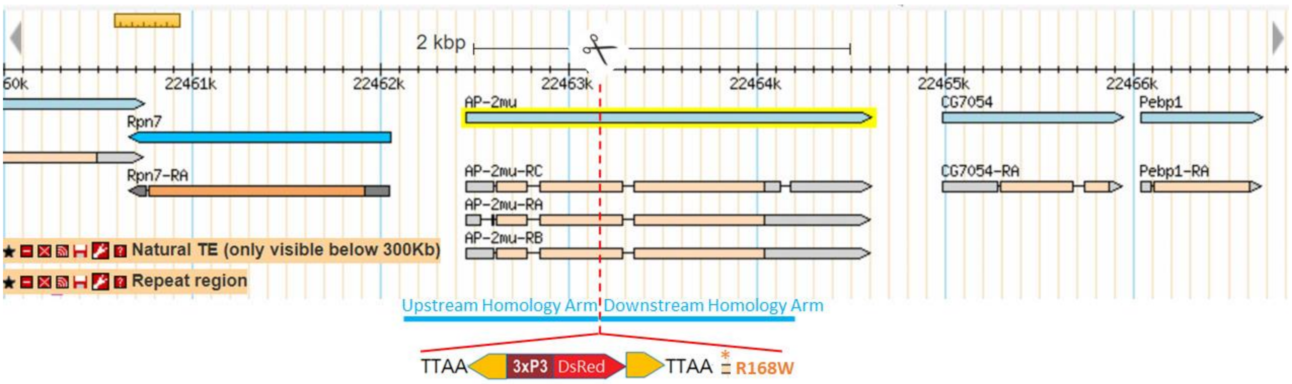

(1) Introduce a point mutation R168 to W, CGC to TGG of *AP-2mu*.

(2) The selection marker PBacDsRed contains 3' PBac terminal repeats, the artificial 3xP3 promoter (three tandem copies of the Pax-6 homodimer binding site and TATA-homology of *hsp70*), DsRed2, SV40 3'UTR, and 5' PBac terminal repeats. It facilitates the genetic screening and can be excised by Piggy Bac transposase. Only one TTAA motif will be left after transposition embedded in mutated exon sequence, and create a silent mutation on I164 of AP-2mu; I164K165 ATCAAG to ATTAAG, IK to IK.

(3) Selection marker needs to be excised when looking at endogenous gene/protein expression.

(4) PAM mutations to make the donor inactive to guide RNA will be incorporated into the edited genome.

+8862-2651-1809
13F.-12, No. 93, Sec. 1, Xintai 5th Rd., Xizhi Dist., New Taipei City 221416, Taiwan (R.O.C)
wellgenetics.com

**CRISPR Target Sites**

This is a targeting design without deletion. In order to keep the possibility of using backup gRNA(s), we include PAM mutations in the donor construct for two or more gRNAs. We will use gRNA 2 in case the first one fails.

**Guide RNA Quality Instruction**

| | Distance | Off target | GC Content | T # |
|---|---|---|---|---|
| ● Strong site | <50bp | 0 | 45%-70% | ≦1 |
| ● Weak site | 50-100bp | Potential off targets on untargeted chromosomes | 40%, 75% | 2 |
| ● Bad site | >100bp | Potential off targets on targeted chromosomes | < 40%, >75% | ≧3 |

1. The distance from knockin site or deletion breakpoint to cutting site of Cas9.
2. Predicted off targets on other chromosomes are indicated.
3. GC content 45% to 70% of target sequence provides the best efficiency.
4. High thymine numbers at 17$^{th}$ to 20$^{th}$ nucleotide of the guide RNA target site may decrease the efficiency.

**gRNA 1**

**CRISPR Target Site [PAM]:** CCAGATTGGCTGGCGTCGCG[AGG]

**CRISPR Target Strand in the Genome:** plus

**Cutting Site:** +549 nt from ATG of *AP-2mu*

**Distance:** -9 nt from upstream breakpoint to cutting site of Cas9 ●

-13 nt from downstream breakpoint to cutting site of Cas9 ●

**GC Content:** 70% ●

**T number at 17$^{th}$ to 20$^{th}$ Nucleotides:** 0 ●

**Off target:** 0 ●

**Guide RNA Primers:**

Sense oligo 5’- CTTCGCCAGATTGGCTGGCGTCGCG

Antisense oligo 5’- AAACCGCGACGCCAGCCAATCTGGC

**PAM mutation:** CCAGATTGGCTGGCGTCGCG[AAG]

GAG to GAA, E to E

☎ +8862-2651-1809 ✉
13F.-12, No. 93, Sec. 1, Xintai 5th Rd., Xizhi Dist., New Taipei City 221416, Taiwan (R.O.C)
wellgenetics.com

**Upstream Homology Arm:** 1,076bp, the -518 nt to +558 nt from ATG of *AP-2mu*

Forward Oligo 5’- ACCAGAGCGATGTCCATCAAG

Reverse Oligo 5’- TGCCTTCGCGACGCCAGCCAAT

**Downstream Homology Arm:** 1,074bp, the +563 nt to +1,636 nt from ATG of *AP-2mu*

Synthesis fragment:

5’- GTACCGGTGGAACGAGCTTTTTCTGGACGTATTGGAGTACGTGAACCTGCTGATGAGCCC

Forward Oligo: 5’- GCAGGGTCAGGTTTTGTCTG

Reverse Oligo: 5’- CGGTATACGGCATAGGGTGT

#### gRNA 2

**CRISPR Target Site [PAM]**: TCACGTACTCCAATACGTCC[AGG]

**CRISPR Target Strand in the Genome:** minus

**Cutting Site:** +589 nt from ATG of *AP-2mu*

**Distance:** +31 nt from upstream breakpoint to cutting site of Cas9 ●

+27 nt from downstream breakpoint to cutting site of Cas9 ●

**GC Content:** 50% ●

**T number at 17$^{th}$ to 20$^{th}$ Nucleotides:** 1 ●

**Off target:** 0 ●

**Guide RNA Primers:**

Sense oligo 5’- CTTCGTCACGTACTCCAATACGTCC

Antisense oligo 5’- AAACGGACGTATTGGAGTACGTGAC

**PAM mutation:** TCACGTACTCCAATACGTCC[AGA]

TTC to TTT, F to F

**Upstream Homology Arm:** same as above

**Downstream Homology Arm:** same as above

☎ +8862-2651-1809 ✉
13F.-12, No. 93, Sec. 1, Xintai 5th Rd., Xizhi Dist., New Taipei City 221416, Taiwan (R.O.C)
wellgenetics.com

**Donor Design**

#### Annotation

NNNNN: Homology Arm / Coding Region

NNNNN: Homology Arm / UTR

nnnnn: Homology Arm/ Intron

nnnnn: Homology Arm/ Intergenic Region

nnnnn: gRNA

**nnnnnn**: Cloning Site

N: Silent mutation

N: Point mutation

nnnn: TTAA

nnnnn: Piggy Bac terminal repeats

nnnnn: 3xP3-hsp70 promoter

NNNNN: DsRed2

nnnnn: SV40 polyA

#### Sequence

accagagcgatgtccatcaagatagcgacgaaattagaacagtgcaattgccaattgggaatttgtattttaatttatttttaaattctgaaag
taatttaaatttaaaaaaaaaacttgagagctgtctagaaaagaactgatgtttcatgataactttgtcgaagaattaagaaatatttagttgt
aaaataattgttgaatctattttttccaataacacgacttatatattttttgaaaatattcgagctaaatcccaagaagtaaactcaatctggg
atttgaagtgcccagaactcgaataaacacttctttttaaataattgtaagaccgtatcacttatggtatatactgacctcgaagggCCACAC
TAAGGGGGAGTGAAAATTGATTTTCTGATAAAAATTTTCGCTTGAAGCTACAGCATCGTCCACTGTCCATgta
tatatcttatatttgcatataaatatatatattacaccgacttggactaaccatcagATAGCGCACAAGATGATTGGCGGCCTGTT
CGTCTACAACCACAAGGGCGAGGTGCTGATCTCGCGAGTTTACCGCGACGACATCGGTCGGAATGCCGTG
GACGCCTTTCGGGTCAACGTCATCCACGCCCGCCAGCAGGTCCGCTCGCCAGTGACCAATATTGCGAGGAC
CAGCTTCTTCCACATCAAGgtcggtgaacaaagttcctatgtcacaatcaccaaccctcccacttatcatcaaaactcctttcacagA
GAGCAAACATTTGGCTGGCGGCTGTGACCAAGCAGAATGTGAACGCCGCGATGGTGTTTGAGTTCCTTTTG
AAGATCATCGAGGTGATGCAATCCTACTTCGGCAAGATCTCGGAGGAGAACATCAAGAATAACTTCGTGCT
CATCTACGAGCTGCTGGATGAGATCCTCGACTTTGGCTACCCGCAGAACACGGACTCCGGCACCCTGAAGA
CCTTCATCACACAGCAGGGCATCAAGTCGGCCACCAAGGAGGAGCAGATGCAGATAACCTCGCAGGTTACC
GGCCAGATTGGCTGGCGTCGCGAAGGCA
TTAAccctagaaagataatcatattgtgacgtacgttaaagataatcatgcgtaaaattgacgcatgtgttttatcggtctgtatatcgaggttt
atttattaatttgaatagatattaagttttattatatttacacttacatactaataataaattcaacaaacaatttatttatgtttatttatttattaaaa
aaaaacaaaaactcaaatttcttctataaagtaacaaaacttttaggatctaattcaattagagactaattcaattagagctaattcaattagg
atccaagcttatcgatttcgaaccctcgaccgccggagtataaatagaggcgcttcgtctacggagcgacaattcaattcaaacaagcaaagtg

+8862-2651-1809
13F.-12, No. 93, Sec. 1, Xintai 5th Rd., Xizhi Dist., New Taipei City 221416, Taiwan (R.O.C)
wellgenetics.com

aacacgtcgctaagcgaaagctaagcaaataaacaagcgcagctgaacaagctaaacaatcggctcgaagccggtcgccaccATGGCCT
CCTCCGAGGACGTCATCAAGGAGTTCATGCGCTTCAAGGTGCGCATGGAGGGCTCCGTGAACGGCCACGAG
TTCGAGATCGAGGGCGAGGGCGAGGGCCGCCCCTACGAGGGCACCCAGACCGCCAAGCTGAAGGTGACCA
AGGGCGGCCCCCTGCCCTTCGCCTGGGACATCCTGTCCCCCCAGTTCCAGTACGGCTCCAAGGTGTACGTGA
AGCACCCCGCCGACATCCCCGACTACAAGAAGCTGTCCTTCCCCGAGGGCTTCAAGTGGGAGCGCGTGATGA
ACTTCGAGGACGGCGGCGTGGTGACCGTGACCCAGGACTCCTCCCTCCAGGACGGCTCCTTCATCTACAAGG
TGAAGTTCATCGGCGTGAACTTCCCCTCCGACGGCCCCGTAATGCAGAAGAAGACTATGGGCTGGGAGGCGT
CCACCGAGCGCCTGTACCCCCGCGACGGCGTGCTGAAGGGCGAGATCCACAAGGCCCTGAAGCTGAAGGAC
GGCGGCCACTACCTGGTGGAGTTCAAGTCCATCTACATGGCCAAGAAGCCCGTGCAGCTGCCCGGCTACTACT
ACGTGGACTCCAAGCTGGACATCACCTCCCACAACGAGGACTACACCATCGTGGAGCAGTACGAGCGCGCCG
AGGGCCGCCACCACCTGTTCCTGTAGcggccgcgactctagatcataatcagccataccacatttgtagaggttttacttgctttaaa
aaacctcccacacctccccctgaacctgaaacataaaatgaatgcaattgttgttgttaacttgtttattgcagcttataatggttacaaataaag
caatagcatcacaaatttcacaaataaagcatttttttcactgcattctagttgtggtttgtccaaactcatcaatgtatcttagatatctataacaa
gaaaatatatatataataagttatcacgtaagtagaacatgaaataacaatataattatcgtatgagttaaatcttaaaagtcacgtaaaagat
aatcatgcgtcattttgactcacgcggtcgttatagttcaaaatcagtgacacttaccgcattgacaagcacgcctcacgggagctccaagcggc
gactgagatgtcctaaatgcacagcgacggattcgcgctatttagaaagagagagcaatatttcaagaatgcatgcgtcaattttacgcagact
atctttctagggttaa
GTACCGGTGGAACGAGCTTTTTCTGGACGTATTGGAGTACGTGAACCTGCTGATGAGCCC
GCAGGGTCAGGTTTTGTCTGCCCACGTGGCCGGCAAGGTGGTAATGAAGTCGTATTTGTCGGgtaagttagcat
aaataatctagacattattccttttaataatcgccatgttgtagGCATGCCCGAGTGCAAGTTCGGGATTAACGACAAGATC
GTGATGGAGTCTAAGGGACGCGGTCTCTCCGGAAATTCAGAGGCGGAAACCTCACGCTCCGGCAAGCCCG
TCGTGGTCATCGATGACTGCCAGTTCCATCAGTGCGTCAAGCTAAGCAAATTCGAGACGGAGCATTCGATC
AGCTTCATCCCGCCGGACGGGGAGTTCGAGCTGATGCGTTACCGTACCACCAAAGACATTTCGCTGCCATTC
CGAGTCATCCCGCTGGTGCGGGAGGTGGGCCGCACCAAGATGGAGGTTAAGGTTGTGCTGAAGTCCAACT
TTAAGCCCTCACTGCTGGGCCAAAAGATCGAGGTGAAGATACCAACCCCGCTCAATACATCGGGCGTGCAG
CTCATCTGCCTAAAGGGCAAAGCCAAATATAAGGCTTCGGAGAACGCGATCGTGTGGAAGATTAAGCGCAT
GGCGGGCATGAAGGAGACACAGCTGTCCGCGGAAATCGAACTTTTGGAGACGGACACCAAGAAGAAGTG
GACTCGGCCGCCCATCTCCATGAACTTTGAGGTGCCATTCGCGCCGTCCGGCTTCAAGGTACGCTACCTGAA
GGTGTTCGAGCCCAAGCTCAACTACTCCGACCACGATGTGGTCAAATGGGTGCGCTACATCGGACGCAGTG
GCTTGTATGAGACGCGCTGCTAGGGCGCCCAAGCATCCCCATCACGGAGCAAGATTATCACAATTGAATTT
AACGAGATGAATGGAGAAAGTGTTACCGCATCTATTGGTATTTCCTCAATCAAATTGAACCAGCAGCTTATG
TATTCCAAGATTAGTTCCAGCAAGTCCGGAGGACGCCCGAGTAGCGGATCCACCCCGGACACCCTATGCCG
TATACCG

+8862-2651-1809
13F.-12, No. 93, Sec. 1, Xintai 5th Rd., Xizhi Dist., New Taipei City 221416, Taiwan (R.O.C)
wellgenetics.com

**Translation of wildtype**

NNNNN: the coding exon2 of *AP-2mu*

nnnnn: gRNA

N: expecting silent mutation

NNNN: expecting TTAA

N: expecting point mutation

**Nucleotides:**

AGAGCAAACATTTGGCTGGCGGCTGTGACCAAGCAGAATGTGAACGCCGCGATGGTGTTTGAGTTCCTTTT
GAAGATCATCGAGGTGATGCAATCCTACTTCGGCAAGATCTCGGAGGAGAACATCAAGAATAACTTCGTGC
TCATCTACGAGCTGCTGGATGAGATCCTCGACTTTGGCTACCCGCAGAACACGGACTCCGGCACCCTGAAG
ACCTTCATCACACAGCAGGGCATCAAGTCGGCCACCAAGGAGGAGCAGATGCAGATAACCTCGCAGGTTAC
CGGCCAGATTGGCTGGCGTCGCGAGGGCATCAAGTACCGGCGCAACGAGCTTTTCCTGGACGTATTGGAG
TACGTGAACCTGCTGATGAGCCCGCAGGGTCAGGTTTTGTCTGCCCACGTGGCCGGCAAGGTGGTAATGAA
GTCGTATTTGTCGG

**Protein:**

R A N I W L A A V T K Q N V N A A Met V F E F L L K I I E V Met Q S Y F G K I S E E N I K N N F V L I Y E
L L D E I L D F G Y P Q N T D S G T L K T F I T Q Q G I K S A T K E E Q Met Q I T S Q V T G Q I G W R R E
G I K Y R R N E L F L D V L E Y V N L L Met S P Q G Q V L S A H V A G K V V Met K S Y L S

**Translation of *AP-2mu R168W* after excision**

NNNNN: the coding exon2 of *AP-2mu R168W*

nnnnn: gRNA

N: Silent mutation

NNNN: TTAA

N: Point mutation

**Nucleotides:**

AGAGCAAACATTTGGCTGGCGGCTGTGACCAAGCAGAATGTGAACGCCGCGATGGTGTTTGAGTTCCTTTT
GAAGATCATCGAGGTGATGCAATCCTACTTCGGCAAGATCTCGGAGGAGAACATCAAGAATAACTTCGTGC
TCATCTACGAGCTGCTGGATGAGATCCTCGACTTTGGCTACCCGCAGAACACGGACTCCGGCACCCTGAAG
ACCTTCATCACACAGCAGGGCATCAAGTCGGCCACCAAGGAGGAGCAGATGCAGATAACCTCGCAGGTTAC
CGGCCAGATTGGCTGGCGTCGCGAAGGCATTAAGTACCGGTGGAACGAGCTTTTTCTGGACGTATTGGAG

☎ +8862-2651-1809 ✉
13F.-12, No. 93, Sec. 1, Xintai 5th Rd., Xizhi Dist., New Taipei City 221416, Taiwan (R.O.C)
wellgenetics.com

TACGTGAACCTGCTGATGAGCCCGCAGGGTCAGGTTTTGTCTGCCCACGTGGCCGGCAAGGTGGTAATGAA
GTCGTATTTGTCGG

**Protein:**

R A N I W L A A V T K Q N V N A A Met V F E F L L K I I E V Met Q S Y F G K I S E E N I K N N F V L I Y E
L L D E I L D F G Y P Q N T D S G T L K T F I T Q Q G I K S A T K E E Q Met Q I T S Q V T G Q I G W R R E
G I K Y R W N E L F L D V L E Y V N L L Met S P Q G Q V L S A H V A G K V V Met K S Y L S

+8862-2651-1809
13F.-12, No. 93, Sec. 1, Xintai 5th Rd., Xizhi Dist., New Taipei City 221416, Taiwan (R.O.C)
wellgenetics.com

### CRISPR Design and Genomic Sequencing Report

**Report No.:** RWGa3459 **Date:** 2023.02.14

*************************************************************************

**Gene:** *Ap-2mu/CG7057* **Case No.:** 220862

**Project:** introducing a point mutation R168W of AP-2mu using PBac system to facilitate genetic screening

**Method:** CRISPR/Cas9-mediated genome editing by homology-dependent repair (HDR) using 1 guide RNA(s) and a dsDNA plasmid donor

+8862-2651-1809
13F.-12, No. 93, Sec. 1, Xintai 5th Rd., Xizhi Dist., New Taipei City 221416, Taiwan (R.O.C)
wellgenetics.com

**Gene and Method**

**Gene**: *Ap-2mu/CG7057*

**Location:** 3R (94A15-94A16)

**Method:** CRISPR/Cas9-mediated genome editing by homology-dependent repair (HDR) using a guide RNA and a dsDNA plasmid donor

**Knock-in cassette:** PBacDsRed

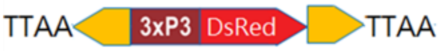

**Injection strain:** *w[1118]*

**Genome Editing:**

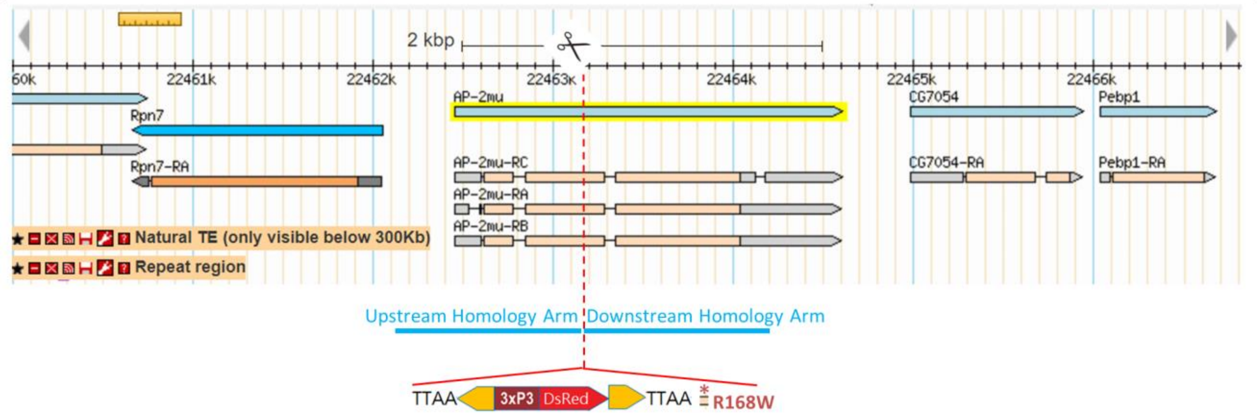

(1) Introduce a point mutation R168 to W, CGC to TGG of *AP-2mu*.

(2) The selection marker PBacDsRed contains 3’ PBac terminal repeats, the artificial 3xP3 promoter (three tandem copies of the Pax-6 homodimer binding site and TATA-homology of *hsp70*), DsRed2, SV40 3’UTR, and 5’ PBac terminal repeats. It facilitates the genetic screening and can be excised by Piggy Bac transposase. Only one TTAA motif will be left after transposition embedded in mutated exon sequence, and create a silent mutation on I164 of AP-2mu; I164K165 ATCAAG to ATTAAG, IK to IK.

(3) Selection marker needs to be excised when looking at endogenous gene/protein expression.

(4) PAM mutations to make the donor inactive to guide RNA will be incorporated into the edited genome.

+8862-2651-1809
13F.-12, No. 93, Sec. 1, Xintai 5th Rd., Xizhi Dist., New Taipei City 221416, Taiwan (R.O.C)
wellgenetics.com

WellGenetics

**CRISPR Target Sites**

This is a targeting design without deletion. In order to keep the possibility of using backup gRNA(s), we include PAM mutations in the donor construct for two or more gRNAs. We will use gRNA 2 in case the first one fails.

**Guide RNA Quality Instruction**

| | Distance | Off target | GC Content | T # |
|---|---|---|---|---|
| ● Strong site | <50bp | 0 | 45%-70% | ≦1 |
| ● Weak site | 50-100bp | Potential off targets on untargeted chromosomes | 40%, 75% | 2 |
| ● Bad site | >100bp | Potential off targets on targeted chromosomes | < 40%, >75% | ≧3 |

1. The distance from knockin site or deletion breakpoint to cutting site of Cas9.
2. Predicted off targets on other chromosomes are indicated.
3. GC content 45% to 70% of target sequence provides the best efficiency.
4. High thymine numbers at $17^{th}$ to $20^{th}$ nucleotide of the guide RNA target site may decrease the efficiency.

**gRNA 1**

**CRISPR Target Site [PAM]:** CCAGATTGGCTGGCGTCGCG[AGG]

**CRISPR Target Strand in the Genome:** plus

**Cutting Site:** +549 nt from ATG of *AP-2mu*

**Distance:** -9 nt from upstream breakpoint to cutting site of Cas9 ●

-13 nt from downstream breakpoint to cutting site of Cas9 ●

**GC Content:** 70% ●

**T number at $17^{th}$ to $20^{th}$ Nucleotides:** 0 ●

**Off target:** 0 ●

**Guide RNA Primers:**

Sense oligo 5’- CTTCGCCAGATTGGCTGGCGTCGCG

Antisense oligo 5’- AAACCGCGACGCCAGCCAATCTGGC

**PAM mutation:** CCAGATTGGCTGGCGTCGCG[AAG]

GAG to GAA, E to E

☎ +8862-2651-1809 ✉
13F.-12, No. 93, Sec. 1, Xintai 5th Rd., Xizhi Dist., New Taipei City 221416, Taiwan (R.O.C)
wellgenetics.com

**Upstream Homology Arm:** 1,076bp, the -518 nt to +558 nt from ATG of *AP-2mu*

Forward Oligo 5'- ACCAGAGCGATGTCCATCAAG

Reverse Oligo 5'- TGCCTTCGCGACGCCAGCCAAT

**Downstream Homology Arm:** 1,074bp, the +563 nt to +1,636 nt from ATG of *AP-2mu*

Synthesis fragment:

5'- GTACCGGTGGAACGAGCTTTTTCTGGACGTATTGGAGTACGTGAACCTGCTGATGAGCCC

Forward Oligo: 5'- GCAGGGTCAGGTTTTGTCTG

Reverse Oligo: 5'- CGGTATACGGCATAGGGTGT

#### gRNA 2

**CRISPR Target Site [PAM]:** TCACGTACTCCAATACGTCC[AGG]

**CRISPR Target Strand in the Genome:** minus

**Cutting Site:** +589 nt from ATG of *AP-2mu*

**Distance:** +31 nt from upstream breakpoint to cutting site of Cas9 ●

+27 nt from downstream breakpoint to cutting site of Cas9 ●

**GC Content:** 50% ●

**T number at 17th to 20th Nucleotides:** 1 ●

**Off target:** 0 ●

**Guide RNA Primers:**

Sense oligo 5'- CTTCGTCACGTACTCCAATACGTCC

Antisense oligo 5'- AAACGGACGTATTGGAGTACGTGAC

**PAM mutation:** TCACGTACTCCAATACGTCC[AGA]

TTC to TTT, F to F

**Upstream Homology Arm:** same as above

**Downstream Homology Arm:** same as above

**Genomic Sequencing Results:**

>w1118_5_OWGb2734

GGATGGAGGAACTCAGATAACTTCGTGCTCATCTACGAGCTGCTGGATGAGATCCTCGACTTTGGCTACCCGC
AGAACACGGACTCCGGTACCCTGAAGACCTTCATCACACAGCAGGGCATCAAGTCGGCCACCAAGGAGGAG
CAGATGCAGATAACCTCGCAGGTCACCGGCCAGATTGGCTGGCGTCGTGAGGGCATCAAGTACCGGCGCAAC
GAGCTTTTCCTGGACGTATTGGAGTACGTGAACCTGCTGATGAGCCCGCAGGGTCAGGTTTTGTCTGCTCACG
TGGCCGGCAAGGTGGTAATGAAGTCGTATTTGTCGGGTAAGTTAGCATAAATAATCTAGACATTATTCCTTTTAA
TAATCGCCATGTTGTAGGCATGCCCGAGTGCAAGTTCGGCATTAACGACAAGATCGTGATGGAGTCTAAGGGA
CGCGGTCTCTCCGGAAATTCAGAGGCGGAAACCTCACGCTCCGGCAAGCCCGTCGTGGTCATCGATGACTGC

+8862-2651-1809
13F.-12, No. 93, Sec. 1, Xintai 5th Rd., Xizhi Dist., New Taipei City 221416, Taiwan (R.O.C)
wellgenetics.com

CAGTTCCATCAGTGCGTCAAGCTAAGCAAATTCGAGACGGAGCATTCGATCAGCTTTATCCCGCCGGACGGG
GAGTTCGAGCTGATGCGTTACCGTACCACCAAAGACATTTCGCTGCCATTCCGAGTCATCCCGCTGGTGCGGG
AGGTGGGCCGCACCAGGGGGAGAGGAGGATATAAA

>w1118_6_OWGb2734
GGGATAGGACTCAGATAACTTCGTGCTCATCTACGAGCTGCTGGATGAGATCCTCGACTTTGGCTACCCGCAG
AACACGGACTCCGGTACCCTGAAGACCTTCATCACACAGCAGGGCATCAAGTCGGCCACCAAGGAGGAGCA
GATGCAGATAACCTCGCAGGTCACCGGCCAGATTGGCTGGCGTCGTGAGGGCATCAAGTACCGGCGCAACG
AGCTTTTCCTGGACGTATTGGAGTACGTGAACCTGCTGATGAGCCCGCAGGGTCAGGTTTTGTCTGCTCACGT
GGCCGGCAAGGTGGTAATGAAGTCGTATTTGTCGGGTAAGTTAGCATAAATAATCTAGACATTATTCCTTTTAAT
AATCGCCATGTTGTAGGCATGCCCGAGTGCAAGTTCGGCATTAACGACAAGATCGTGATGGAGTCTAAGGGA
CGCGGTCTCTCCGGAAATTCAGAGGCGGAAACCTCACGCTCCGGCAAGCCCGTCGTGGTCATCGATGACTGC
CAGTTCCATCAGTGCGTCAAGCTAAGCAAATTCGAGACGGAGCATTCGATCAGCTTTATCCCGCCGGACGGG
GAGTTCGAGCTGATGCGTTACCGTACCACCAAAGACATTTCGCTGCCATTCCGAGTCATCCCGCTGGTGCGGG
AGGTGGGCCGCACCAGGGGGAGGGGGTTATAAAA

>w1118_7_OWGb2734
GGGCCGAGGACTCAGAATAACTTCGTGCTCATCTACGAGCTGCTGGATGAGATCCTCGACTTTGGCTACCCGC
AGAACACGGACTCCGGTACCCTGAAGACCTTCATCACACAGCAGGGCATCAAGTCGGCCACCAAGGAGGAG
CAGATGCAGATAACCTCGCAGGTCACCGGCCAGATTGGCTGGCGTCGTGAGGGCATCAAGTACCGGCGCAAC
GAGCTTTTCCTGGACGTATTGGAGTACGTGAACCTGCTGATGAGCCCGCAGGGTCAGGTTTTGTCTGCTCACG
TGGCCGGCAAGGTGGTAATGAAGTCGTATTTGTCGGGTAAGTTAGCATAAATAATCTAGACATTATTCCTTTTAA
TAATCGCCATGTTGTAGGCATGCCCGAGTGCAAGTTCGGCATTAACGACAAGATCGTGATGGAGTCTAAGGGA
CGCGGTCTCTCCGGAAATTCAGAGGCGGAAACCTCACGCTCCGGCAAGCCCGTCGTGGTCATCGATGACTGC
CAGTTCCATCAGTGCGTCAAGCTAAGCAAATTCGAGACGGAGCATTCGATCAGCTTTATCCCGCCGGACGGG
GAGTTCGAGCTGATGCGTTACCGTACCACCAAAGACATTTCGCTGCCATTCCGAGTCATCCCGCTGGTGCGGG
AGGTGGGCCGCACCAAGGGGGAGAGAGTTATAAAA

+8862-2651-1809
13F.-12, No. 93, Sec. 1, Xintai 5th Rd., Xizhi Dist., New Taipei City 221416, Taiwan (R.O.C)
wellgenetics.com

**Analysis of Sequencing Results**:

Three independent sequencing reads were obtained from 3 independent genomic PCR of injection strain. Guide RNA 2 sequence can be found in three independent sequencing reads and each nucleotide showed in single peak in chromatographic view, suggesting that it is an identical read from both chromosome copies. **These results suggested that the target sequence of guide RNA 2 is present in the genome of injection strain.**

Chromatographic views of gRNA 2:

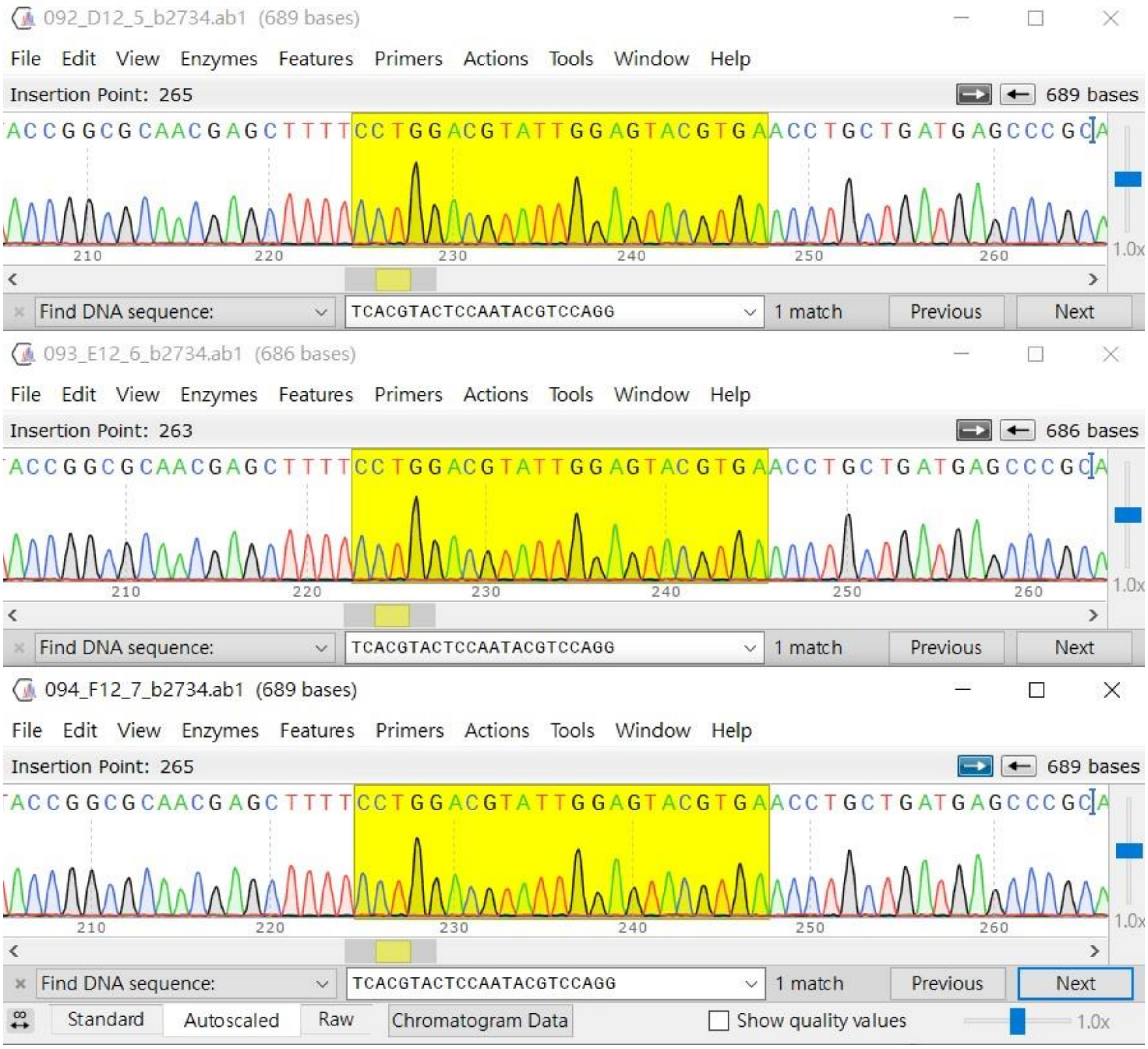

+8862-2651-1809
13F.-12, No. 93, Sec. 1, Xintai 5th Rd., Xizhi Dist., New Taipei City 221416, Taiwan (R.O.C)
wellgenetics.com

However, guide RNA 1 sequence cannot be found in three independent sequencing reads. We then analyze it by blat (see below). Guide RNA 1 locus is as indicated. One SNP is found on target sequence of guide RNA 1 in three independent reads in the injection strain. We search for another gRNA nearby instead shown as gRNA3 (next session).

Blat view of gRNA1:

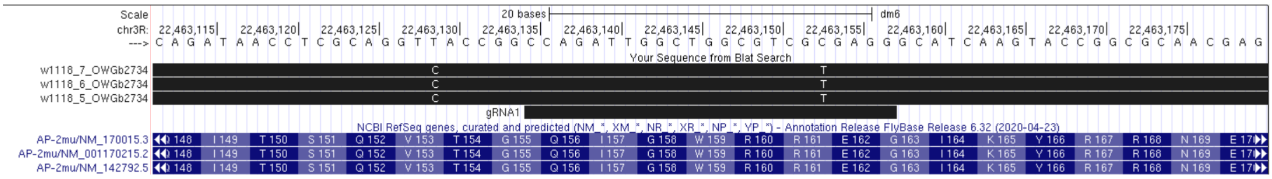

<u>**gRNA 3**</u>

**CRISPR Target Site [PAM]:** GACCCTGCGGGCTCATCAGC[AGG]

**CRISPR Target Strand in the Genome:** minus

**Cutting Site:** +613 nt from ATG of *AP-2mu*

**Distance:** +55 nt from upstream breakpoint to cutting site of Cas9

+41 nt from downstream breakpoint to cutting site of Cas9

**GC Content:** 70%

**T number at 17th to 20th Nucleotides:** 0

**Off target:** 0

**Guide RNA Primers:**

Sense oligo 5’- CTTCGACCCTGCGGGCTCATCAGC

Antisense oligo 5’- AAACGCTGATGAGCCCGCAGGGTC

**PAM mutation:** GACCCTGCGGGCTCATCAGC[AGA]

AAC to AAT, N to N

**Upstream Homology Arm:** 1,076bp, the -518 nt to +558 nt from ATG of *AP-2mu*

Forward Oligo 5’- ACCAGAGCGATGTCCATCAAG

Reverse Oligo 5’- TGCCCTCGCGACGCCAG

**Downstream Homology Arm:** 1,074bp, the +563 nt to +1,636 nt from ATG of *AP-2mu*

Synthesis fragment:

5’- GTACCGGTGGAACGAGCTTTTTCTGGACGTATTGGAGTACGTGAATCTGCTGATGAGCCC

Forward Oligo: 5’- GCAGGGTCAGGTTTTGTCTG

Reverse Oligo: 5’- CGGTATACGGCATAGGGTGT

+8862-2651-1809
13F.-12, No. 93, Sec. 1, Xintai 5th Rd., Xizhi Dist., New Taipei City 221416, Taiwan (R.O.C)
wellgenetics.com

**Analysis of Sequencing Results:**

Guide RNA 3 sequence can be found in three independent sequencing reads and each nucleotide showed in single peak in chromatographic view, suggesting that it is an identical read from both chromosome copies. **We will use guide RNA 2 in the first round microinjection and guide RNA 3 is the backup.**

Chromatographic views of gRNA3:

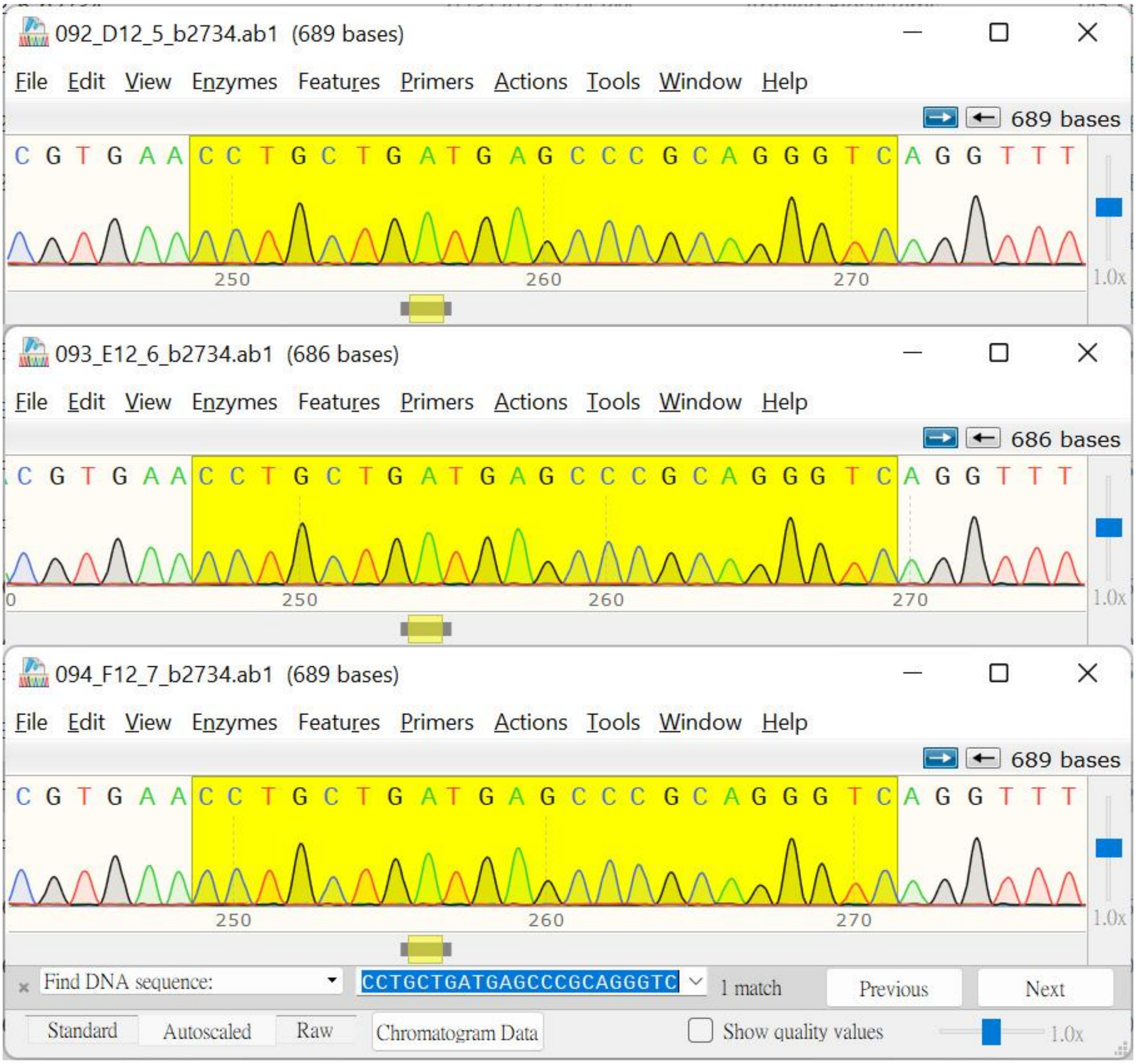

+8862-2651-1809

13F.-12, No. 93, Sec. 1, Xintai 5th Rd., Xizhi Dist., New Taipei City 221416, Taiwan (R.O.C)

wellgenetics.com

**Methods:**

Genomic DNA was obtained from genomic DNA of injection strain. PCR was performed using KOD_FX (TOYOBO) on BioRad S1000 Thermal Cycler. Sequencing results were shown in Snap Gene® viewer and gRNA sequences were found and highlighted in yellow or blue. If guide RNA sequences were not found in the sequencing results, sequencing results were further Blat[1] against *Drosophila melanogaster* genome (Aug 2014 Assembly, BDGP Release 6) using UCSC Genome Bioinformatics[2].

[1] Kent WJ. BLAT - the BLAST-like alignment tool. Genome Res. 2002 Apr;12(4):656-64.

[2] Kent WJ, Sugnet CW, Furey TS, Roskin KM, Pringle TH, Zahler AM, Haussler D. The human genome browser at UCSC. Genome Res. 2002 Jun;12(6):996-1006.

+8862-2651-1809
13F.-12, No. 93, Sec. 1, Xintai 5th Rd., Xizhi Dist., New Taipei City 221416, Taiwan (R.O.C)
wellgenetics.com

**Donor Design**

**Annotation**

NNNNN: Homology Arm / Coding Region

NNNNN: Homology Arm / UTR

nnnnn: Homology Arm/ Intron

nnnnn: Homology Arm/ Intergenic Region

nnnnn: gRNA

**nnnnnn**: Cloning Site

N: Silent mutation

N: Point mutation

nnnn: TTAA

nnnnn: Piggy Bac terminal repeats

nnnnn: 3xP3-hsp70 promoter

NNNNN: DsRed2

nnnnn: SV40 polyA

N: SNP

**Sequence**

accagagcgatgtccatcaagatagcgacgaaattagaacagtgcaattgccaattgggaatttgtattttaatttatttttaaattctgaaag
taatttaaatttaaaaaaaaaacttgagagctgtctagaaaagaactgatgtttcatgataactttgtcgaagaattaagaaatatttagttgt
aaaataattgttgaatctatttttttccaataacacgacttatatatttttgaaaatattcgagctaaatcccaagaagtaaactcaatctggg
atttgaagtgcccagaactcgaataaacacttctttttaaataattgtaagaccgtatcacttatggtatatactgacctcgaagggCCACAC
TAAGGGGGAGTGAAAATTGATTTTCTGATAAAAATTTTCGCTTGAAGCTACAGCATCGTCCACTGTCCATgta
tatatcttatatttgcatataaatatatatattacaccgacttggactaaccatcagATAGCGCACAAGATGATTGGCGGCCTGTT
CGTCTACAACCACAAGGGCGAGGTGCTGATCTCGCGAGTTTACCGCGACGACATCGGTCGGAATGCCGTG
GACGCCTTTCGGGTCAACGTCATCCACGCCCGCCAGCAGGTCCGCTCGCCAGTGACCAATATTGCGAGGAC
CAGCTTCTTCCACATCAAGgtcggtgaacaaagttcctatgtcacaatcaccaaccctcccacttatcatcaaaactcctttcacagA
GAGCAAACATTTGGCTGGCGGCTGTGACCAAGCAGAATGTGAACGCCGCGATGGTGTTTGAGTTCCTTTTG
AAGATCATCGAGGTGATGCAATCCTACTTCGGCAAGATCTCGGAGGAGAACATCAAGAATAACTTCGTGCT
CATCTACGAGCTGCTGGATGAGATCCTCGACTTTGGCTACCCGCAGAACACGGACTCCGGCACCCTGAAGA
CCTTCATCACACAGCAGGGCATCAAGTCGGCCACCAAGGAGGAGCAGATGCAGATAACCTCGCAGGTTACC
GGCCAGATTGGCTGGCGTCGCGAGGGCA
TTAAccctagaaagataatcatattgtgacgtacgttaaagataatcatgcgtaaaattgacgcatgtgttttatcggtctgtatatcgaggttt
atttattaatttgaatagatattaagttttattatatttacacttacatactaataataaattcaacaaacaatttatttatgtttatttatttattaaaa
aaaaacaaaaactcaaaatttcttctataaagtaacaaaactttaggatctaattcaattagagactaattcaattagagctaattcaattagg

+8862-2651-1809
13F.-12, No. 93, Sec. 1, Xintai 5th Rd., Xizhi Dist., New Taipei City 221416, Taiwan (R.O.C)
wellgenetics.com

atccaagcttatcgatttcgaaccctcgaccgccggagtataaatagaggcgcttcgtctacggagcgacaattcaattcaaacaagcaaagtg
aacacgtcgctaagcgaaagctaagcaaataaacaagcgcagctgaacaagctaaacaatcggctcgaagccggtcgccaccATGGCCT
CCTCCGAGGACGTCATCAAGGAGTTCATGCGCTTCAAGGTGCGCATGGAGGGCTCCGTGAACGGCCACGAG
TTCGAGATCGAGGGCGAGGGCGAGGGCCGCCCCTACGAGGGCACCCAGACCGCCAAGCTGAAGGTGACCA
AGGGCGGCCCCCTGCCCTTCGCCTGGGACATCCTGTCCCCCCAGTTCCAGTACGGCTCCAAGGTGTACGTGA
AGCACCCCGCCGACATCCCCGACTACAAGAAGCTGTCCTTCCCCGAGGGCTTCAAGTGGGAGCGCGTGATGA
ACTTCGAGGACGGCGGCGTGGTGACCGTGACCCAGGACTCCTCCCTCCAGGACGGCTCCTTCATCTACAAGG
TGAAGTTCATCGGCGTGAACTTCCCCTCCGACGGCCCCGTAATGCAGAAGAAGACTATGGGCTGGGAGGCGT
CCACCGAGCGCCTGTACCCCCGCGACGGCGTGCTGAAGGGCGAGATCCACAAGGCCCTGAAGCTGAAGGAC
GGCGGCCACTACCTGGTGGAGTTCAAGTCCATCTACATGGCCAAGAAGCCCGTGCAGCTGCCCGGCTACTACT
ACGTGGACTCCAAGCTGGACATCACCTCCCACAACGAGGACTACACCATCGTGGAGCAGTACGAGCGCGCCG
AGGGCCGCCACCACCTGTTCCTGTAGcggccgcgactctagatcataatcagccataccacatttgtagaggttttacttgctttaaa
aaacctcccacacctccccctgaacctgaaacataaaatgaatgcaattgttgttgttaacttgtttattgcagcttataatggttacaaataaag
caatagcatcacaaatttcacaaataaagcatttttttcactgcattctagttgtggtttgtccaaactcatcaatgtatcttagatatctataacaa
gaaaatatatatataataagttatcacgtaagtagaacatgaaataacaatataattatcgtatgagttaaatcttaaaagtcacgtaaaagat
aatcatgcgtcattttgactcacgcggtcgttatagttcaaaatcagtgacacttaccgcattgacaagcacgcctcacgggagctccaagcggc
gactgagatgtcctaaatgcacagcgacggattcgcgctatttagaaagagagagcaatatttcaagaatgcatgcgtcaattttacgcagact
atctttctagggttaa
GTACCGGTGGAACGAGCTTTTTCTGGACGTATTGGAGTACGTGAATCTGCTGATGAGCCC
GCAGGGTCAGGTTTTGTCTGCCCACGTGGCCGGCAAGGTGGTAATGAAGTCGTATTTGTCGGgtaagttagcat
aaataatctagacattattccttttaataatcgccatgttgtagGCATGCCCGAGTGCAAGTTCGGGATTAACGACAAGATC
GTGATGGAGTCTAAGGGACGCGGTCTCTCCGGAAATTCAGAGGCGGAAACCTCACGCTCCGGCAAGCCCG
TCGTGGTCATCGATGACTGCCAGTTCCATCAGTGCGTCAAGCTAAGCAAATTCGAGACGGAGCATTCGATC
AGCTTCATCCCGCCGGACGGGGAGTTCGAGCTGATGCGTTACCGTACCACCAAAGACATTTCGCTGCCATTC
CGAGTCATCCCGCTGGTGCGGGAGGTGGGCCGCACCAAGATGGAGGTTAAGGTTGTGCTGAAGTCCAACT
TTAAGCCCTCACTGCTGGGCCAAAAGATCGAGGTGAAGATACCAACCCCGCTCAATACATCGGGCGTGCAG
CTCATCTGCCTAAAGGGCAAAGCCAAATATAAGGCTTCGGAGAACGCGATCGTGTGGAAGATTAAGCGCAT
GGCGGGCATGAAGGAGACACAGCTGTCCGCGGAAATCGAACTTTTGGAGACGGACACCAAGAAGAAGTG
GACTCGGCCGCCCATCTCCATGAACTTTGAGGTGCCATTCGCGCCGTCCGGCTTCAAGGTACGCTACCTGAA
GGTGTTCGAGCCCAAGCTCAACTACTCCGACCACGATGTGGTCAAATGGGTGCGCTACATCGGACGCAGTG
GCTTGTATGAGACGCGCTGCTAGGGCGCCCAAGCATCCCCATCACGGAGCAAGATTATCACAATTGAATTT
AACGAGATGAATGGAGAAAGTGTTACCGCATCTATTGGTATTTCCTCAATCAAATTGAACCAGCAGCTTATG
TATTCCAAGATTAGTTCCAGCAAGTCCGGAGGACGCCCGAGTAGCGGATCCACCCCGGACACCCTATGCCG
TATACCG

☎ +8862-2651-1809 ✉
13F.-12, No. 93, Sec. 1, Xintai 5th Rd., Xizhi Dist., New Taipei City 221416, Taiwan (R.O.C)
wellgenetics.com

**Translation of wildtype**

NNNNN: the coding exon2 of *AP-2mu*

nnnnn: gRNA

N: expecting silent mutation

NNNN: expecting TTAA

N: expecting point mutation

N: SNP

**Nucleotides:**

AGAGCAAACATTTGGCTGGCGGCTGTGACCAAGCAGAATGTGAACGCCGCGATGGTGTTTGAGTTCCTTTT
GAAGATCATCGAGGTGATGCAATCCTACTTCGGCAAGATCTCGGAGGAGAACATCAAGAATAACTTCGTGC
TCATCTACGAGCTGCTGGATGAGATCCTCGACTTTGGCTACCCGCAGAACACGGACTCCGGCACCCTGAAG
ACCTTCATCACACAGCAGGGCATCAAGTCGGCCACCAAGGAGGAGCAGATGCAGATAACCTCGCAGGTTAC
CGGCCAGATTGGCTGGCGTCGCGAGGGCATCAAGTACCGGCGCAACGAGCTTTTCCTGGACGTATTGGAG
TACGTGAACCTGCTGATGAGCCCGCAGGGTCAGGTTTTGTCTGCCCACGTGGCCGGCAAGGTGGTAATGAA
GTCGTATTTGTCGG

**Protein:**

R A N I W L A A V T K Q N V N A A Met V F E F L L K I I E V Met Q S Y F G K I S E E N I K N N F V L I Y E
L L D E I L D F G Y P Q N T D S G T L K T F I T Q Q G I K S A T K E E Q Met Q I T S Q V T G Q I G W R R E
G I K Y R R N E L F L D V L E Y V N L L Met S P Q G Q V L S A H V A G K V V Met K S Y L S

**Translation of *AP-2mu R168W* after excision**

NNNNN: the coding exon2 of *AP-2mu R168W*

nnnnn: gRNA

N: Silent mutation

NNNN: TTAA

N: Point mutation

N: SNP

**Nucleotides:**

AGAGCAAACATTTGGCTGGCGGCTGTGACCAAGCAGAATGTGAACGCCGCGATGGTGTTTGAGTTCCTTTT
GAAGATCATCGAGGTGATGCAATCCTACTTCGGCAAGATCTCGGAGGAGAACATCAAGAATAACTTCGTGC
TCATCTACGAGCTGCTGGATGAGATCCTCGACTTTGGCTACCCGCAGAACACGGACTCCGGCACCCTGAAG

+8862-2651-1809
13F.-12, No. 93, Sec. 1, Xintai 5th Rd., Xizhi Dist., New Taipei City 221416, Taiwan (R.O.C)
wellgenetics.com

ACCTTCATCACACAGCAGGGCATCAAGTCGGCCACCAAGGAGGAGCAGATGCAGATAACCTCGCAGGTTAC
CGGCCAGATTGGCTGGCGTCGCGAGGGCATTAAGTACCGGTGGAACGAGCTTTTTCTGGACGTATTGGAG
TACGTGAATCTGCTGATGAGCCCGCAGGGTCAGGTTTTGTCTGCCCACGTGGCCGGCAAGGTGGTAATGAA
GTCGTATTTGTCGG

**Protein:**

R A N I W L A A V T K Q N V N A A Met V F E F L L K I I E V Met Q S Y F G K I S E E N I K N N F V L I Y E
L L D E I L D F G Y P Q N T D S G T L K T F I T Q Q G I K S A T K E E Q Met Q I T S Q V T G Q I G W R R E
G I K Y R W N E L F L D V L E Y V N L L Met S P Q G Q V L S A H V A G K V V Met K S Y L S

+8862-2651-1809
13F.-12, No. 93, Sec. 1, Xintai 5th Rd., Xizhi Dist., New Taipei City 221416, Taiwan (R.O.C)
wellgenetics.com

WELLGENETICS

### CRISPR Cloning Report

| | | | |
|---|---|---|---|
| **Report No.:** | RWGa3634 | **Date:** | 2023.03.15 |
| **Reporter:** | Chien-Hsiang Wang | **E-mail:** | |

*************************************************************************

**Gene:** *Ap-2mu/CG7057* **Case No.:** 220862

**Project:** introducing a point mutation R168W of AP-2mu using PBac system to facilitate genetic screening

**Method:** CRISPR/Cas9-mediated genome editing by homology-dependent repair (HDR) using 1 guide RNA(s) and a dsDNA plasmid donor

**Clone:** *PWG8438* *pUC57-Kan-220862 donor*
Kanamycin Resistant

☎ +8862-2651-1809 ✉
13F.-12, No. 93, Sec. 1, Xintai 5th Rd., Xizhi Dist., New Taipei City 221416, Taiwan (R.O.C)
wellgenetics.com

#### Gene and Method

**Gene**: *Ap-2mu/CG7057*

**Location:** 3R (94A15-94A16)

**Method:** CRISPR/Cas9-mediated genome editing by homology-dependent repair (HDR) using a guide RNA and a dsDNA plasmid donor

**Knock-in cassette:** PBacDsRed

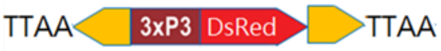

**Injection strain:** *w[1118]*

**Genome Editing:**

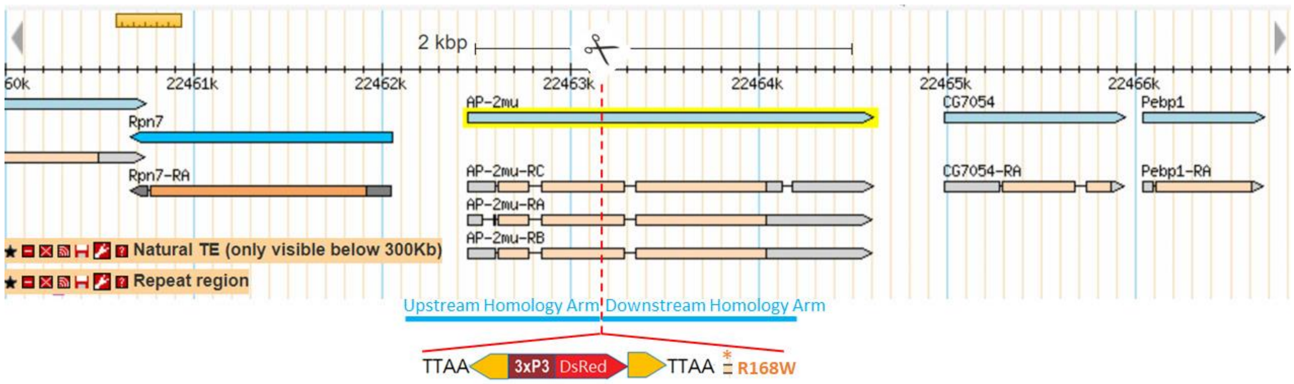

(1) Introduce a point mutation R168 to W, CGC to TGG of *AP-2mu*.

(2) The selection marker PBacDsRed contains 3' PBac terminal repeats, the artificial 3xP3 promoter (three tandem copies of the Pax-6 homodimer binding site and TATA-homology of *hsp70*), DsRed2, SV40 3'UTR, and 5' PBac terminal repeats. It facilitates the genetic screening and can be excised by Piggy Bac transposase. Only one TTAA motif will be left after transposition embedded in mutated exon sequence, and create a silent mutation on I164 of AP-2mu; I164K165 ATCAAG to ATTAAG, IK to IK.

(3) Selection marker needs to be excised when looking at endogenous gene/protein expression.

(4) PAM mutations to make the donor inactive to guide RNA will be incorporated into the edited genome.

☎ +8862-2651-1809 ✉
13F.-12, No. 93, Sec. 1, Xintai 5th Rd., Xizhi Dist., New Taipei City 221416, Taiwan (R.O.C)
wellgenetics.com

WellGenetics

Donor Plasmid Information

**Vector:** pUC57-Kan

**Vector map:**

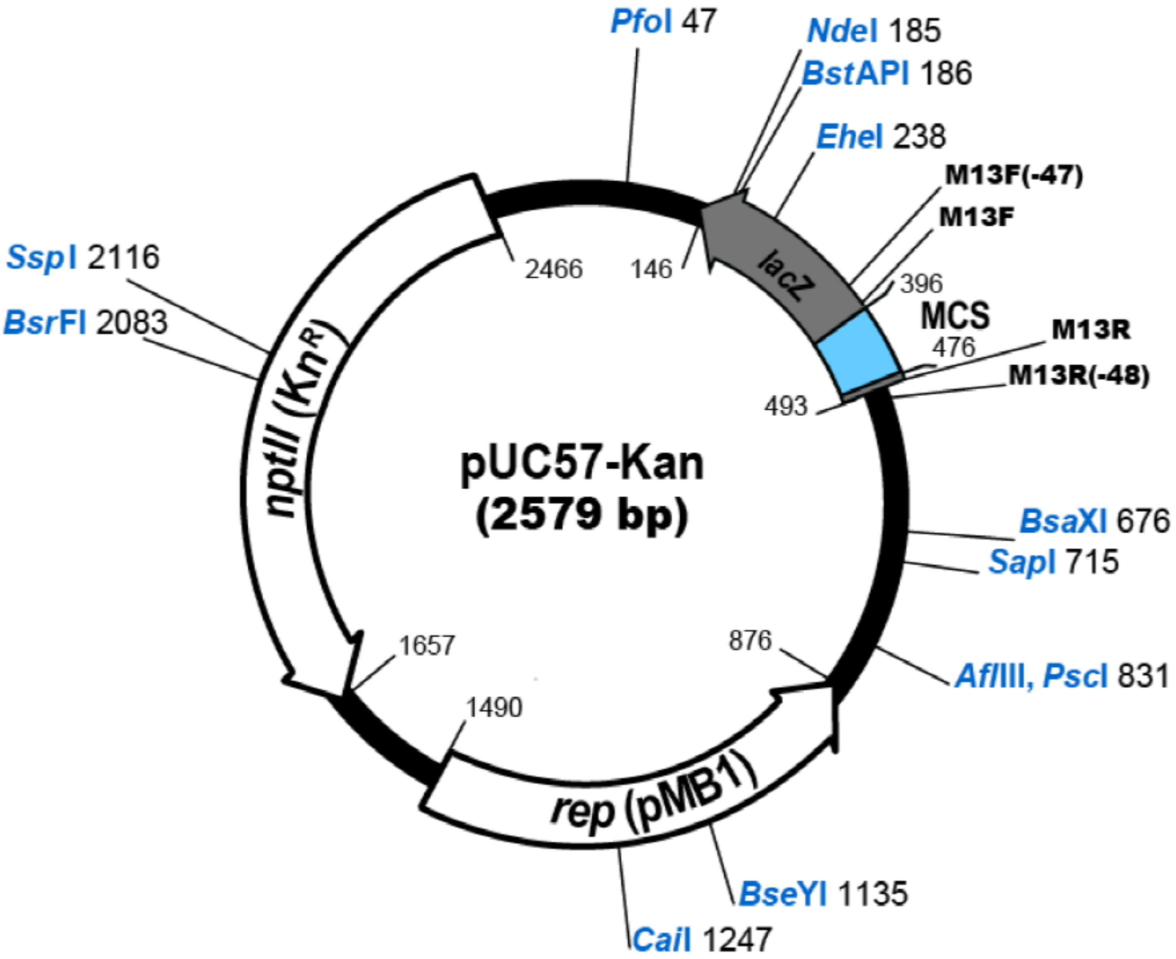

**PCR amplification template:** $w^{1118}$

**Annotation**

*nnnnn*: vector backbone

NNNNN: Homology Arm / Coding Region

NNNNN: Homology Arm / UTR

nnnnn: Homology Arm/ Intron

nnnnn: Homology Arm/ Intergenic Region

nnnnn: gRNA

**nnnnnn**: Cloning Site

N: Silent mutation

+8862-2651-1809
13F.-12, No. 93, Sec. 1, Xintai 5th Rd., Xizhi Dist., New Taipei City 221416, Taiwan (R.O.C)
wellgenetics.com

N: Point mutation
nnnn: TTAA
nnnnn: Piggy Bac terminal repeats
nnnnn: 3xP3-hsp70 promoter
NNNNN: DsRed2
nnnnn: SV40 polyA

**Sequence**

*tcgcgcgtttcggtgatgacggtgaaaacctctgacacatgcagctcccggagacggtcacagcttgtctgtaagcggatgccgggag*
*cagacaagcccgtcagggcgcgtcagcgggtgttggcgggtgtcggggctggcttaactatgcggcatcagagcagattgtactgag*
*agtgcaccatatgcggtgtgaaataccgcacagatgcgtaaggagaaaataccgcatcaggcgccattcgccattcaggctgcgcaa*
*ctgttgggaagggcgatcggtgcgggcctcttcgctattacgccagctggcgaaagggggatgtgctgcaaggcgattaagttgggta*
*acgccagggttttcccagtcacgacgttgtaaaacgacggcca**gagaattc**gcggccgc**tctaga***accagagcgatgtccatcaagata
gcgacgaaattagaacagtgcaattgccaattgggaatttgtattttaatttatttttaaattctgaaagtaatttaaatttaaaaaaaaactt
gagagctgtctagaaaagaactgatgtttcatgataactttgtcgaagaattaagaaatatttagttgtaaaataattgttgaatctatttttttc
caataacacgacttatatatttttgaaaatattcgagctaaatcccaagaagtaaactcaatctgggatttgaagtgcccagaactcgaata
aacacttctttttaaataattgtaagaccgtatcacttatggtatatactgacctcgaagggCCACACTAAGGGGGAGTGAAAATT
GATTTTCTGATAAAAATTTTCGCTTGAAGCTACAGCATCGTCCACTGTCCATgtatatatcttatatttgcatataaatat
atatattacaccgacttggactaaccatcagATAGCGCACAAGATGATTGGCGGCCTGTTCGTCTACAACCACAAGGG
CGAGGTGCTGATCTCGCGAGTTTACCGCGACGACATCGGTCGGAATGCCGTGGACGCCTTTCGGGTCAACG
TCATCCACGCCCGCCAGCAGGTCCGCTCGCCAGTGACCAATATTGCGAGGACCAGCTTCTTCCACATCAAGg
tcggtgaacaaagttcctatgtcacaatcaccaaccctcccacttatcatcaaaactcctttcacagAGAGCAAACATTTGGCTGGC
GGCTGTGACCAAGCAGAATGTGAACGCCGCGATGGTGTTTGAGTTCCTTTTGAAGATCATCGAGGTGATGC
AATCCTACTTCGGCAAGATCTCGGAGGAGAACATCAAGAATAACTTCGTGCTCATCTACGAGCTGCTGGAT
GAGATCCTCGACTTTGGCTACCCGCAGAACACGGACTCCGGCACCCTGAAGACCTTCATCACACAGCAGGG
CATCAAGTCGGCCACCAAGGAGGAGCAGATGCAGATAACCTCGCAGGTTACCGGCCAGATTGGCTGGCGT
CGCGAGGGCATTAAccctagaaagataatcatattgtgacgtacgttaaagataatcatgcgtaaaattgacgcatgtgttttatcggtct
gtatatcgaggtttatttattaatttgaatagatattaagttttattatatttacacttacatactaataataaattcaacaaacaatttatttatgttt
atttatttattaaaaaaaaacaaaaactcaaaatttcttctataaagtaacaaaactttta ggatctaattcaattagagactaattcaattagag
ctaattcaattaggatccaagcttatcgatttcgaaccctcgaccgccggagtataaatagaggcgcttcgtctacggagcgacaattcaattca
aacaagcaaagtgaacacgtcgctaagcgaaagctaagcaaataaacaagcgcagctgaacaagctaaacaatcggctcgaagccggtcg
ccaccATGGCCTCCTCCGAGGACGTCATCAAGGAGTTCATGCGCTTCAAGGTGCGCATGGAGGGCTCCGTGAA
CGGCCACGAGTTCGAGATCGAGGGCGAGGGCGAGGGCCGCCCCTACGAGGGCACCCAGACCGCCAAGCTG
AAGGTGACCAAGGGCGGCCCCCTGCCCTTCGCCTGGGACATCCTGTCCCCCCAGTTCCAGTACGGCTCCAAG

+8862-2651-1809
13F.-12, No. 93, Sec. 1, Xintai 5th Rd., Xizhi Dist., New Taipei City 221416, Taiwan (R.O.C)
wellgenetics.com

GTGTACGTGAAGCACCCCGCCGACATCCCCGACTACAAGAAGCTGTCCTTCCCCGAGGGCTTCAAGTGGGAG
CGCGTGATGAACTTCGAGGACGGCGGCGTGGTGACCGTGACCCAGGACTCCTCCCTCCAGGACGGCTCCTTC
ATCTACAAGGTGAAGTTCATCGGCGTGAACTTCCCCTCCGACGGCCCCGTAATGCAGAAGAAGACTATGGGCT
GGGAGGCGTCCACCGAGCGCCTGTACCCCCGCGACGGCGTGCTGAAGGGCGAGATCCACAAGGCCCTGAA
GCTGAAGGACGGCGGCCACTACCTGGTGGAGTTCAAGTCCATCTACATGGCCAAGAAGCCCGTGCAGCTGCC
CGGCTACTACTACGTGGACTCCAAGCTGGACATCACCTCCCACAACGAGGACTACACCATCGTGGAGCAGTAC
GAGCGCGCCGAGGGCCGCCACCACCTGTTCCTGTAGcggccgcgactctagatcataatcagccataccacatttgtagaggt
tttacttgctttaaaaaacctcccacacctccccctgaacctgaaacataaaatgaatgcaattgttgttgttaacttgtttattgcagcttataat
ggttacaaataaagcaatagcatcacaaatttcacaaataaagcatttttttcactgcattctagttgtggtttgtccaaactcatcaatgtatctt
agatatctataacaagaaaatatatatataataagttatcacgtaagtagaacatgaaataacaatataattatcgtatgagttaaatcttaaaa
gtcacgtaaaagataatcatgcgtcattttgactcacgcggtcgttatagttcaaaatcagtgacacttaccgcattgacaagcacgcctcacgg
gagctccaagcggcgactgagatgtcctaaatgcacagcgacggattcgcgctatttagaaagagagagcaatatttcaagaatgcatgcgtc
aattttacgcagactatctttctagggttaaGTACCGGTGGAACGAGCTTTTTCTGGACGTATTGGAGTACGTGAATCTG
CTGATGAGCCCGCAGGGTCAGGTTTTGTCTGCCCACGTGGCCGGCAAGGTGGTAATGAAGTCGTATTTGTC
GGgtaagttagcataaataatctagacattattccttttaataatcgccatgttgtagGCATGCCCGAGTGCAAGTTCGGGATTA
ACGACAAGATCGTGATGGAGTCTAAGGGACGCGGTCTCTCCGGAAATTCAGAGGCGGAAACCTCACGCTC
CGGCAAGCCCGTCGTGGTCATCGATGACTGCCAGTTCCATCAGTGCGTCAAGCTAAGCAAATTCGAGACGG
AGCATTCGATCAGCTTCATCCCGCCGGACGGGGAGTTCGAGCTGATGCGTTACCGTACCACCAAAGACATT
TCGCTGCCATTCCGAGTCATCCCGCTGGTGCGGGAGGTGGGCCGCACCAAGATGGAGGTTAAGGTTGTGC
TGAAGTCCAACTTTAAGCCCTCACTGCTGGGCCAAAAGATCGAGGTGAAGATACCAACCCCGCTCAATACA
TCGGGCGTGCAGCTCATCTGCCTAAAGGGCAAAGCCAAATATAAGGCTTCGGAGAACGCGATCGTGTGGA
AGATTAAGCGCATGGCGGGCATGAAGGAGACACAGCTGTCCGCGGAAATCGAACTTTTGGAGACGGACAC
CAAGAAGAAGTGGACTCGGCCGCCCATCTCCATGAACTTTGAGGTGCCATTCGCGCCGTCCGGCTTCAAGG
TACGCTACCTGAAGGTGTTCGAGCCCAAGCTCAACTACTCCGACCACGATGTGGTCAAATGGGTGCGCTAC
ATCGGACGCAGTGGCTTGTATGAGACGCGCTGCTAGGGCGCCCAAGCATCCCCATCACGGAGCAAGATTAT
CACAATTGAATTTAACGAGATGAATGGAGAAAGTGTTACCGCATCTATTGGTATTTCCTCAATCAAATTGAA
CCAGCAGCTTATGTATTCCAAGATTAGTTCCAGCAAGTCCGGAGGACGCCCGAGTAGCGGATCCACCCCGG
ACACCCTATGCCGTATACCG***ggcgcgccaagctt****ggtgtaatcatggtcatagctgtttcctgtgtgaaattgttatccgctcaca*
*attccacacaacatacgagccggaagcataaagtgtaaagcctggggtgcctaatgagtgagctaactcacattaattgcgttgcgctc*
*actgcccgctttccagtcgggaaacctgtcgtgccagctgcattaatgaatcggccaacgcgcggggagaggcggtttgcgtattgggc*
*gctcttccgcttcctcgctcactgactcgctgcgctcggtcgttcggctgcggcgagcggtatcagctcactcaaaggcggtaatacggtt*
*atccacagaatcaggggataacgcaggaaagaacatgtgagcaaaaggccagcaaaaggccaggaaccgtaaaaaggccgcgtt*
*gctggcgtttttccataggctccgcccccctgacgagcatcacaaaaatcgacgctcaagtcagaggtggcgaaacccgacaggactat*
*aaagataccaggcgtttccccctggaagctccctcgtgcgctctcctgttccgaccctgccgcttaccggatacctgtccgcctttctcccttc*
*gggaagcgtggcgctttctcatagctcacgctgtaggtatctcagttcggtgtaggtcgttcgctccaagctgggctgtgtgcacgaaccc*

+8862-2651-1809
13F.-12, No. 93, Sec. 1, Xintai 5th Rd., Xizhi Dist., New Taipei City 221416, Taiwan (R.O.C)
wellgenetics.com

*cccgttcagcccgaccgctgcgccttatccggtaactatcgtcttgagtccaacccggtaagacacgacttatcgccactggcagcagcc*
*actggtaacaggattagcagagcgaggtatgtaggcggtgctacagagttcttgaagtggtggcctaactacggctacactagaaga*
*acagtatttggtatctgcgctctgctgaagccagttaccttcggaaaaagagttggtagctcttgatccggcaaacaaaccaccgctggt*
*agcggtggtttttttgtttgcaagcagcagattacgcgcagaaaaaaaggatctcaagaagatcctttgatcttttctacggggtctgac*
*gctcagtggaacgaaaactcacgttaagggattttggtcatgagattatcaaaaaggatcttcacctagatccttttaaattaaaaatg*
*aagttttaaatcaagcccaatctgaataatgttacaaccaattaaccaattctgattagaaaaactcatcgagcatcaaatgaaactgc*
*aatttattcatatcaggattatcaataccatatttttgaaaaagccgtttctgtaatgaaggagaaaactcaccgaggcagttccatagg*
*atggcaagatcctggtatcggtctgcgattccgactcgtccaacatcaatacaacctattaatttcccctcgtcaaaaataaggttatcaa*
*gtgagaaatcaccatgagtgacgactgaatccggtgagaatggcaaaagtttatgcatttctttccagacttgttcaacaggccagcca*
*ttacgctcgtcatcaaaatcactcgcatcaaccaaaccgttattcattcgtgattgcgcctgagcgagacgaaatacgcgatcgctgtta*
*aaaggacaattacaaacaggaatcgaatgcaaccggcgcaggaacactgccagcgcatcaacaatattttcacctgaatcaggatat*
*tcttctaatacctggaatgctgtttttccggggatcgcagtggtgagtaaccatgcatcatcaggagtacggataaaatgcttgatggtc*
*ggaagaggcataaattccgtcagccagtttagtctgaccatctcatctgtaacatcattggcaacgctacctttgccatgtttcagaaaca*
*actctggcgcatcgggcttcccatacaagcgatagattgtcgcacctgattgcccgacattatcgcgagcccatttatacccatataaatc*
*agcatccatgttggaatttaatcgcggcctcgacgtttcccgttgaatatggctcataacaccccttgtattactgtttatgtaagcagaca*
*gttttattgttcatgatgatatatttttatcttgtgcaatgtaacatcagagattttgagacacgggccagagctgca*

+8862-2651-1809
13F.-12, No. 93, Sec. 1, Xintai 5th Rd., Xizhi Dist., New Taipei City 221416, Taiwan (R.O.C)
wellgenetics.com

**Colony Sequencing Results:**

>220862ad1_OWG5476

TACGCTACTTCGCATTACGCCAGCTGGCGAAGGGGGATGTGCTGCAAGGCGATTAAGTTGGGTAACGCCAGG
GTTTTCCCAGTCACGACGTTGTAAAACGACGGCCAGAGAATTCGCGGCCGCTCTAGAACCAGAGCGATGTCC
ATCAAGATAGCGACGAAATTAGAACAGTGCAATTGCCAATTGGGAATTTGTATTTTAATTTATTTTTAAATTCTG
AAAGTAATTTTAATTTAAAAAAAACTTGAGAGCTGTCTAGAAAAGAACTGATGTTTCATGATAACTTTGTCGAA
GAATTAAGAAATATTTAGTTGTAAAATAATTGTTGAAGCTATTTTTTTCCAATAACACGACTTATATATTTTTTGA
AAATATTCGAGCTAAATCCCAAGAAGTAAACTCAATCTGGGATTTGAAGTGCCCAGAACTCGAATAAACACTT
CTTTTTAAATAATTGTAAGACCGTATCACTTATGGTATATACTGACCTCGAAGGGCCACACTAAGGGGGAGTGA
AAATTGATTTTCTGATAAAAATTTTCGCTTGAAGCTCCAGCATCGTCCACTGTCCATGTATATATCTTATATTTGCA
TATAAATATATATATTACACCGACTTGGACTAACCATCAGATAGCGCACAAGATGATTGGCGGCCTGTTCGTCTA
CAACCACAAGGGCGAGGTGCTGATCTCGCGAGTTTACCGCGACGACATCGGTCGGAATGCCGTGGACGCCTT
TCGGGTCAACGTCATCCACGCCCGCCAGCAGGTCCGCTCGCCAGTGACCAATATTGCGAGGACCAGCTTCTTC
CACATCAAGGTCGGTGAACAAAGTTCCTATGTCACAATCACCAACCCTCCCACTCATCATCAAAACTCCTTTCA
CAGAGAGCAAACATTTGGCTGGCGGCTGTGACCAAGCAGAATGTGAACGCCGCGATGGTGTTTGAGTTCCTT
TTGAAGATCATCGAGGTGATGCAATCCTACTTCGGCAGATCTCAGAGGAGACATCAGAATAACTTCGTGCTCAT
CTACGAGCTGCTGGATGAGATCCTCGACTTTGGCTACCCGCAGACACGGACTCCGGTACCCTGAGACTCATCA
CACAGCAGGCATCAGTCGCACAGGAGGAGCAAATGCAGAATAACCTCGCAGGTCACGTCGAATGGCTGGCG
TCGCGAGGGCATTAACTAAAAGATATCCATATTGTGGACGTACGTTAAAGAATATCATGCTAAATTGAGCCAAG
TGTTTACCGCTGTTAATTCCAGGTTATTATTAAATTTGGGAGAGACTG

>220862ad1_OWG0156

ACGTCACTGATTAGCTCTATTGATTAGTCTCTAATTGAATTAGATCCTAAAAGTTTTGTTACTTTATAGAAGAAAT
TTTGAGTTTTTGTTTTTTTTTAATAAATAAATAAACATAAATAAATTGTTTGTTGAATTTATTATTAGTATGTAAGT
GTAAATATAATAAAACTTAATATCTATTCAAATTAATAAATAAACCTCGATATACAGACCGATAAAACACATGCGT
CAATTTTACGCATGATTATCTTTAACGTACGTCACAATATGATTATCTTTCTAGGGTTAATGCCCTCGCGACGCCA
GCCAATCTGGCCGGTGACCTGCGAGGTTATCTGCATCTGCTCCTCCTTGGTGGCCGACTTGATGCCCTGCTGT
GTGATGAAGGTCTTCAGGGTACCGGAGTCCGTGTTCTGCGGGTAGCCAAAGTCGAGGATCTCATCCAGCAGC
TCGTAGATGAGCACGAAGTTATTCTTGATGTTCTCCTCTGAGATCTTGCCGAAGTAGGATTGCATCACCTCGAT
GATCTTCAAAAGGAACTCAAACACCATCGCGGCGTTCACATTCTGCTTGGTCACAGCCGCCAGCCAAATGTTT
GCTCTCTGTGAAAGGAGTTTTGATGATGAGTGGGAGGGTTGGTGATTGTGACATAGGAACTTTGTTCACCGA
CCTTGATGTGGAAGAAGCTGGTCCTCGCAATATTGGTCACTGGCGAGCGGACCTGCTGGCGGGCGTGGATGA
CGTTGACCCGAAAGGCGTCCACGGCATTCCGACCGATGTCGTCGCGGTAAACTCGCGAGATCAGCACCTCGC
CCTTGTGGTTGTAGACGAACAGGCCGCCAATCATCTTGTGCGCTATCTGATGGTTAGTCCAAGTCGGTGTAATA
TATATATTTATATGCAAATATAAGATATATACATGGACAGTGGACGATGCTGGAGCTTCAAGCGAAAATTTTTATC

+8862-2651-1809
13F.-12, No. 93, Sec. 1, Xintai 5th Rd., Xizhi Dist., New Taipei City 221416, Taiwan (R.O.C)
wellgenetics.com

AGAAAATCAATTTTCACTCCCCCTTAGTGTGCCCTTCGAGGTCAGTATATACCATAAGTGATACGGTCTTACATTA
TTTAAAAGAGTGTTTATTCGAGTTCTGGGCACTTCAAATCCCAGATGAGTTACTCTGGGATTTAGCTCGATATTT
CAAAAATATTATAAGTCGTGTTATTGAAAAAAATAGCTCACATATTTACACTTAATTTCTATCTCGACAAGTTATCT
GACTCAGTCTTTCCGAGACAGCCCTCAGTTTTTACTATAGAAATATACCTCTCGGAATTGA

>220862ad1_OWG2409
GGATGGTGACTTACCGCATTGACAGCACGCCTCACGGGAGCTCCAAGCGGCGACTGAGATGTCCTAAATGCA
CAGCGACGGATTCGCGCTATTTAGAAAGAGAGAGCAATATTTCAAGAATGCATGCGTCAATTTTACGCAGACT
ATCTTTCTAGGGTTAAGTACCGGTGGAACGAGCTTTTTCTGGACGTATTGGAGTACGTGAATCTGCTGATGAG
CCCGCAGGGTCAGGTTTTGTCTGCTCACGTGGCCGGCAAGGTGGTAATGAAGTCGTATTTGTCGGGTAAGTT
AGCATAAATAATCTAGACATTATTCCTTTTAATAATCGCCATGTTGTAGGCATGCCCGAGTGCAAGTTCGGCATT
AACGACAAGATCGTGATGGAGTCTAAGGGACGCGGTCTCTCCGGAAATTCAGAGGCGGAAACCTCACGCTCC
GGCAAGCCCGTCGTGGTCATCGATGACTGCCAGTTCCATCAGTGCGTCAAGCTAAGCAAATTCGAGACGGAG
CATTCGATCAGCTTTATCCCGCCGGACGGGGAGTTCGAGCTGATGCGTTACCGTACCACCAAAGACATTTCGC
TGCCATTCCGAGTCATCCCGCTGGTGCGGGAGGTGGGCCGCACCAAGATGGAGGTTAAGGTTGTGCTGAAG
TCCAACTTTAAGCCCTCACTGCTGGGCCAAAAGATCGAGGTGAAGATACCAACCCCGCTCAATACATCGGGCG
TGCAGCTCATCTGCCTAAAGGGCAAAGCCAAATACAAGGCTTCGGAGAACGCGATCGTGTGGAAGATTAAGC
GCATGGCGGGCATGAAGGAGACACAGCTGTCCGCGGAAATCGAGCTTTTGGAGACGGACACCAAGAAGAA
GTGGACTCGGCCGCCCATCTCCATGAACTTTGAGGTGCCATTCGCGCCGTCCGGCTTCAAGGTACGCTACCTG
AAAGTGTTCGAGCCCAAGCTCAACTACTCCGACCACGATGTGGTCAAATGGGTGCGCTACATCGGACGCAGT
GGCTTGTATGAGACGCGCTGCTAGGCGCCCAGCATCCCCATCACGGAGCAGATTATCACAATTGAATTTACGA
GATGATGAGAAAATGTACGCATCTTATGTATTCTCATCAAATGACAGCAGCTTATGTATTCAGATAGTCAGCAGT
CGGAGGACGCCCGGTTAGCGAATCCAACTCGACTCCCTATGCGTACGGGCGGCAGCTGGGGGTATCATGGGC
TAGCGGCTTCTCTGTGGTGGAATGTGTTTAATATTTG

>220862ad1_OWG5477
CCAGCTCTGCTCGTATGTTGTGTGGATTGTGAGCGGATAACAATTTCACACAGGAAACAGCTATGACCATGATT
ACACCAAGCTTGGCGCGCCCGGTATACGGCATAGGGTGTCCGGGGTGGATCCGCTACTCGGGCGTCCTCCGG
ACTTGCTGGAACTAATCTTGGAATACATAAGCTGCTGGTTCAATTTGATTGAGGAAATACCAATAGATGCGGTA
ACATTTTCTCCATTCATCTCGTTAAATTCAATTGTGATAATCTTGCTCCGTGATGGGGATGCTTGGGCGCCCTAG
CAGCGCGTCTCATACAAGCCACTGCGTCCGATGTAGCGCACCCATTTGACCACATCGTGGTCGGAGTAGTTGA
GCTTGGGCTCGAACACCTTCAGGTAGCGTACCTTGAAGCCGGACGGCGCGAATGGCACCTCAAAGTTCATGG
AGATGGGCGGCCGAGTCCACTTCTTCTTGGTGTCCGTCTCCAAAAGCTCGATTTCCGCGGACAGCTGTGTCTC
CTTCATGCCCGCCATGCGCTTAATCTTCCACACGATCGCGTTCTCCGAAGCCTTGTATTTGGCTTTGCCCTTTAG
GCAGATGAGCTGCACGCCCGATGTATTGAGCGGGGTTGGTATCTTCACCTCGATCTTTTGGCCCAGCAGTGAG

+8862-2651-1809
13F.-12, No. 93, Sec. 1, Xintai 5th Rd., Xizhi Dist., New Taipei City 221416, Taiwan (R.O.C)
wellgenetics.com

GGCTTAAAGTTGGACTTCAGCACAACCTTAACCTCCATCTTGGTGCGGCCCACCTCCCGCACCAGCGGGATGA
CTCGGAATGGCAGCGAAATGTCTTTGGTGGTACGGTAACGCATCAGCTCGAACTCCCCGTCCGGCGGGATAA
AGCTGATCGAATGCTCCGTCTCGAATTTGCTTAGCTTGACGCACTGATGGAACTGGCAGTCATCGATGACCAC
GACGGGCTTGCCGGAGCGTGAGGTTTCCGCCTCTGAATTTCCGGAGAGACCGCGTCCCTTAGACTCCATCAC
GATCTTGTCGTTAATGCCGAACTTGCACTCGGGCATGCCTACACATGGCGATATTAAAGGATATGTCTAGATTAT
TTATGCTAACTTACCGACAATACGACTCATACCACCTGCCGGCCACGTGAGCAGACAAACTGACCCTGCGGGC
TCATCAGCAGATCACGTACTCATACGTCAGAAAAGCTCGTTCACGTACTACCTAGAAGATAGTCTGGCGTAAAA
TTGACCATGCATTCTGGAAATTGTCCTCTCCATCATAGCGAATCCGTTGCGTGCCATAGAACTCTCGTCGCGCCT
GGGAGGATCTCTGT

**Blat Results and Conclusions:**

Both upstream and downstream homology arms in the donor plasmid were sequenced to be specific to *Ap-2mu/CG7057* (see overview below). Eight mismatches (boxed in blue) within the homology arm region were found in the CDS of *Ap-2mu/CG7057.* All mismatches (boxed in blue) that do not lead to amino acid change, are considered as silent SNPs. Orange box indicates the designed point mutation of *Ap-2mu/CG7057*.

Overview:

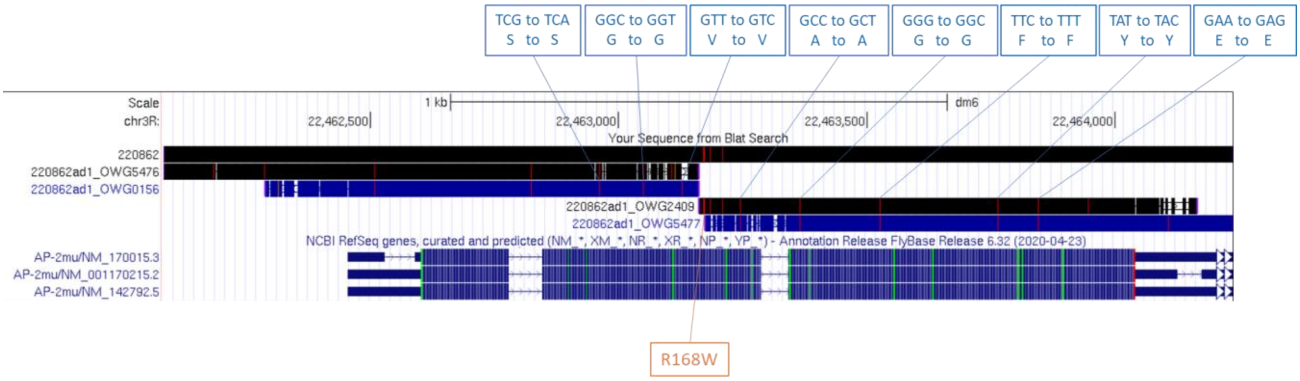

+8862-2651-1809
13F.-12, No. 93, Sec. 1, Xintai 5th Rd., Xizhi Dist., New Taipei City 221416, Taiwan (R.O.C)
wellgenetics.com

Sequence reads (>220862ad1_OWG0156 and >220862ad1_OWG2409) were aligned to 220862 donor using Blast to check the junctions of cassette PBacDsRed insertion. The sequences of PiggyBac (bracketed in purple) were correctly inserted in the coding exon (bracketed in blue) of *Ap-2mu/CG7057* as design. Designed point mutation was shaded in orange. **We will use clone 220862ad1 for microinjection.**

220862ad1_OWG0156 (Sbjct) vs 220862 donor design (Query)

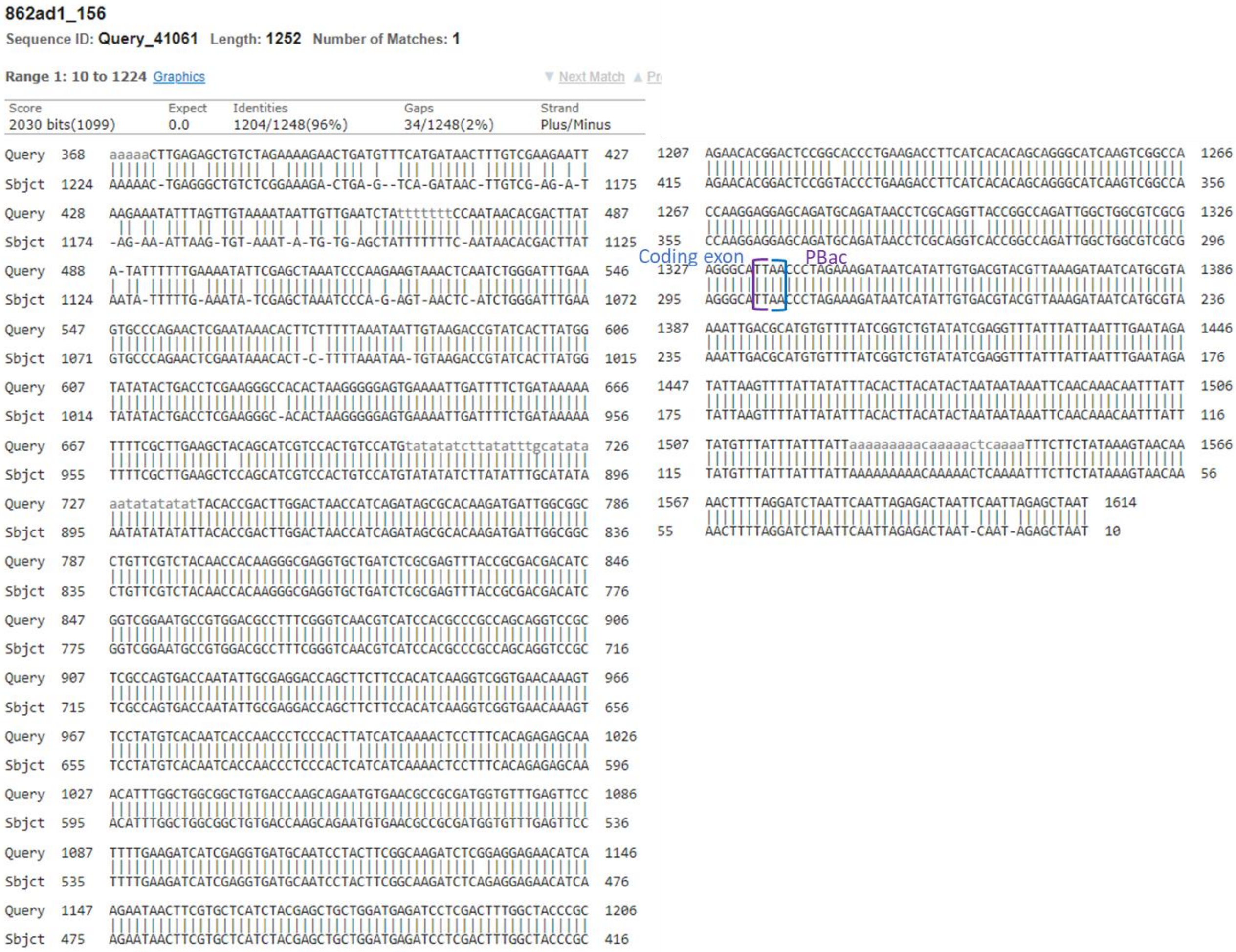

☎ +8862-2651-1809 ✉
13F.-12, No. 93, Sec. 1, Xintai 5th Rd., Xizhi Dist., New Taipei City 221416, Taiwan (R.O.C)
wellgenetics.com

220862ad1_OWG2409 (Sbjct) vs 220862 donor design (Query)

**862ad1_2409**

Sequence ID: **Query_41062** Length: **1270** Number of Matches: **1**

**Range 1: 9 to 1261** Graphics ▼ Next Match ▲ Pr

| Score | Expect | Identities | Gaps | Strand |
|---|---|---|---|---|
| 2026 bits(1097) | 0.0 | 1229/1286(96%) | 36/1286(2%) | Plus/Plus |

```
Query  2879  ACTTACCGCATTGACAAGCACGCCTCACGGGAGCTCCAAGCGGCGACTGAGATGTCCTAA  2938
             ||||||||||||||| ||||||||||||||||||||||||||||||||||||||||||||
Sbjct  9     ACTTACCGCATTGAC-AGCACGCCTCACGGGAGCTCCAAGCGGCGACTGAGATGTCCTAA  67

Query  2939  ATGCACAGCGACGGATTCGCGCTATTTAGAAAGAGAGAGCAATATTTCAAGAATGCATGC  2998
             ||||||||||||||||||||||||||||||||||||||||||||||||||||||||||||
Sbjct  68    ATGCACAGCGACGGATTCGCGCTATTTAGAAAGAGAGAGCAATATTTCAAGAATGCATGC  127
                                        PBac               R168W
Query  2999  GTCAATTTTACGCAGACTATCTTTCTAGGGTTAAGTACCGGTGGAACGAGCTTTTTCTGG  3058
             ||||||||||||||||||||||||||||||||||||||||||||||||||||||||||||
Sbjct  128   GTCAATTTTACGCAGACTATCTTTCTAGGGTTAAGTACCGGTGGAACGAGCTTTTTCTGG  187
                                                               Coding exon
Query  3059  ACGTATTGGAGTACGTGAATCTGCTGATGAGCCCGCAGGGTCAGGTTTTGTCTGCCCACG  3118
             ||||||||||||||||||||||||||||||||||||||||||||||||||||||| ||||
Sbjct  188   ACGTATTGGAGTACGTGAATCTGCTGATGAGCCCGCAGGGTCAGGTTTTGTCTGCTCACG  247

Query  3119  TGGCCGGCAAGGTGGTAATGAAGTCGTATTTGTCGGGTAAGTTAGCATAAATAATCTAGA  3178
             ||||||||||||||||||||||||||||||||||||||||||||||||||||||||||||
Sbjct  248   TGGCCGGCAAGGTGGTAATGAAGTCGTATTTGTCGGGTAAGTTAGCATAAATAATCTAGA  307

Query  3179  CATTATTCCTTTTAATAATCGCCATGTTGTAGGCATGCCCGAGTGCAAGTTCGGGATTAA  3238
             |||||||||||||||||||||||||||||||||||||||||||||||||||||| |||||
Sbjct  308   CATTATTCCTTTTAATAATCGCCATGTTGTAGGCATGCCCGAGTGCAAGTTCGGCATTAA  367

Query  3239  CGACAAGATCGTGATGGAGTCTAAGGGACGCGGTCTCTCCGGAAATTCAGAGGCGGAAAC  3298
             ||||||||||||||||||||||||||||||||||||||||||||||||||||||||||||
Sbjct  368   CGACAAGATCGTGATGGAGTCTAAGGGACGCGGTCTCTCCGGAAATTCAGAGGCGGAAAC  427

Query  3299  CTCACGCTCCGGCAAGCCCGTCGTGGTCATCGATGACTGCCAGTTCCATCAGTGCGTCAA  3358
             ||||||||||||||||||||||||||||||||||||||||||||||||||||||||||||
Sbjct  428   CTCACGCTCCGGCAAGCCCGTCGTGGTCATCGATGACTGCCAGTTCCATCAGTGCGTCAA  487

Query  3359  GCTAAGCAAATTCGAGACGGAGCATTCGATCAGCTTCATCCCGCCGGACGGGGAGTTCGA  3418
             |||||||||||||||||||||||||||||||||||| |||||||||||||||||||||||
Sbjct  488   GCTAAGCAAATTCGAGACGGAGCATTCGATCAGCTTTATCCCGCCGGACGGGGAGTTCGA  547

Query  3419  GCTGATGCGTTACCGTACCACCAAAGACATTTCGCTGCCATTCCGAGTCATCCCGCTGGT  3478
             ||||||||||||||||||||||||||||||||||||||||||||||||||||||||||||
Sbjct  548   GCTGATGCGTTACCGTACCACCAAAGACATTTCGCTGCCATTCCGAGTCATCCCGCTGGT  607

Query  3479  GCGGGAGGTGGGCCGCACCAAGATGGAGGTTAAGGTTGTGCTGAAGTCCAACTTTAAGCC  3538
             ||||||||||||||||||||||||||||||||||||||||||||||||||||||||||||
Sbjct  608   GCGGGAGGTGGGCCGCACCAAGATGGAGGTTAAGGTTGTGCTGAAGTCCAACTTTAAGCC  667

Query  3539  CTCACTGCTGGGCCAAAAGATCGAGGTGAAGATACCAACCCCGCTCAATACATCGGGCGT  3598
             ||||||||||||||||||||||||||||||||||||||||||||||||||||||||||||
Sbjct  668   CTCACTGCTGGGCCAAAAGATCGAGGTGAAGATACCAACCCCGCTCAATACATCGGGCGT  727

Query  3599  GCAGCTCATCTGCCTAAAGGGCAAAGCCAAATATAAGGCTTCGGAGAACGCGATCGTGTG  3658
             |||||||||||||||||||||||||||||||| |||||||||||||||||||||||||||
Sbjct  728   GCAGCTCATCTGCCTAAAGGGCAAAGCCAAATACAAGGCTTCGGAGAACGCGATCGTGTG  787
```

☎ +8862-2651-1809 ✉
13F.-12, No. 93, Sec. 1, Xintai 5th Rd., Xizhi Dist., New Taipei City 221416, Taiwan (R.O.C)
wellgenetics.com

**Cloning Method:**

The two homology arms of *Ap-2mu/CG7057* were amplified by Phusion High-Fidelity DNA Polymerase (Thermo Scientific) from genomic DNA at optimized condition, which reflects sequences of the injection strain *in vivo*. Any mismatches found in coding regions are considered as polymorphisms. The cassette PBacDsRed was obtained from sequence verified plasmid stock by restriction enzyme digestion. The cassette and two homology arms with point mutation were cloned by sequence-and ligation-independent method into vector pUC57-Kan, followed by standard transformation protocol, colony PCR selection and sequencing. Donor plasmid sequences shown in this report were fetched from FlyBase. Two homology arms and junctions of cassette fragment(s) were confirmed by PCR and sequencing. Only SNPs affecting amino acid sequences will be highlighted in plasmid sequence and will be further confirmed by independent genomic sequencing.

**Blat and Blast Method:**

Sequencing results were Blat[1] against *Drosophila melanogaster* genome (Aug 2014 Assembly, BDGP Release 6) using UCSC Genome Bioinformatics[2]. Sequence alignment shown here were using BLAST[3] (The Basic Local Alignment Search Tool) finds regions of local similarity between sequences.

[1] Kent WJ. BLAT - the BLAST-like alignment tool. Genome Res. 2002 Apr;12(4):656-64.

[2] Kent WJ, Sugnet CW, Furey TS, Roskin KM, Pringle TH, Zahler AM, Haussler D. The human genome browser at UCSC. Genome Res. 2002 Jun;12(6):996-1006.

[3] BLAST is a registered trademark of the National Library of Medicine.

+8862-2651-1809
13F.-12, No. 93, Sec. 1, Xintai 5th Rd., Xizhi Dist., New Taipei City 221416, Taiwan (R.O.C)
wellgenetics.com

WELLGENETICS

### CRISPR Validation Report

**Report No.:** RWGa4451 **Date:** 2023.07.21

****************************************************************************

**Gene:** *Ap-2mu/CG7057* **Case No.:** 220862

**Project:** introducing a point mutation R168W of AP-2mu using PBac system to facilitate genetic screening

**Method:** CRISPR/Cas9-mediated genome editing by homology-dependent repair (HDR) using 1 guide RNA(s) and a dsDNA plasmid donor

**Alleles:**

*220862A* *w[*];; AP-2mu R168W CRISPR{PBacDsRed} / TM6B, Tb[1]*

*SWGa7526* Insertion locus is validated by genomic PCR and sequencing. Homozygous lethal.

*220862B* *w[*];; AP-2mu R168W CRISPR{PBacDsRed} / TM6B, Tb[1]*

*SWGa7527* Insertion locus is validated by genomic PCR. Homozygous lethal.

*220862C* *w[*];; AP-2mu R168W CRISPR{PBacDsRed} / TM6B, Tb[1]*

*SWGa7528* Insertion locus is validated by genomic PCR. Homozygous lethal.

☎ +8862-2651-1809 ✉
13F.-12, No. 93, Sec. 1, Xintai 5th Rd., Xizhi Dist., New Taipei City 221416, Taiwan (R.O.C)
wellgenetics.com

Methods

**Genome Editing Map:**

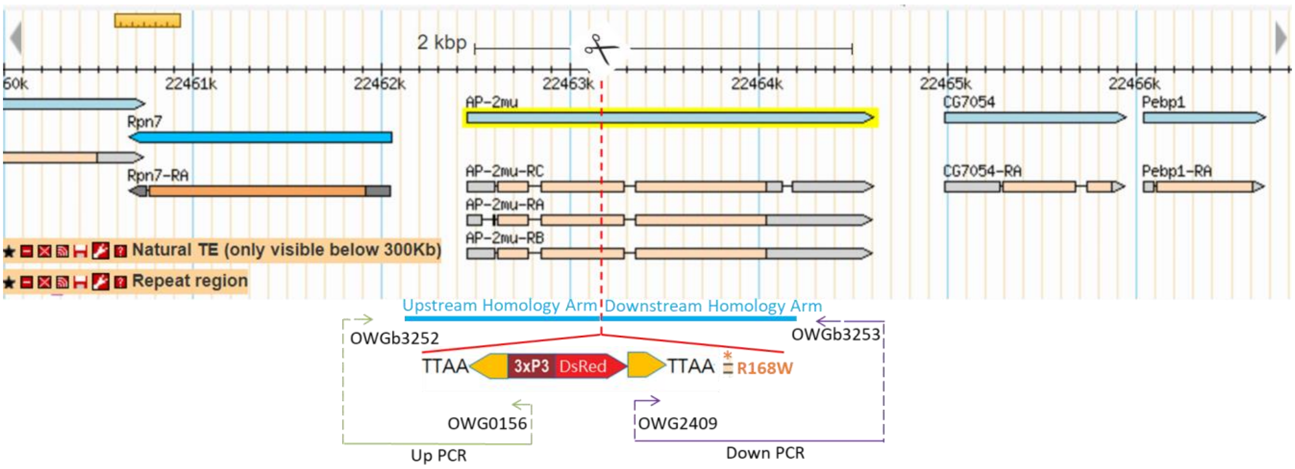

**Strategy:**

Using genomic PCR and sequencing methods to verify CRISPR alleles of *Ap-2mu/CG7057* by testing if the knock-in cassette PBacDsRed is correctly used as template for homology-directed repair and has been incorporated into guide RNA cutting site.

**Upstream PCR:**

Forward primer (OWGb3252) is designed at 5' outside of upstream homology arm and reverse primer (OWG0156) is designed at 3xP3 promoter. Only cassette PBacDsRed is inserted into the gene locus will have the amplicon.

**Downstream PCR:**

Forward primer (OWG2409) is designed at 5' PBac terminal repeats and reverse primer (OWGb3253) is designed at 3' outside of downstream homology arm as illustrated above. Only cassette PBacDsRed is inserted into the gene locus will have the amplicon.

**Primers:**

OWGb3252 5'- TTGTATTCCGCCAGGGTAAG
OWG0156 5'- CGAGGGTTCGAAATCGATAA
OWG2409 5'- TTTGACTCACGCGGTCGTTA
OWGb3253 5'- ACGTGAGCGTGAGATTTGTG

+8862-2651-1809
13F.-12, No. 93, Sec. 1, Xintai 5th Rd., Xizhi Dist., New Taipei City 221416, Taiwan (R.O.C)
wellgenetics.com

#### Genomic PCR

**Gel:**

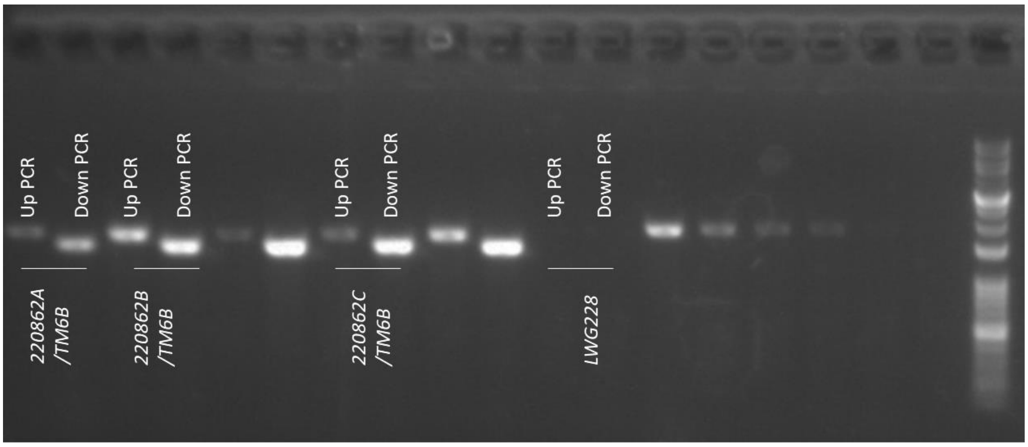

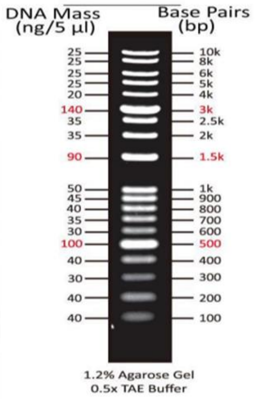

**Conclusion:**

PCR band at expected size was observed from heterozygous samples of *220862A, 220862B* and *220862C* for both upstream PCR (1652bp) and downstream PCR (1420bp), suggesting that cassette PBacDsRed was inserted into *Ap-2mu/CG7057* gene locus at correct orientation. No band at expected sizes was observed from negative control sample *LWG228*, indicating that high specificity of the PCR reactions. PCR products of *220862A* were sent for sequencing confirmation.

**Methods:**

Genomic DNA was obtained from single fly of each stock following single-fly DNA prep. Injection strain *[LWG228] w[1118]* was used as a negative control. PCR was performed using KOD-FX (TOYOBO) on BioRad S1000 Thermal Cycler. 1kb plus DNA Ladder from Biomate was used as reference.

+8862-2651-1809
13F.-12, No. 93, Sec. 1, Xintai 5th Rd., Xizhi Dist., New Taipei City 221416, Taiwan (R.O.C)
wellgenetics.com

**Genomic Sequencing**

**Sequences:**

>220862A_upPCR_OWGb3252

TTGCCATGACTGGACAGCTCAGGTTCGGGTTCTTCTCCAGACCCTGCTCCTCCAAGTTTTCGGCAGGCATTTT
GGTCGGATTTGTTGGCTTCGGTTTTTTAAACACAGAAAAATAGTTGATTTTCTTGACAAAACTCAATTTTGTTT
TTGTCAAATCGCAGAGTCATGTACTACCAGTGTGACCCCAAATTATCGATAAATTATACCGCATATTTTTACATTG
CCAAAAATACCAGAGCGATGTCCATCAAGATAGCGACGAAATTAGAACAGTGCAATTGCCAATTGGGAATTTG
TATTTTAATTTATTTTTAAATTCTGAAAGTAATTTTAATTTAAAAAAAACTTGAGAGCTGTCTAGAAAAGAACTG
ATGTTTCATGATAACTTTGTCGAAGAATTAAGAAATATTTAGTTGTAAAATAATTGTTGAAGCTATTTTTTTCCAA
TAACACGACTTATATATTTTTTGAAAATATTCGAGCTAAATCCCAAGAAGTAAACTCAATCTGGGATTTGAAGTG
CCCAGAACTCGAATAAACACTTCTTTTTAAATAATTGTAAGACCGTATCACTTATGGTATATACTGACCTCGAAG
GGCCACACTAAGGGGGAGTGAAAATTGATTTTCTGATAAAAATTTTCGCTTGAAGCTCCAGCATCGTCCACTG
TCCATGTATATATCTTATATTTGCATATAAATATATATATTACACCGACTTGGACTAACCATCAGATAGCGCACAAGA
TGATTGGCGGCCTGTTCGTCTACAACCACAAGGGCGAGGTGCTGATCTCGCGAGTTTACCGCGACGACATCG
GTCGGAATGCCGTGGACGCCTTTCGGGTCAACGTCATCCACGCCCGCCAGCAGGTCCGCTCGCCAGTGACCA
ATATTGCGAGGACCAGCTTCTTCCACATCAAGGTCGGTGAACAAAGTTCCTATGTCACAATCACCAACCCTCCC
ACTCATCATCAAAACTCCTTTCACAGAGAGCAAACATTTGGCTGGCGGCTGTGACCAAGCAGAATGTGAACG
CGCGATGGTGTTTGAGTTCCTTTTGAGATCATCGAGTGATGCAATCTACTTCGGCAGATCTCAGAGGAGACAT
CAGATACTCGTGCTCATCTACGAGCTGCTGATGGAATCTCGACTGCTACCGCAGACACGACTCGTACCTGAGA
CTCATCACACAGCAGCATCAGTCGCCACAAGAGACG

>220862A_upPCR_OWG0156

ACGTCAACTGATTAGCTCTATTGATTAGTCTCTAATTGAATTAGATCCTAAAAGTTTTGTTACTTTATAGAAGAAA
TTTTGAGTTTTTGTTTTTTTTTAATAAATAAATAAACATAAATAAATTGTTTGTTGAATTTATTATTAGTATGTAAG
TGTAAATATAATAAAACTTAATATCTATTCAAATTAATAAATAAACCTCGATATACAGACCGATAAAACACATGCGT
CAATTTTACGCATGATTATCTTTAACGTACGTCACAATATGATTATCTTTCTAGGGTTAATGCCCTCGCGACGCCA
GCCAATCTGGCCGGTGACCTGCGAGGTTATCTGCATCTGCTCCTCCTTGGTGGCCGACTTGATGCCCTGCTGT
GTGATGAAGGTCTTCAGGGTACCGGAGTCCGTGTTCTGCGGGTAGCCAAAGTCGAGGATCTCATCCAGCAGC
TCGTAGATGAGCACGAAGTTATTCTTGATGTTCTCCTCTGAGATCTTGCCGAAGTAGGATTGCATCACCTCGAT
GATCTTCAAAAGGAACTCAAACACCATCGCGGCGTTCACATTCTGCTTGGTCACAGCCGCCAGCCAAATGTTT
GCTCTCTGTGAAAGGAGTTTTGATGATGAGTGGGAGGGTTGGTGATTGTGACATAGGAACTTTGTTCACCGA
CCTTGATGTGGAAGAAGCTGGTCCTCGCAATATTGGTCACTGGCGAGCGGACCTGCTGGCGGGCGTGGATGA
CGTTGACCCGAAAGGCGTCCACGGCATTCCGACCGATGTCGTCGCGGTAAACTCGCGAGATCAGCACCTCGC
CCTTGTGGTTGTAGACGAACAGGCCGCCAATCATCTTGTGCGCTATCTGATGGTTAGTCCAAGTCGGTGTAATA
TATATATTTATATGCAAATATAAGATATATACATGGACAGTGGACGATGCTGGAGCTTCAAGCGAAAATTTTTATC

+8862-2651-1809
13F.-12, No. 93, Sec. 1, Xintai 5th Rd., Xizhi Dist., New Taipei City 221416, Taiwan (R.O.C)
wellgenetics.com

AGAAAATCAATTTTCACTCCCCCTTAGTGTGCCCTTCGAGGTCAGTATATACCATAAGTGATACGGTCTTACAAT
TATTTAAAAAGAAGTGTTTTATTCGAGTTCTTGGGCCACTTCAAATCCCAGATTGAGTTTACTTCTTGGGATTTA
GCTCGAATATTTTTCAAAAAATTATATAAGTCGTGTTTATTGGAAAAAATAGCTCACATTATTTACACTAAATATT
TTCTTAGTTCTTTCGACAAGTTATTCATGAAACATCCAGTTCTTTTTCCTAGAACCGCCTCTCTCAG

>220862A_downPCR_OWG2409
GGGATGGTGACTTACCGCATTGACAGCACGCCTCACGGGAGCTCCAAGCGGCGACTGAGATGTCCTAAATGC
ACAGCGACGGATTCGCGCTATTTAGAAAGAGAGAGCAATATTTCAAGAATGCATGCGTCAATTTTACGCAGAC
TATCTTTCTAGGGTTAAGTACCGGTGGAACGAGCTTTTTCTGGACGTATTGGAGTACGTGAATCTGCTGATGAG
CCCGCAGGGTCAGGTTTTGTCTGCTCACGTGGCCGGCAAGGTGGTAATGAAGTCGTATTTGTCGGGTAAGTT
AGCATAAATAATCTAGACATTATTCCTTTTAATAATCGCCATGTTGTAGGCATGCCCGAGTGCAAGTTCGGCATT
AACGACAAGATCGTGATGGAGTCTAAGGGACGCGGTCTCTCCGGAAATTCAGAGGCGGAAACCTCACGCTCC
GGCAAGCCCGTCGTGGTCATCGATGACTGCCAGTTCCATCAGTGCGTCAAGCTAAGCAAATTCGAGACGGAG
CATTCGATCAGCTTTATCCCGCCGGACGGGGAGTTCGAGCTGATGCGTTACCGTACCACCAAAGACATTTCGC
TGCCATTCCGAGTCATCCCGCTGGTGCGGGAGGTGGGCCGCACCAAGATGGAGGTTAAGGTTGTGCTGAAG
TCCAACTTTAAGCCCTCACTGCTGGGCCAAAAGATCGAGGTGAAGATACCAACCCCGCTCAATACATCGGGCG
TGCAGCTCATCTGCCTAAAGGGCAAAGCCAAATACAAGGCTTCGGAGAACGCGATCGTGTGGAAGATTAAGC
GCATGGCGGGCATGAAGGAGACACAGCTGTCCGCGGAAATCGAGCTTTTGGAGACGGACACCAAGAAGAA
GTGGACTCGGCCGCCCATCTCCATGAACTTTGAGGTGCCATTCGCGCCGTCCGGCTTCAAGGTACGCTACCTG
AAAGTGTTCGAGCCCAAGCTCAACTACTCCGACCACGATGTGGTCAAATGGGTGCGCTACATCGGACGCAGT
GGCTTGTATGAGACGCGCTGCTAGGGCGCCCAAGCATCCCCATCACGGAGCAGATTATCACATTGATTTAACG
AGATGATGGAGAAAATGTACGCATCTATTGGTATTTCCTCATCAAATGACAGCAGCTTATGTATTCAGATAGTCA
GCAGTCGAGACGCCCGGAGTAGCGATCCACCCCCGGAAACCCTATTGCCGTATATACGGTA

>220862A_downPCR_OWGb3253
GTGGGTACGAGCAATAAAACAACATAGTTTACATTTACATAGTAGGGTAAATCATAGTACCTAGTAGTGGTATGT
CTATGGGGAACACGCCCCGCCCGCATAGCCAGTGCTCGCGGAGGCAGTGTGTGCGGTATACGGCATAGGGTG
TCCGGGGTGGATCCGCTACTCGGGCGTCCTCCGGACTTGCTGGAACTAATCTTGGAATACATAAGCTGCTGGT
TCAATTTGATTGAGGAAATACCAATAGATGCGGTAACATTTTCTCCATTCATCTCGTTAAATTCAATTGTGATAAT
CTTGCTCCGTGATGGGGATGCTTGGGCGCCCTAGCAGCGCGTCTCATACAAGCCACTGCGTCCGATGTAGCGC
ACCCATTTGACCACATCGTGGTCGGAGTAGTTGAGCTTGGGCTCGAACACCTTCAGGTAGCGTACCTTGAAGC
CGGACGGCGCGAATGGCACCTCAAAGTTCATGGAGATGGGCGGCCGAGTCCACTTCTTCTTGGTGTCCGTCT
CCAAAAGCTCGATTTCCGCGGACAGCTGTGTCTCCTTCATGCCCGCCATGCGCTTAATCTTCCACACGATCGCG
TTCTCCGAAGCCTTGTATTTGGCTTTGCCCTTTAGGCAGATGAGCTGCACGCCCGATGTATTGAGCGGGGTTG
GTATCTTCACCTCGATCTTTTGGCCCAGCAGTGAGGGCTTAAAGTTGGACTTCAGCACAACCTTAACCTCCATC

TTGGTGCGGCCCACCTCCCGCACCAGCGGGATGACTCGGAATGGCAGCGAAATGTCTTTGGTGGTACGGTAA
CGCATCAGCTCGAACTCCCCGTCCGGCGGGATAAAGCTGATCGAATGCTCCGTCTCGAATTTGCTTAGCTTGA
CGCACTGATGGAACTGGCAGTCATCGATGACCACGACGGGCTTGCCGGAGCGTGAGGTTTCCGCCTCTGAAT
TTCCGGAGAGACCGCGTCCCTTAGACTCCATCACGATCTTGTCGTTAATGCCGAACTTGCACTCGGGCATGCCT
ACACATGGCGATTATTAAAAGGATATGTCTAGATTATTTATGCTAACTTACCCGACAATACGACTTCATTACCACC
TTTGCCGGCCACGTGAGCAGACAAACCTGACCCTGCGGGCCTCATCAGCAGAATCCACGTACTCCATACGTCC
AGAAAAAAGCCTCGTTCCACCCGGGTACTTACCCTAGAAAAGATAGTCTCTGCGCGTTA

**Blat Results:**

Upstream and downstream homology arms of donor plasmid are as indicated. Sequences of *220862A* were read from flanking region to homology arms, suggesting these sequences were specific to gene *Ap-2mu/CG7057* and cassette PBacDsRed was inserted into designed guide RNA cutting site at correct orientation in the genome of *220862A* (see overview below). Eight mismatches (boxed in blue) within the homology arm region were found in the CDS of *Ap-2mu/CG7057.* All mismatches have been reported in cloning report (RWGa3634). Orange box indicates the designed point mutation of *Ap-2mu/CG7057*.

Overview:

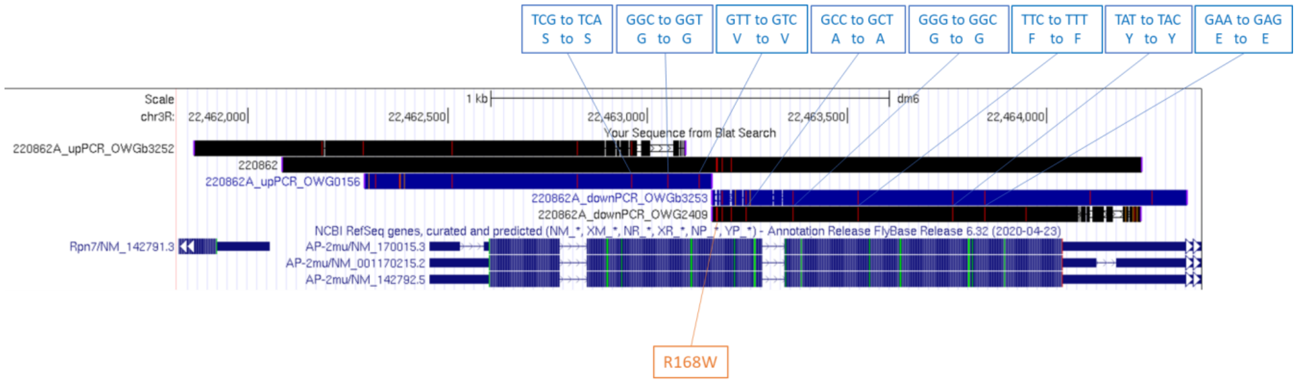

☎ +8862-2651-1809 ✉
13F.-12, No. 93, Sec. 1, Xintai 5th Rd., Xizhi Dist., New Taipei City 221416, Taiwan (R.O.C)
wellgenetics.com

Sequence reads (>220862A_upPCR_OWG0156 and >220862A_downPCR_OWG2409) were aligned to 220862 donor design using BLAST to check the junction of cassette PBacDsRed insertion. The sequences of PiggyBac (bracketed in purple) were present in *220862A*. The cassette PBacDsRed was correctly inserted in coding exon (bracketed in blue) of *Ap-2mu/CG7057* in *220862A.* Design point mutations were shaded in orange.

220862A_upPCR_OWG0156 (Sbjct) vs 220862 donor design (Query)

**862bAbb3252_156**

Sequence ID: **Query_54519** Length: **1257** Number of Matches: **1**

**Range 1: 11 to 1236** Graphics ▼ Next Match ▲ Pre

| Score | Expect | Identities | Gaps | Strand |
|---|---|---|---|---|
| 2115 bits(1145) | 0.0 | 1209/1236(98%) | 20/1236(1%) | Plus/Minus |

```
Query  548   AAAAGAACT-GATGTTTCATG-ATAACTTTGTCG-AAGAATTAAG-AAATATTTAGTTGT  603
Sbjct  1236  AAAAGAACTGGATGTTTCATGAATAAC-TTGTCGAAAGAACTAAGAAAATATTTAG-TGT  1179

Query  604   AAAATAATTGTTGAATCTAttttttCCAAT-AACACGACTTATATA-TTTTTTG-AAAA  660
Sbjct  1178  -AAATAA-TG-TG-AGCTA-TTTTTTCCAATAAACACGACTTATATAATTTTTTGAAAAA  1124

Query  661   TATTCGAGCTAAATCCCAAGAAGTAAACTCAATCTGGGATTTGAAGTG-CCC-AGAACTC  718
Sbjct  1123  TATTCGAGCTAAATCCCAAGAAGTAAACTCAATCTGGGATTTGAAGTGGCCCAAGAACTC  1064

Query  719   GAAT-AAACACTTCTTTTTAAATAATTGTAAGACCGTATCACTTATGGTATATACTGACC  777
Sbjct  1063  GAATAAAACACTTCTTTTTAAATAATTGTAAGACCGTATCACTTATGGTATATACTGACC  1004

Query  778   TCGAAGGGCCACACTAAGGGGGAGTGAAAATTGATTTTCTGATAAAAATTTTCGCTTGAA  837
Sbjct  1003  TCGAAGGGC-ACACTAAGGGGGAGTGAAAATTGATTTTCTGATAAAAATTTTCGCTTGAA  945

Query  838   GCTACAGCATCGTCCACTGTCCATGtatatatcttatatttgcatataaatatatatatT  897
Sbjct  944   GCTCCAGCATCGTCCACTGTCCATGTATATATCTTATATTTGCATATAAATATATATATT  885

Query  898   ACACCGACTTGGACTAACCATCAGATAGCGCACAAGATGATTGGCGGCCTGTTCGTCTAC  957
Sbjct  884   ACACCGACTTGGACTAACCATCAGATAGCGCACAAGATGATTGGCGGCCTGTTCGTCTAC  825

Query  958   AACCACAAGGGCGAGGTGCTGATCTCGCGAGTTTACCGCGACGACATCGGTCGGAATGCC  1017
Sbjct  824   AACCACAAGGGCGAGGTGCTGATCTCGCGAGTTTACCGCGACGACATCGGTCGGAATGCC  765

Query  1018  GTGGACGCCTTTCGGGTCAACGTCATCCACGCCCGCCAGCAGGTCCGCTCGCCAGTGACC  1077
Sbjct  764   GTGGACGCCTTTCGGGTCAACGTCATCCACGCCCGCCAGCAGGTCCGCTCGCCAGTGACC  705

Query  1078  AATATTGCGAGGACCAGCTTCTTCCACATCAAGGTCGGTGAACAAAGTTCCTATGTCACA  1137
Sbjct  704   AATATTGCGAGGACCAGCTTCTTCCACATCAAGGTCGGTGAACAAAGTTCCTATGTCACA  645

Query  1138  ATCACCAACCCTCCCACTTATCATCAAAACTCCTTTCACAGAGAGCAAACATTTGGCTGG  1197
Sbjct  644   ATCACCAACCCTCCCACTCATCATCAAAACTCCTTTCACAGAGAGCAAACATTTGGCTGG  585

Query  1198  CGGCTGTGACCAAGCAGAATGTGAACGCCGCGATGGTGTTTGAGTTCCTTTTGAAGATCA  1257
Sbjct  584   CGGCTGTGACCAAGCAGAATGTGAACGCCGCGATGGTGTTTGAGTTCCTTTTGAAGATCA  525

Query  1258  TCGAGGTGATGCAATCCTACTTCGGCAAGATCTCGGAGGAGAACATCAAGAATAACTTCG  1317
Sbjct  524   TCGAGGTGATGCAATCCTACTTCGGCAAGATCTCAGAGGAGAACATCAAGAATAACTTCG  465
```

```
1318  TGCTCATCTACGAGCTGCTGGATGAGATCCTCGACTTTGGCTACCCGCAGAACACGGACT  1377
464   TGCTCATCTACGAGCTGCTGGATGAGATCCTCGACTTTGGCTACCCGCAGAACACGGACT  405

1378  CCGGCACCCTGAAGACCTTCATCACACAGCAGGGCATCAAGTCGGCCACCAAGGAGGAGC  1437
404   CCGGTACCCTGAAGACCTTCATCACACAGCAGGGCATCAAGTCGGCCACCAAGGAGGAGC  345
                                                    Coding exon   PBac
1438  AGATGCAGATAACCTCGCAGGTTACCGGCCAGATTGGCTGGCGTCGCGAGGGCATTAACC  1497
344   AGATGCAGATAACCTCGCAGGTCACCGGCCAGATTGGCTGGCGTCGCGAGGGCATTAACC  285

1498  CTAGAAAGATAATCATATTGTGACGTACGTTAAAGATAATCATGCGTAAAATTGACGCAT  1557
284   CTAGAAAGATAATCATATTGTGACGTACGTTAAAGATAATCATGCGTAAAATTGACGCAT  225

1558  GTGTTTTATCGGTCTGTATATCGAGGTTTATTTATTAATTTGAATAGATATTAAGTTTTA  1617
224   GTGTTTTATCGGTCTGTATATCGAGGTTTATTTATTAATTTGAATAGATATTAAGTTTTA  165

1618  TTATATTTACACTTACATACTAATAATAAATTCAACAAACAATTTATTTATGTTTATTTA  1677
164   TTATATTTACACTTACATACTAATAATAAATTCAACAAACAATTTATTTATGTTTATTTA  105

1678  TTTATTaaaaaaaaacaaaaactcaaaaTTTCTTCTATAAAGTAACAAAACTTTTAGGAT  1737
104   TTTATTAAAAAAAAACAAAAACTCAAAATTTCTTCTATAAAGTAACAAAACTTTTAGGAT  45

1738  CTAATTCAATTAGAGACTAATTCAATTAGAGCTAAT  1773
44    CTAATTCAATTAGAGACTAAT-CAAT-AGAGCTAAT  11
```

☎ +8862-2651-1809 ✉
13F.-12, No. 93, Sec. 1, Xintai 5th Rd., Xizhi Dist., New Taipei City 221416, Taiwan (R.O.C)
wellgenetics.com

220862A_downPCR_OWG2409 (Sbjct) vs 220862 donor design (Query)

**862bAbb3253_2409**

**Sequence ID: Query_54521 Length: 1223 Number of Matches: 1**

**Range 1: 10 to 1216** Graphics ▼ Next Match ▲ Pr

| Score | Expect | Identities | Gaps | Strand |
|---|---|---|---|---|
| 2069 bits(1120) | 0.0 | 1196/1228(97%) | 24/1228(1%) | Plus/Plus |

```
Query  3038  ACTTACCGCATTGACAAGCACGCCTCACGGGAGCTCCAAGCGGCGACTGAGATGTCCTAA  3097
             ||||||||||||||| ||||||||||||||||||||||||||||||||||||||||||||
Sbjct  10    ACTTACCGCATTGAC-AGCACGCCTCACGGGAGCTCCAAGCGGCGACTGAGATGTCCTAA  68

Query  3098  ATGCACAGCGACGGATTCGCGCTATTTAGAAAGAGAGAGCAATATTTCAAGAATGCATGC  3157
             ||||||||||||||||||||||||||||||||||||||||||||||||||||||||||||
Sbjct  69    ATGCACAGCGACGGATTCGCGCTATTTAGAAAGAGAGAGCAATATTTCAAGAATGCATGC  128
                                          PBac      Coding exon
Query  3158  GTCAATTTTACGCAGACTATCTTTCTAGGGTTAAGTACCGGTGGAACGAGCTTTTTCTGG  3217
             ||||||||||||||||||||||||||||||||||||||||||||||||||||||||||||
Sbjct  129   GTCAATTTTACGCAGACTATCTTTCTAGGGTTAAGTACCGGTGGAACGAGCTTTTTCTGG  188
                                                      R168W
Query  3218  ACGTATTGGAGTACGTGAATCTGCTGATGAGCCCGCAGGGTCAGGTTTTGTCTGCCCACG  3277
             ||||||||||||||||||||||||||||||||||||||||||||||||||||||| ||||
Sbjct  189   ACGTATTGGAGTACGTGAATCTGCTGATGAGCCCGCAGGGTCAGGTTTTGTCTGCTCACG  248

Query  3278  TGGCCGGCAAGGTGGTAATGAAGTCGTATTTGTCGGGTAAGTTAGCATAAATAATCTAGA  3337
             ||||||||||||||||||||||||||||||||||||||||||||||||||||||||||||
Sbjct  249   TGGCCGGCAAGGTGGTAATGAAGTCGTATTTGTCGGGTAAGTTAGCATAAATAATCTAGA  308

Query  3338  CATTATTCCTTTTAATAATCGCCATGTTGTAGGCATGCCCGAGTGCAAGTTCGGGATTAA  3397
             |||||||||||||||||||||||||||||||||||||||||||||||||||||| |||||
Sbjct  309   CATTATTCCTTTTAATAATCGCCATGTTGTAGGCATGCCCGAGTGCAAGTTCGGCATTAA  368

Query  3398  CGACAAGATCGTGATGGAGTCTAAGGGACGCGGTCTCTCCGGAAATTCAGAGGCGGAAAC  3457
             ||||||||||||||||||||||||||||||||||||||||||||||||||||||||||||
Sbjct  369   CGACAAGATCGTGATGGAGTCTAAGGGACGCGGTCTCTCCGGAAATTCAGAGGCGGAAAC  428

Query  3458  CTCACGCTCCGGCAAGCCCGTCGTGGTCATCGATGACTGCCAGTTCCATCAGTGCGTCAA  3517
             ||||||||||||||||||||||||||||||||||||||||||||||||||||||||||||
Sbjct  429   CTCACGCTCCGGCAAGCCCGTCGTGGTCATCGATGACTGCCAGTTCCATCAGTGCGTCAA  488

Query  3518  GCTAAGCAAATTCGAGACGGAGCATTCGATCAGCTTCATCCCGCCGGACGGGGAGTTCGA  3577
             |||||||||||||||||||||||||||||||||||| |||||||||||||||||||||||
Sbjct  489   GCTAAGCAAATTCGAGACGGAGCATTCGATCAGCTTTATCCCGCCGGACGGGGAGTTCGA  548

Query  3578  GCTGATGCGTTACCGTACCACCAAAGACATTTCGCTGCCATTCCGAGTCATCCCGCTGGT  3637
             ||||||||||||||||||||||||||||||||||||||||||||||||||||||||||||
Sbjct  549   GCTGATGCGTTACCGTACCACCAAAGACATTTCGCTGCCATTCCGAGTCATCCCGCTGGT  608

Query  3638  GCGGGAGGTGGGCCGCACCAAGATGGAGGTTAAGGTTGTGCTGAAGTCCAACTTTAAGCC  3697
             ||||||||||||||||||||||||||||||||||||||||||||||||||||||||||||
Sbjct  609   GCGGGAGGTGGGCCGCACCAAGATGGAGGTTAAGGTTGTGCTGAAGTCCAACTTTAAGCC  668

Query  3698  CTCACTGCTGGGCCAAAAGATCGAGGTGAAGATACCAACCCCGCTCAATACATCGGGCGT  3757
             ||||||||||||||||||||||||||||||||||||||||||||||||||||||||||||
Sbjct  669   CTCACTGCTGGGCCAAAAGATCGAGGTGAAGATACCAACCCCGCTCAATACATCGGGCGT  728

Query  3758  GCAGCTCATCTGCCTAAAGGGCAAAGCCAAATATAAGGCTTCGGAGAACGCGATCGTGTG  3817
             ||||||||||||||||||||||||||||||||| ||||||||||||||||||||||||||
Sbjct  729   GCAGCTCATCTGCCTAAAGGGCAAAGCCAAATACAAGGCTTCGGAGAACGCGATCGTGTG  788

Query  3818  GAAGATTAAGCGCATGGCGGGCATGAAGGAGACACAGCTGTCCGCGGAAATCGAACTTTT  3877
             |||||||||||||||||||||||||||||||||||||||||||||||||||||| |||||
Sbjct  789   GAAGATTAAGCGCATGGCGGGCATGAAGGAGACACAGCTGTCCGCGGAAATCGAGCTTTT  848
```

☎ +8862-2651-1809 ✉
📍 13F.-12, No. 93, Sec. 1, Xintai 5th Rd., Xizhi Dist., New Taipei City 221416, Taiwan (R.O.C)
wellgenetics.com

**Methods:**

PCR bands of *220862A* were excised and submitted to Mission Biotech for gel extraction and sequencing. Sequencing results were Blat[1] against *Drosophila melanogaster* genome (Aug 2014 Assembly, BDGP Release 6) using UCSC Genome Bioinformatics[2]. Sequence alignment shown here were using BLAST[3] (The Basic Local Alignment Search Tool) finds regions of local similarity between sequences.

[1] Kent WJ. BLAT - the BLAST-like alignment tool. Genome Res. 2002 Apr;12(4):656-64.

[2]Kent WJ, Sugnet CW, Furey TS, Roskin KM, Pringle TH, Zahler AM, Haussler D. The human genome browser at UCSC. Genome Res. 2002 Jun;12(6):996-1006.

[3]BLASTis a registered trademark of the National Library of Medicine.

+8862-2651-1809
13F.-12, No. 93, Sec. 1, Xintai 5th Rd., Xizhi Dist., New Taipei City 221416, Taiwan (R.O.C)
wellgenetics.com

WellGenetics

### CRISPR Integration Report

| | | | |
|---|---|---|---|
| **Report No.:** | RWGa4452 | **Date:** | 2023.07.21 |
| **Reporter:** | Wei-Chi Ke | **E-mail:** | |

****************************************************************************

**Gene:** *Ap-2mu/CG7057* **Case No.:** 220862

**Project:** introducing a point mutation R168W of AP-2mu using PBac system to facilitate genetic screening

**Method:** To test donor plasmid integration in CRISPR editing alleles by genomic PCR

**Alleles:**

*220862A* *w[*];; AP-2mu R168W CRISPR{PBacDsRed} / TM6B, Tb[1]*
*SWGa7526* No donor plasmid integration.

*220862B* *w[*];; AP-2mu R168W CRISPR{PBacDsRed} / TM6B, Tb[1]*
*SWGa7527* No donor plasmid integration.

*220862C* *w[*];; AP-2mu R168W CRISPR{PBacDsRed} / TM6B, Tb[1]*
*SWGa7528* No donor plasmid integration.

+8862-2651-1809
13F.-12, No. 93, Sec. 1, Xintai 5th Rd., Xizhi Dist., New Taipei City 221416, Taiwan (R.O.C)
wellgenetics.com

**Method**

**Strategy:**

CRISPR/HDR (homology-dependent repair) may go through different repairing routes. Integration of donor plasmid in CRISPR editing alleles is a common phenomenon. Figure 1 shows three examples of HDR occurred in CRISPR editing. “No Integration” contains only one copy of selection marker. “Single Integration of Donor Plasmid” contains vector backbone, two copies of homology arms and selection marker. “Double Integration of Donor Plasmid” contains multiple copies of vector backbone, homology arms and selection marker. Integration Test is to exclude alleles with donor integration and only keep the lines with simple editing to avoid the genomic complications.

**Integration PCR 1 (F1 & R1):**

Forward primer (OWG5476) is designed at outside of upstream homology arm in backbone plasmid and reverse primer (OWG0156) is designed at3xP3 promoter. Single integration of donor plasmid and double integration of donor plasmids will have the amplicon. Correct editing will not have the amplicon. # indicates this set of primers could amplify large PCR product in double integration of donor plasmids.

**Integration PCR 2 (F2 & R2):**

Forward primer (OWG2409) is designed at 5’ PBac terminal repeat and reverse primer (OWG5477) is designed at outside of downstream homology arm in backbone plasmid. Single integration of donor plasmid and double integration of donor plasmids will have the amplicon. Correct editing will not have the amplicon. # indicates this set of primers could amplify large PCR product in double integration of donor plasmids.

**Primers:**

| | |
|---|---|
| OWG5476 | 5’-CAACTGTTGGGAAGGGCGAT |
| OWG0156 | 5’- CGAGGGTTCGAAATCGATAA |
| OWG2409 | 5’- TTTGACTCACGCGGTCGTTA |
| OWG5477 | 5’-CATTAGGCACCCCAGGCTTT |

☎ +8862-2651-1809 ✉
13F.-12, No. 93, Sec. 1, Xintai 5th Rd., Xizhi Dist., New Taipei City 221416, Taiwan (R.O.C)
wellgenetics.com

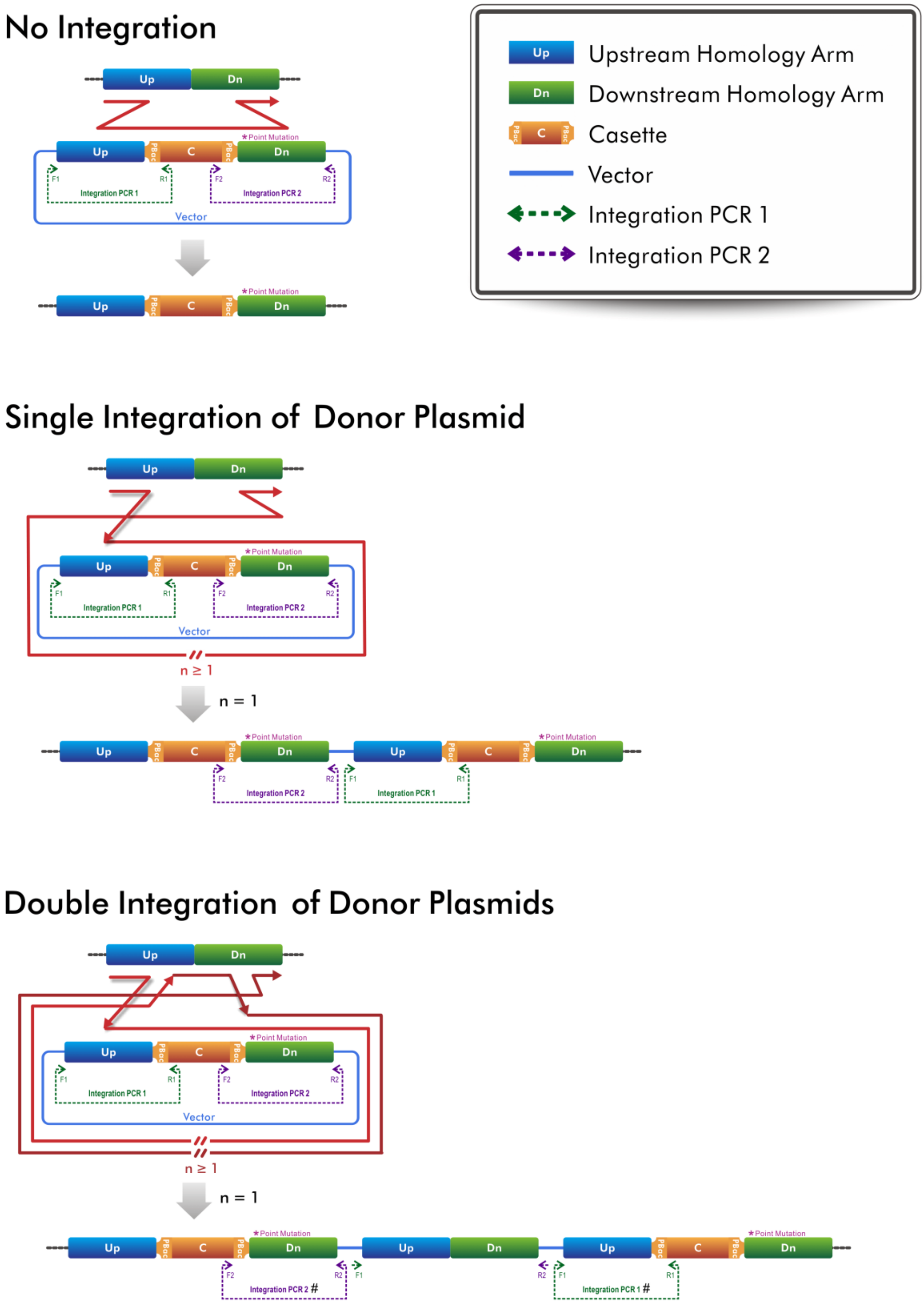

**Figure 1. Three examples of HDR occurred in CRISPR editing**

☎ +8862-2651-1809 ✉
13F.-12, No. 93, Sec. 1, Xintai 5th Rd., Xizhi Dist., New Taipei City 221416, Taiwan (R.O.C)
wellgenetics.com

#### Genomic PCR

**Gel:**

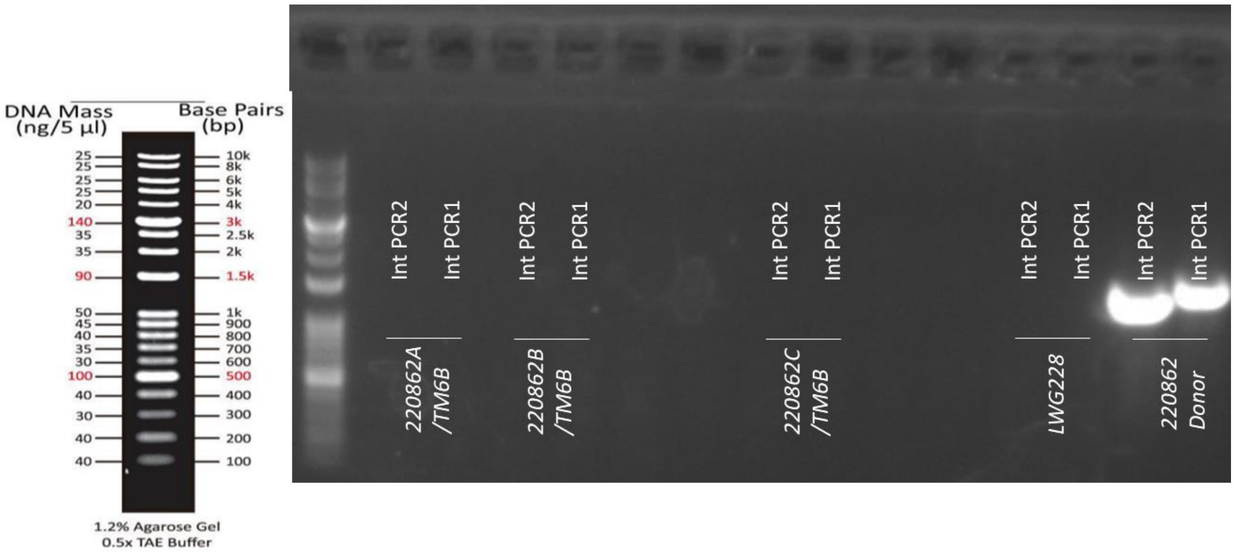

**Conclusion:**

No PCR bands at expected sizes were observed from heterozygous samples of *220862A, 220862B* and *220862C* for both Integration PCR 2 (1388bp) and Integration PCR 1 (1551bp), suggesting that no donor integration has occurred at *Ap-2mu/CG7057* gene locus. PCR bands at expected sizes were observed from positive control plasmid, *220862 donor* and no bands at expected sizes were observed from negative control sample *LWG228*, indicating that the primer pairs for integration test are highly specific.

**Methods:**

Genomic DNA was obtained from single fly of each stock following single-fly DNA prep. Injection strain *[LWG228] w[1118]* was used as a negative control and *220862 donor* was a positive control for integration examination. PCR was performed using KOD-FX (TOYOBO) on BioRad S1000 Thermal Cycler. 1kb plus DNA Ladder from Biomate was used as reference.

☎ +8862-2651-1809 ✉
13F.-12, No. 93, Sec. 1, Xintai 5th Rd., Xizhi Dist., New Taipei City 221416, Taiwan (R.O.C)
wellgenetics.com

### Excision Validation Report

| | | | |
|---|---|---|---|
| **Report No.:** | RWGa4913 | **Date:** | 2023.10.02 |
| **Reporter:** | Wei-Chi Ke | **E-mail:** | |

**************************************************************************

**Gene:** *Ap-2mu/CG7057* **Case No.:** 220862ex

**Project:** introducing a point mutation R168W of AP-2mu using PBac system to facilitate genetic screening

**Method:** Excision of selection marker by *PiggyBac* (PBac) Transposition

**Progenitor:** *[SWGa7526] 220862A/TM6B*
*w[*];; AP-2mu R168W CRISPR{PBacDsRed} / TM6B, Tb[1]*
Derived from Bloomington 8285
*w*; CyO, P{Tub-PBac\T}2 / wgSp-1 ; l(3)*[*] / TM6C, Sb, Tb, Hu, dfd-eYFP w+*

**Alleles:**

*220862ex2* *w[*];; AP-2mu R168W CRISPR / TM6B, Tb[1]*
*SWGa8276* Excision is validated by sequencing. Homozygous viable.

*220862ex3* *w[*];; AP-2mu R168W CRISPR / TM6B, Tb[1]*
*SWGa8277* Excision is **NOT** validated by genomic PCR. Homozygous viable.

*220862ex4* *w[*];; AP-2mu R168W CRISPR / TM6B, Tb[1]*
*SWGa8278* Excision is **NOT** validated by genomic PCR. Homozygous viable.

*220862ex6* *w[*];; AP-2mu R168W CRISPR / TM6B, Tb[1]*
*SWGa8279* Excision is **NOT** validated by genomic PCR. Homozygous viable.

**Methods**

#### Genome Editing Map:

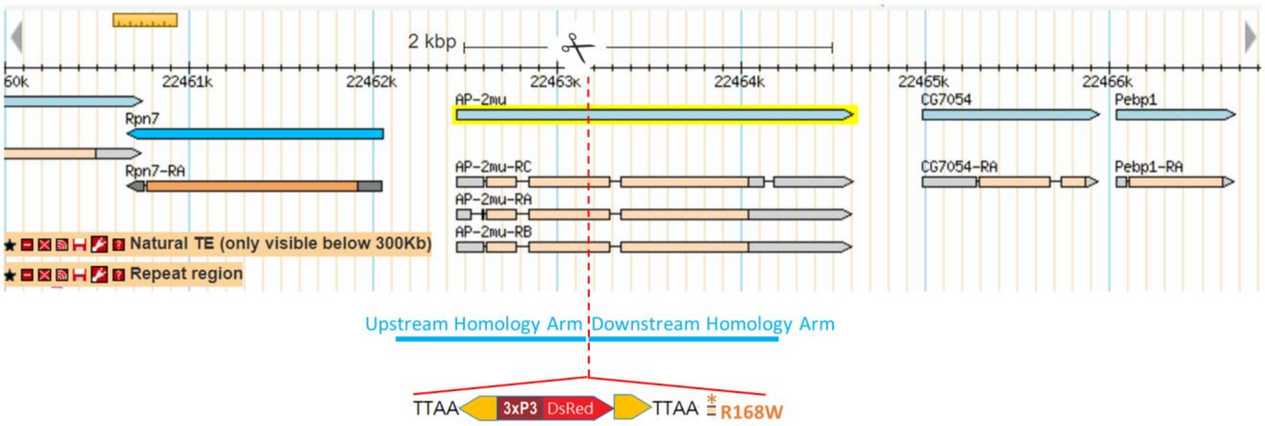

↓ **PBac Excision**

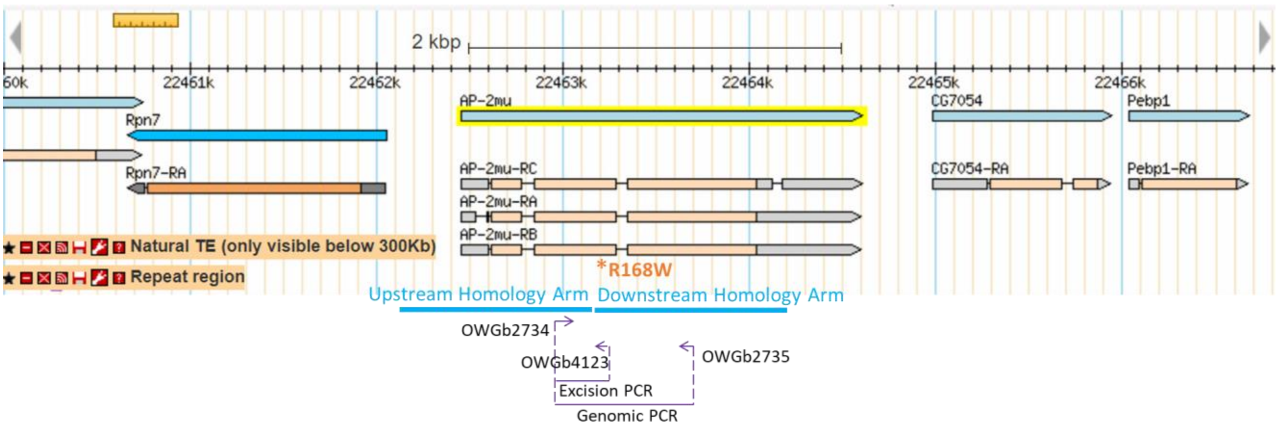

**Strategy:**

Using genomic PCR and sequencing methods to verify PiggyBac transposition alleles of *w[*];; AP-2mu R168W CRISPR{PBacDsRed} / TM6B, Tb[1]* fly by testing if the selection marker (3XP3-DsRed) is precisely excised and one TTAA sequence is left and embedded in coding exon of *Ap-2mu/CG7057*.

**Excision PCR:**

Forward primer (OWGb2734) is designed at upstream homology arm and reverse primer (OWGb4123) is specifically designed at R168W mutation as illustrated above. The R168W mutation

☎ +8862-2651-1809 ✉
13F.-12, No. 93, Sec. 1, Xintai 5th Rd., Xizhi Dist., New Taipei City 221416, Taiwan (R.O.C)
wellgenetics.com

is located at 3’ end of primer which leads no PCR product in injection strain control. The product length is 1956bp before excision and 260bp after excision.

**Genomic PCR:**

Forward primer (OWGb2734) is designed at upstream homology arm and reverse primer (OWGb2735) is designed at downstream homology arm as illustrated above. One of excision PCR-positive alleles will be performed genomic PCR. The genomic PCR band from excision allele will be sent for sequencing.

**Primers:**

OWGb2734 5’- GTGATGCAATCCTACTTCGGC
OWGb4123 5’- TCCAGAAAAAGCTCGTTCCA
OWGb2735 5’- TTAACCTCCATCTTGGTGCGG

☎ +8862-2651-1809 ✉
13F.-12, No. 93, Sec. 1, Xintai 5th Rd., Xizhi Dist., New Taipei City 221416, Taiwan (R.O.C)
wellgenetics.com

**Genomic PCR**

**Gel:**

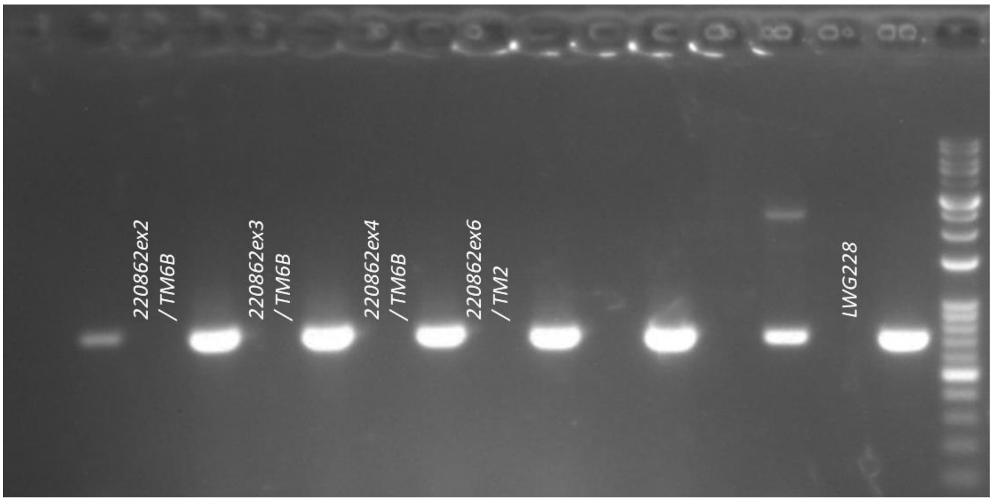

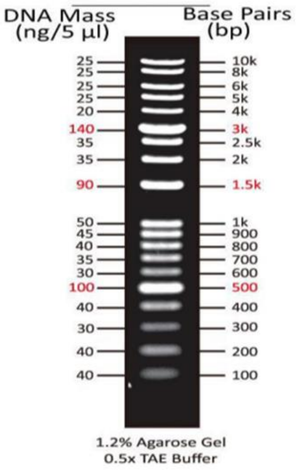

**Conclusion:**

No PCR bands at expected size were observed from heterozygous sample(s) of *220862ex2, 220862ex3, 220862ex4* and *220862ex6* lines for Excision PCR (260bp), suggesting that no specificity of the PCR reaction from primer pair, OWGb2734/OWGb4123. Since there was no conclusion from Excision PCR, one of alleles was randomly selected for sequencing confirmation. PCR product of *220862ex2* from primer pair, OWGb2734/OWGb2735, was chose and sent for sequencing.

**Methods:**

Genomic DNA was obtained from single fly of each stock following single-fly DNA prep. Injection strain *[LWG228] w[1118]* was used as a negative control. PCR was performed using KOD-FX (TOYOBO) on BioRad S1000 Thermal Cycler. 1kb plus DNA Ladder from Biomate was used as reference.

+8862-2651-1809
13F.-12, No. 93, Sec. 1, Xintai 5th Rd., Xizhi Dist., New Taipei City 221416, Taiwan (R.O.C)
wellgenetics.com

WELLGENETICS

**Genomic Sequencing**

**Sequences:**

>220862ex2_OWGb2734

TAGATACTTCGTGCTCATCTACGAGCTGCTGGATGAGATCCTCGACTTTGGCTACCCGCAGAACACGGACTCC
GGTACCCTGAAGACCTTCATCACACAGCAGGGCATCAAGTCGGCCACCAAGGAGGAGCAGATGCAGATAAC
CTCGCAGGTCACCGGCCAGATTGGCTGGCGTCGCGAGGGCATTAAGTACCGGCGGAACGAGCTTTTCCTGG
ACGTATTGGAGTACGTGAATCTGCTGATGAGCCCGCAGGGTCAGGTTTTGTCTGCCCACGTGGCCGGCAAGG
TGGTAATGAAGTCGTATTTGTCGGGTAAGTTAGCATAAATAATCTAGACATTATTCCTTTTAATAATCGCCATGTT
GTAGGCATGCCCGAGTGCAAGTTCGGCATTAACGACAAGATCGTGATGGAGTCTAAGGGACGCGGTCTCTCC
GGAAATTCAGAGGCGGAAACCTCACGCTCCGGCAAGCCCGTCGTGGTCATCGATGACTGCCAGTTCCATCAG
TGCGTCAAGCTAAGCAAATTCGAGACGGAGCATTCGATCAGCTTCATCCCGCCGGACGGGGAGTTCGAGCTG
ATGCGTTACCGTACCACCAAAGACATTTCGCTGCCATTCCGAGTCATCCCGCTGGTGCGGGAGGTGGGCCGCA
CCAGGGGAGAGAAGGTAAAA

+8862-2651-1809
13F.-12, No. 93, Sec. 1, Xintai 5th Rd., Xizhi Dist., New Taipei City 221416, Taiwan (R.O.C)
wellgenetics.com

**Blast Result:**

Sequence read (>220862ex2_OWGb2734, Sbjct) was aligned with 220862 excised donor sequence (Query) using BLAST. One endogenous TTAA motif (boxed in purple) was left and embedded in coding exon of *Ap-2mu/CG7057* after excision in *220862ex2*. Designed point mutation R168W of *Ap-2mu/CG7057* was boxed in orange.

220862ex2_OWGb2734 (Sbjct) vs 220862 excised donor (Query)

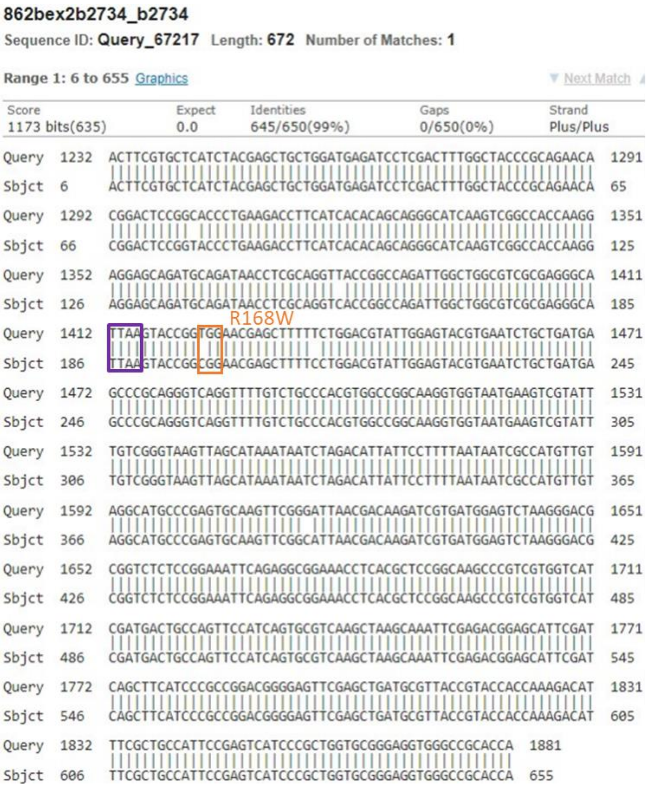

**862bex2b2734_b2734**

Sequence ID: **Query_67217** Length: **672** Number of Matches: **1**

Range 1: 6 to 655 Graphics ▼ Next Match

| Score | Expect | Identities | Gaps | Strand |
|---|---|---|---|---|
| 1173 bits(635) | 0.0 | 645/650(99%) | 0/650(0%) | Plus/Plus |

```
Query  1232  ACTTCGTGCTCATCTACGAGCTGCTGGATGAGATCCTCGACTTTGGCTACCCGCAGAACA  1291
Sbjct  6     ACTTCGTGCTCATCTACGAGCTGCTGGATGAGATCCTCGACTTTGGCTACCCGCAGAACA  65

Query  1292  CGGACTCCGGCACCCTGAAGACCTTCATCACACAGCAGGGCATCAAGTCGGCCACCAAGG  1351
Sbjct  66    CGGACTCCGGTACCCTGAAGACCTTCATCACACAGCAGGGCATCAAGTCGGCCACCAAGG  125

Query  1352  AGGAGCAGATGCAGATAACCTCGCAGGTTACCGGCCAGATTGGCTGGCGTCGCGAGGGCA  1411
Sbjct  126   AGGAGCAGATGCAGATAACCTCGCAGGTCACCGGCCAGATTGGCTGGCGTCGCGAGGGCA  185

                          R168W
Query  1412  TTAAGTACCGGTGGAACGAGCTTTTTCTGGACGTATTGGAGTACGTGAATCTGCTGATGA  1471
Sbjct  186   TTAAGTACCGGCGGAACGAGCTTTTCCTGGACGTATTGGAGTACGTGAATCTGCTGATGA  245

Query  1472  GCCCGCAGGGTCAGGTTTTGTCTGCCCACGTGGCCGGCAAGGTGGTAATGAAGTCGTATT  1531
Sbjct  246   GCCCGCAGGGTCAGGTTTTGTCTGCCCACGTGGCCGGCAAGGTGGTAATGAAGTCGTATT  305

Query  1532  TGTCGGGTAAGTTAGCATAAATAATCTAGACATTATTCCTTTTAATAATCGCCATGTTGT  1591
Sbjct  306   TGTCGGGTAAGTTAGCATAAATAATCTAGACATTATTCCTTTTAATAATCGCCATGTTGT  365

Query  1592  AGGCATGCCCGAGTGCAAGTTCGGGATTAACGACAAGATCGTGATGGAGTCTAAGGGACG  1651
Sbjct  366   AGGCATGCCCGAGTGCAAGTTCGGCATTAACGACAAGATCGTGATGGAGTCTAAGGGACG  425

Query  1652  CGGTCTCTCCGGAAATTCAGAGGCGGAAACCTCACGCTCCGGCAAGCCCGTCGTGGTCAT  1711
Sbjct  426   CGGTCTCTCCGGAAATTCAGAGGCGGAAACCTCACGCTCCGGCAAGCCCGTCGTGGTCAT  485

Query  1712  CGATGACTGCCAGTTCCATCAGTGCGTCAAGCTAAGCAAATTCGAGACGGAGCATTCGAT  1771
Sbjct  486   CGATGACTGCCAGTTCCATCAGTGCGTCAAGCTAAGCAAATTCGAGACGGAGCATTCGAT  545

Query  1772  CAGCTTCATCCCGCCGGACGGGGAGTTCGAGCTGATGCGTTACCGTACCACCAAAGACAT  1831
Sbjct  546   CAGCTTCATCCCGCCGGACGGGGAGTTCGAGCTGATGCGTTACCGTACCACCAAAGACAT  605

Query  1832  TTCGCTGCCATTCCGAGTCATCCCGCTGGTGCGGGAGGTGGGCCGCACCA  1881
Sbjct  606   TTCGCTGCCATTCCGAGTCATCCCGCTGGTGCGGGAGGTGGGCCGCACCA  655
```

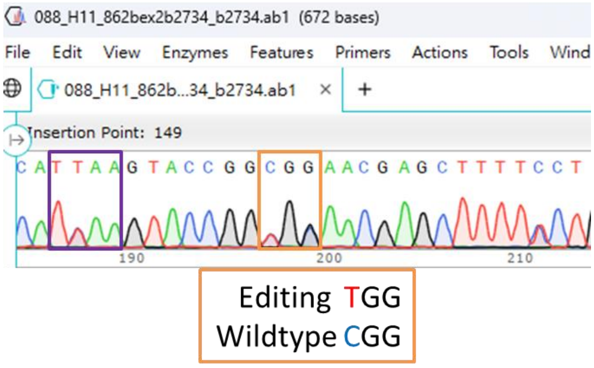

☎ +8862-2651-1809 ✉
13F.-12, No. 93, Sec. 1, Xintai 5th Rd., Xizhi Dist., New Taipei City 221416, Taiwan (R.O.C)
wellgenetics.com

WellGenetics

**Methods:**

PCR band of *220862ex2* was excised and submitted to Mission Biotech for gel extraction and sequencing. Sequence alignment shown here were using BLAST[1] (The Basic Local Alignment Search Tool) finds regions of local similarity between sequences.

[1] BLAST is a registered trademark of the National Library of Medicine.

+8862-2651-1809
13F.-12, No. 93, Sec. 1, Xintai 5th Rd., Xizhi Dist., New Taipei City 221416, Taiwan (R.O.C)
wellgenetics.com
